# Similarity in Early Life Stress Exposure is Associated with Similarity in Neural Representations in Early Adulthood

**DOI:** 10.1101/2024.12.02.626296

**Authors:** Miro Ilomäki, Jallu Lindblom, Marjo Flykt, Mervi Vänskä, Raija-Leena Punamäki, Patrik Wikman

## Abstract

Early life stress (ELS) has profound implications for developmental trajectories, yet the neural mechanisms underlying its long-term effects remain incompletely understood. In the present study, we examined whether interindividual similarity in ELS exposure aligns with similarity in neural representations and behavioral task performance in early adulthood. Leveraging a 20-year longitudinal dataset of Finnish families, we evaluated 87 young adults who underwent functional magnetic resonance imaging (fMRI) during an emotional go/no-go task. Intersubject representational similarity analysis (IS-RSA) was used to assess the associations between pairwise similarities in prospectively or retrospectively measured ELS, neural representations in 360 cortical regions, and task performance. We incorporated multidimensional scaling and Procrustes alignment to visualize interindividual differences in representational spaces. Prospective ELS, but not Retrospective ELS, was significantly associated with neural representational similarity across 40 cortical regions, including the anterior insula, frontal operculum, and anterior cingulate cortex. Higher Prospective ELS was also linked to reduced detection sensitivity, mediated by neural responses to angry facial expressions. These findings highlight the systematic and chronic effects of more moderate ELS on brain development and emphasize the value of prospective measurements and advanced similarity analyses in capturing the nuanced influences of ELS. By integrating spatial and shape analytical techniques, the present study provides new insights into the long-term neurobiological correlates of ELS and introduces novel methodological tools for neurodevelopmental research.

## INTRODUCTION

The complex relationship between early childhood experiences and brain development is a central topic in neuroscientific research. Much of the existing literature has focused on the impact of extreme forms of early life stress (ELS), such as abuse, neglect, and maltreatment, on various developmental outcomes. Research has extensively documented the long-term associations of extreme ELS on mental health (e.g., depression and anxiety; Baldwin et al., 2023; Gershon et al., 2013), emotion processing and self-regulation (e.g., learning, memory and emotion regulation; Wade et al., 2022; Pechtel & Pizzagalli, 2011; Mueller et al., 2010), structural brain changes (e.g., in amygdala and the medial prefrontal cortex; Ancelin et al, 2021; McLaughlin et al., 2019), and functional alterations (e.g., increased amygdala reactivity to negative emotional cues; Ross et al., 2021; Cohodes et al., 2020; Herzberg & Gunnar, 2020; Kraaijenvanger et al., 2020; Kaiser et al., 2018). However, the effects of more moderate but chronic ELS, such as strained family relationships and parental mental health problems, are less well understood. Despite being relatively common stressors, studying their long-term influences on brain functioning presents numerous challenges. One challenge is the common practice of categorizing participants into dichotomous “exposed” and “unexposed” groups. Such an approach may obscure the more nuanced and gradual effects of ELS on brain functioning. Additionally, many studies rely on retrospective self-report assessments of childhood adversities. While retrospective methods are efficient and often accurate, they are prone to various biases and limitations, such as childhood amnesia (Bell & Bell, 2018; Reuben et al., 2016; Lalande & Bonanno, 2011; Brennan et al., 2006; Hardt & Rutter, 2004). In contrast, longitudinal and prospective approaches, while costly and time consuming, mitigate memory-related biases and errors. As such, they provide a more reliable method to examine the effects of ELS on adult brain functioning.

An additional limitation in neurodevelopmental research is the frequent focus on univariate analyses of mean activity or functional connectivity of a limited number of brain regions. Advances in neuroscientific paradigms and computational analysis methods relating to understanding neural representations (Barack & Krakauer, 2021; Kriegeskorte & Kievit, 2013; Decharms & Zador, 2000) and individual nuances in brain function are still relatively rare in the neurodevelopmental literature, especially in ELS research. Notably, a large portion of neurodevelopmental research on structural and functional changes has focused on subcortical regions, especially on the amygdala and the hippocampus, as these regions have been strongly implicated in emotion and memory related processes (Phelps, 2004). Additionally, abundant research has focused on binodal functional connections between (limbic) subcortical regions and (largely frontal) cortical regions hypothesized to be involved in emotional experiences and self-regulation, such as the medial frontal gyrus, orbitofrontal cortex, the insula, and the cingulate cortex (Cohodes et al., 2020). While serving a necessary role, this univariate activation (or connectivity) analysis framework offers an incomplete picture (Freund et al., 2021). A gap thus exists in our understanding of how ELS might alter the development of larger-scale cortical representations of emotion processing and related behavior. Regardless, some neurodevelopmental studies utilizing novel computational neuroscientific methods have surfaced. For example, Saragosa-Harris et al. (2024) used representational similarity analysis (RSA; Kriegeskorte et al., 2008) to demonstrate that greater retrospectively evaluated ELS was associated with greater similarity in neural representations of ambiguous and threatening facial expression stimuli. Equivalent similarity approaches have been utilized in general neuroscientific studies of, for example, facial feature-related trustworthiness evaluation (e.g., Tashjian et al., 2019; FeldmanHall et al., 2018). By employing such multivariate methods that account for individual nuances in brain function, these approaches promise a shift in the neuroscientific paradigm and an elucidation of the complex outcomes of ELS that might have been missed by univariate approaches.

In the present study we leverage a unique, 20-year-spanning longitudinal dataset of Finnish families to examine the influence of ELS on task-based cortical representations in young adults. More specifically, we explore whether interindividual similarity in ELS (i.e., in terms of overall exposure) is associated with interindividual similarity in cortical neural representations of valence-related performance in an emotional go/no-go task that utilizes facial expression stimuli. The original go/no-go task was developed, and has been used, to study cognitive control (Gratton et al., 2018; Schulz et al., 2007). The emotional go/no-go is an adaptation of the task that utilizes emotional stimuli, such as emotional facial expressions, that interfere with cognitive control and is thus used to assess implicit emotion regulation (Ahmed et al., 2015). Importantly, previous research has reported ELS-related behavioral and neural alterations in facial emotion processing and recognition (Saarinen et al., 2021; Doretto & Scivoletto, 2018; da Silva Ferreira et al., 2014), emotion regulation (Miu et al., 2022; Schweizer et al., 2016), and inhibitory control (Wade et al., 2022; Brieant et al., 2023). However, results from various studies remain mixed or inconclusive. The general findings, however, suggest that those who have experienced adversities or chronic stress during childhood display altered behavioral and neural responses to tasks requiring facial emotion processing, emotion regulation, and cognitive control.

To elucidate this evasive area of inquiry, we utilize RSA in conjunction with intersubject representational similarity analysis (IS-RSA) to gauge whether interindividual similarity in ELS maps onto interindividual similarity in neural representation of the emotional go/no-go task. IS-RSA is a non-parametric computational analysis approach that essentially allows for an exploration of the association between interindividual, or pairwise, similarities in almost any variables of interest, and has been increasingly utilized during the current decade (e.g., Sheng et al., 2023; Hsiao et al., 2023; Ilomäki et al., 2022; Finn et al., 2020; Rhoads et al., 2020). For comparative purposes, we also employ more commonly used linear models and utilize both prospectively and retrospectively evaluated ELS. By leveraging these methods and the unique dataset, we aim to demonstrate how similarity in ELS is empirically mirrored in similarity in complex cortical neural representations of social-emotional information and related behavior even in early adulthood.

The present study aims to answer the following questions: (1) are individuals with similar Prospective or Retrospective ELS also more similar in terms of their cortical neural representations of the task, (2) is Prospective or Retrospective ELS associated with emotional suppression related neural activity, especially when inhibiting responses to social stimuli representing threat or negative valence (angry facial expressions), and (3) does Prospective or Retrospective ELS influence task performance in the emotional go/no-go task as measured by response sensitivity and reaction times. Our approach also attempts to highlight how different operationalizations of ELS might contribute to varying results and to assess the methodological viability and benefits of the similarity analysis approach.

## MATERIAL AND METHODS

### Participants

The current study is a segment of a Finnish longitudinal research project known as Miracles of Development (MIDE). In the project, 953 Finnish families have been monitored since pregnancy. Approximately half of these families conceived their child via successful assisted reproductive treatment (ART: *n* = 484 (51%); naturally conceived (NC): *n* = 469 (49%)). During pregnancy, the inclusion criteria for the parents included being Finnish-speaking, and additionally for the NC group, having no history of infertility and the mother’s age being over 25 years to match the higher age of ART mothers. See e.g., Vänskä et al. (2011) or Flykt et al. (2021), for a more detailed description of the larger study sample.

To ensure comprehensive representation of the study sample, disproportionate stratified sampling (Parsons, 2017) was used in the selection of the fMRI study participants. The sampling was based on the Prospective ELS index, which included 20 variables indicating maternal and paternal mental health and family relationship problems during the pregnancy (2^nd^ trimester) and when the child was 2 and 12 months old. See section “Prospective ELS operationalization” for a more detailed definition of the Prospective ELS index. The original MIDE sample was divided into four equal strata using this index (*z* < −0.42 for low; −0.42 ≤ *z* < 0.24 for moderate-low; 0.24 ≤ *z* < 0.90 for moderate-high; *z* ≥ 0.90 for high). We planned to select 24 participants from each of the four strata with a balanced selection for child’s sex and parents’ infertility history. Only cases with a maximum of 8 missing values in the ELS index were considered eligible. In total, 92 participants were successfully sampled, with some deviations from the original plan (e.g., some cells were depleted and replaced by nearby cells). The sample represented all strata (*χ^2^* (3) = 0.35, *p* = 0.951) and was balanced for child’s sex (*χ^2^* (3) = 1.55, *p* = 0.671) and parents’ fertility history (*χ^2^* (3) = 0.21, *p* = 0.976) within each stratum.

For the present neuroimaging substudy, both resting-state and task-based fMRI data were obtained from 92 young adults aged 18–21 years (*M* = 19.06, *SD* = 0.77; 55% female). Inclusion criteria for the fMRI study participation were right-handedness, being a native Finnish speaker, normal hearing, normal or corrected vision, and having no current psychiatric or neurological diagnoses. Due to excessive head motion (>0.2 mm mean framewise displacement) during the fMRI procedure, five participants were excluded from further analysis, leaving a final sample of 87 young adults for the current analyses. The fMRI experiment, along with the previous stages of the study, received approvals from the Ethics Committee of the Hospital District of Helsinki and Uusimaa, Finland.

### Prospective ELS operationalization

Prospective ELS was indexed by different domains, involving timing (during pregnancy (T1), when the child was 2 months old (T2), and when the child was 12 months old(T3)) through questionnaires regarding different problem domains (relationships and mental health symptoms) completed by both the mother and the father. Mental health problems in the family were measured using the General Health Questionnaire (GHQ36; mothers: α = 0.91–0.94, fathers: α = 0.92–0.94) (Goldberg & Hillier, 1979) and the Beck Depression Inventory (BDI-13; mothers: α = 0.75–0.84, fathers: α = 0.80–0.83) (Beck et al., 1961) at all three time points. GHQ assesses depression, anxiety, insomnia, and social dysfunction, whereas BDI assesses depression symptoms only.

Relationship problems within the family were evaluated using the Dyadic Adjustment Scale (DAS; mothers: α = 0.92–0.93, fathers: α = 0.91–0.91) (Spanier, 1976) and the Parenting Stress Index (PSI-36; mothers: α = 0.90–0.90, fathers: α = 0.91–0.91) (Abidin, 1997) when the child was 2 and 12 months old, but not during pregnancy. The DAS measures conflicts and low affection between parents, and the PSI assesses parenting distress and difficulties in parent-child relationships.

Complete Prospective ELS data, comprising 20 variables, were available for 84% (*n* = 77) of the participants, with 9% (*n* = 8) missing eight variables, and 7% (*n* = 7) missing one to four variables. Expectation-Maximization (EM) imputation, using data from the larger study sample, was employed to handle missing data. To create a total Prospective ELS score, a cumulative risk score (see Ettekal et al., 2019) was calculated by averaging the scores from each questionnaire over time and between parents, then standardizing and averaging these scores (*M* = 0.00, *SD* = 0.86, range = -1.5-2.34). The variables had moderately high internal consistency (α = 0.88).

### Retrospective ELS operationalization

Retrospective ELS was assessed using the self-report questionnaire items adapted from the Revised Adverse Childhood Experiences questionnaire (Finkelhor et al., 2015) approximately a year prior to fMRI data collection when the participants were 17–19 years old (*M* = 18.23, *SD* = 0.34).

Two items with a three-point Likert scale (0 = *never*, 1 = *sometimes*, 2 = *often*) assessed the following: Emotional abuse (e.g., “Did a parent or other adult in the household … swear at, insult, or put you down?”); Physical abuse (e.g., “Did a parent or other adult in the household … push, grab, shove, or slap you?”); Emotional neglect (e.g., “Did you … feel that no one in your family loved you or thought you were important or special?”); and Parent treated violently (e.g., “Was your parent … pushed, grabbed, slapped, or had something thrown at her/him?”). Two additional items were added to capture common ELS events for interparental psychological violence (e.g., “Have you seen your parent being threatened by violence at home?”; Ellonen et al., 2008).

Binary response items (0 = *no*, 2 = *yes*) assessed the following: Family alcohol and drug problems (“Did you live with anyone who was a problem drinker or alcoholic, or who used street drugs?”); Peer victimization (“Have you been bullied in school?”); Parents’ divorce (“Did your parents separate/divorce?”); Family mental illness (“Was a household member mentally ill?”); Death of a close person (“Have you ever lost anyone close to you by death?”); Family somatic illness (“Has any family member had a serious illness during your life?”); and Other serious adversities (“Have you experienced other adversities, such as accidents, victimization, or natural catastrophes?”). The total Retrospective ELS score was calculated by averaging the two-item domains and then summing the scores of the 12 domains (*M* = 4.45, *SD* = 2.60; range = 0-11.50).

### Task description and Stimuli

Participants completed an emotional go/no-go task (Hare et al., 2008) while undergoing fMRI (Figure 1). In this task, participants were instructed to press a button as fast as possible when a target emotional facial expression (neutral, happy, or angry; henceforth referred to as *valence*) appeared on the screen (‘go’ condition) and to withhold their response when a non-target facial expression appeared (‘no-go’ condition). A total of ten blocks were performed, each consisting of 48 trials: 32 target trials and 16 non-target trials. One of these blocks was a control task. Stimuli consisted of photographs of three facial expressions (happy, angry, neutral) of 16 different identities from the Umeå University Database of Facial Expressions (Samuelsson et al., 2012). Each stimulus was presented for 500 ms, followed by a jittered intertrial interval ranging from 2 to 14.5 seconds. Each task block comprised a specific pairing of two targets that were instructed before the start of each block, with blocks pseudorandomized across blocks (e.g., happy-angry, happy-neutral, or angry-neutral). Each task block was also accompanied with one of three types of feedback conditions (no feedback, positive feedback, negative feedback). In one-third of the blocks, points were awarded for correct trial responses (positive feedback). In one-third, points were deducted for incorrect responses (negative feedback). In one-third, no feedback was given. The feedback condition was instructed before the instruction for the target facial expressions (a dot for no feedback, “+” for positive feedback, and “-” for negative feedback). Feedback was given by presenting text on the screen, e.g., “+5”. Positive feedback was given on random trials with correct responses (on average 5 times over all correct responses within a positive feedback block). Positive feedback was an addition of five points to participant’s total score. However, approximately 1 out of every 5 positive feedback presentations was a larger number of points (“+30” points). Negative feedback was always a subtraction of five points and was always given after an incorrect response in the negative feedback blocks. To make the blocks with no feedback as visually similar as the blocks with feedback, random presentations of Xs with the same number of tokens as in the feedback conditions were presented in a similar manner. Whenever feedback was displayed in a trial, it was presented immediately after the response. The total score was visually represented using a semicircular gauge, akin to a car speedometer or compass. The display consisted of a half-circle, with an arrow that indicated the score by pointing to specific positions along the semicircle. The leftmost side of the half-circle represented the lowest possible score, and the arrow moved clockwise as the score increased, with the rightmost end representing the highest possible score. This allowed participants to quickly grasp their performance without relying on numerical values. There were short breaks between blocks, approximately 15 seconds, during which participants received instructions for the feedback condition and the targets for the next block. During this time, participants viewed a fixation cross before the next block of trials began. There was an additional control task block, during which participants were presented with the same facial expression stimuli under the same timing conditions but were instructed to respond to every stimulus. No feedback was given during the control block (analyses pertaining to this task were not included in the manuscript).

**Figure 1:**
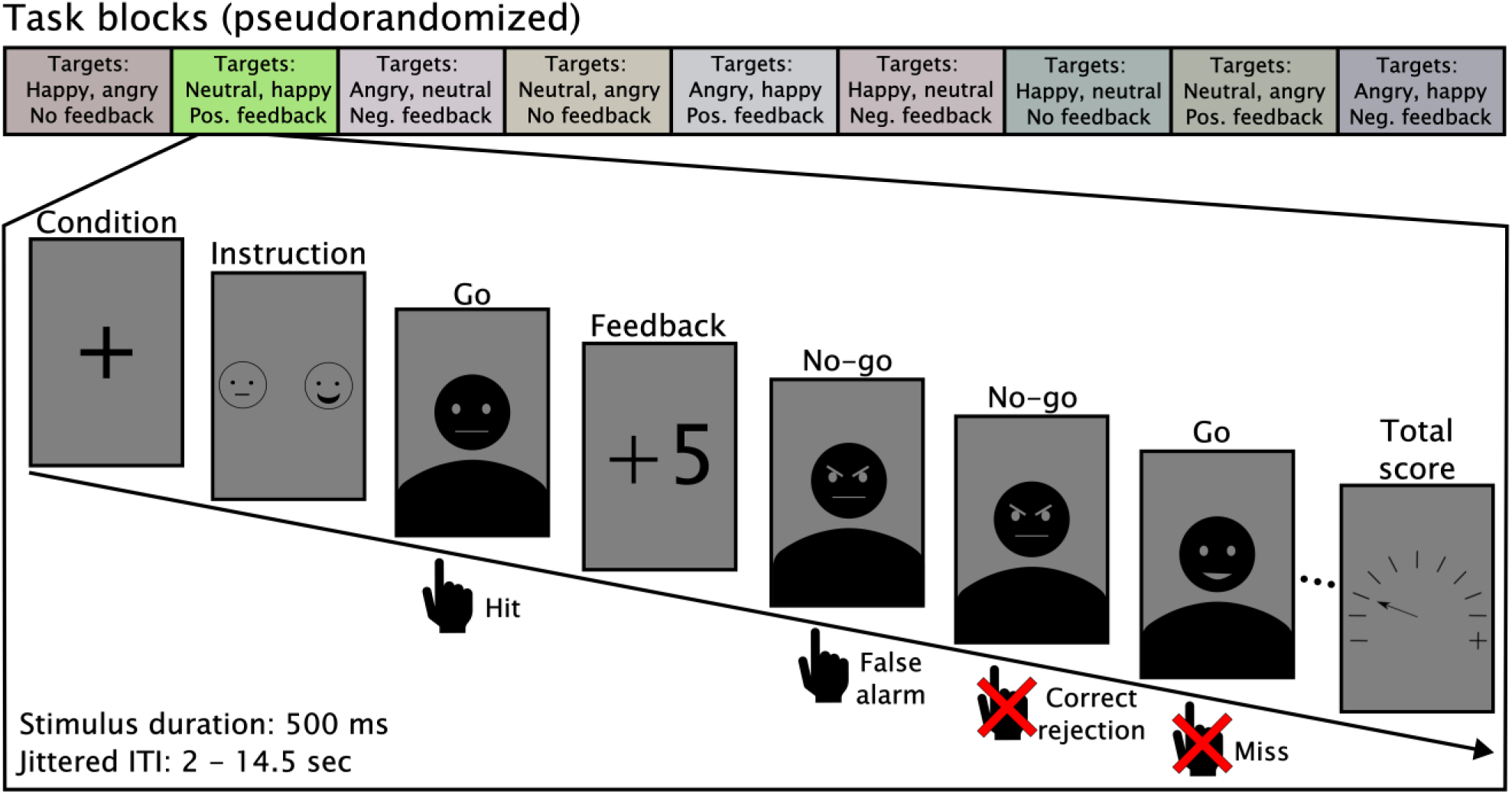
Schematic illustration of the emotional go/no-go task. Displayed on top are the nine variants of the task blocks (control task excluded). Blocks were pseudorandomized and varied based on the target stimuli and the feedback condition. A task block with neutral and happy emotional expression stimuli as targets and angry facial expressions as non-targets is highlighted as an example. Participants were instructed to press a button as fast as possible when a target facial expression appeared, and depending on the task block, points were given, taken, or not adjusted after responses. Each block began with instructions about the feedback condition and the target facial expressions. Participant responses were categorized according to signal detection theory: hits and correct rejections for correct commission and omission respectively, and false alarms and misses for commission and omission errors respectively. The total score was always displayed at the end of each block.

### Behavioral task performance

The primary dependent variables of the behavioral analyses were d-prime (*d’*), criterion (*c*), and reaction times (RT) to hits (responses to target stimuli). D-prime is a metric for signal detection sensitivity, while *c* is a metric for bias, or the tendency to favor one type of response over another. Higher values of *d’* indicate greater ability to distinguish target stimuli from non-target stimuli, while higher absolute values of c reflect a stronger tendency toward one type of response. To calculate *d’*, the *z*-transformed false alarm (FA) rate was subtracted from the *z*-transformed hit rate (HR): *d’* = *z(HR) - z(FA)*, while the criterion was calculated by taking the average of the z-transformed HR and FA and multiplying it by -1: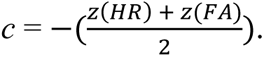

Linear mixed-effects models were employed to analyze the behavioral task performance metrics. RT, *d’* and *c* were modeled separately to investigate the effects of ELS and other covariates on task performance. Separate models were also run for Prospective and Retrospective ELS. ELS, valence (neutral, angry, happy), mother’s age, mother’s socioeconomic status (SES; indexed as mother’s education using 5-level grading from 0 = *university level education* to 4 = *no vocational education*), participant sex, and ART were included as fixed effects variable, and the subject identifier was specified as a random effect variable. Interaction effects were modeled for ELS × emotional expression type, ELS × participant sex, ELS × mother’s age, ELS × mother’s SES, and ELS × ART-status. Likelihood ratio tests (LRTs) were used to compare the fit of models with and without ELS predictors to determine the added explanatory power of ELS. F-statistics with Satterthwaite approximations were used to evaluate the significance of main effects and interaction terms within the linear mixed-effects framework. RT data were log-transformed to correct for skewness prior to analysis. Additionally, d’ was used in its respective similarity analyses (see “Similarity analyses” section). All analyses were conducted in R using the lme4 package for mixed-effects modeling.

### Fmri data

#### Acquisition

All data were acquired using a 3T MAGNETOM Skyra whole-body scanner (Siemens Healthcare, Erlangen, Germany) with a 20-channel head coil at the Advanced Magnetic Imaging (AMI) Centre, Aalto NeuroImaging, Aalto University School of Science, Espoo, Finland. During the imaging procedure, resting-state, structural, and task-based data were collected. The first task in the scanning session was the go/no-go task. The second task was a Reading the Mind in the Eyes task (RMET), involving photographs of faces cropped to show eyes only (adapted from Moor et al., 2012), where participants had to pick a word that either describes (1) the thoughts or feelings of the person in the photograph, or (2) the gender and age of the person in the photograph. Next, an anatomical scan [MPRAGE; high-resolution 3D T1 anatomical images (voxel matrix 256 × 256, in-plane resolution 1 mm × 1 mm × 1 mm)] was obtained. These anatomical images were used in the current study. Next, participants completed a social media task in which they posted opinions to a bogus Facebook group created by the experimenters and received peer feedback on those opinions (see Wikman et al., 2022). Finally, resting state data (participants were asked to lie still with eyes open) was acquired. All functional runs used echo-planar imaging (EPI) with an imaging area covering the whole brain comprising 43 contiguous oblique slices (TR 2,500 ms, TE 32 ms, flip angle 75◦, voxel matrix 64 × 64, field of view 20 cm, slice thickness 3.0 mm, in-plane resolution 3.1 mm × 3.1 mm × 3.0 mm). Participants were reimbursed 15 €/hour (2–3 h) for their time.

#### Preprocessing

We used fMRIPrep 20.2.5 (Esteban et al., 2018; Esteban et al., 2018) to preprocess the functional and structural MRI data. The anatomical T1-weighted (T1w) image was corrected for intensity non-uniformity (INU) with N4BiasFieldCorrection (Tustison et al., 2010), distributed with ANTs 2.3.3 (Avants et al., 2008) and skull-stripped with a Nipype implementation of the antsBrainExtraction.sh workflow (from ANTs), using OASIS30ANTs as target template. Brain tissue segmentation of cerebrospinal fluid (CSF), white-matter (WM) and gray-matter (GM) was performed on the brain-extracted T1w using fast (FSL 5.0.9, Zhang et al., 2001). Spatial normalization to standard space (MNI152NLin6Asym) was performed through nonlinear registration with antsRegistration (ANTs 2.3.3), using brain-extracted versions of both T1w reference and the T1w template. The following preprocessing was performed for the fMRI data: First, a reference volume and its skull-stripped version were generated using a custom methodology of fMRIPrep. A deformation field to correct for susceptibility distortions was estimated based on fMRIPrep’s fieldmap less approach. The deformation field is that resulting from co-registering the BOLD reference to the same-participant T1w-reference with its intensity inverted (Huntenburg, 2014; Wang et al., 2017). Registration is performed with antsRegistration (ANTs 2.3.3), and the process regularized by constraining deformation to be nonzero only along the phase-encoding direction and modulated with an average fieldmap template (Treiber et al., 2016). Based on the estimated susceptibility distortion, a corrected EPI reference was calculated for a more accurate co-registration with the anatomical reference. The BOLD reference was then co-registered to the T1w reference using bbregister (FreeSurfer) which implements boundary-based registration (Greve & Fischl, 2009). Co-registration was configured with six degrees of freedom. Head-motion parameters with respect to the BOLD reference (transformation matrices, and six corresponding rotation and translation parameters) are estimated before any spatiotemporal filtering using mcflirt (FSL 5.0.9, Jenkinson et al. 2002). BOLD runs were slice-time corrected using 3dTshift from AFNI 20160207 (Cox & Hyde, 1997) and resampled onto their original, native space by applying a single, composite transform to correct for head-motion and susceptibility distortions. The BOLD time-series were then resampled into standard space (MNI152NLin6Asym). Confounding time-series for framewise displacement (FD), DVARS and three region-wise global signals were calculated based on the preprocessed BOLD. FD was computed using two formulations following Power (absolute sum of relative motions, Power et al., 2014) and Jenkinson (relative root mean square displacement between affines, Jenkinson et al., 2002). FD and DVARS are calculated for each functional run, both using their implementations in Nipype (following the definitions by Power et al., 2014). The three global signals are extracted within the cerebrospinal fluid (CSF), the white matter (WM), and the whole-brain masks (global signal, GS). Additionally, a set of physiological regressors were extracted to 1 allow for component-based noise correction (aCompCor, Behzadi et al., 2007). Gridded (volumetric) resamplings were performed in a single interpolation step using antsApplyTransforms (ANTs), configured with Lanczos interpolation to minimize the smoothing effects of other kernels (Lanczos, 1964).

#### First level model

We used FEAT (FMRI Expert Analysis Tool, version 6.00), part of FSL (FMRIB’s Software Library, www.fmrib.ox.ac.uk/fsl), to perform two separate first level general linear model (GLM) analyses. Altogether possible estimatable variables comprised combinations of factors Valence (neutral, happy, angry), Correctness (correct, incorrect), Motor response (response, no-response), and Feedback (no-feedback, positive feedback, negative feedback). However, because these conditions depended on the behavior of the participant, it was impossible to estimate all the combinations of the conditions for each participant. Therefore, we opted to run two separate GLMs: the first model included factors Valence (neutral, happy, angry), Correctness (correct, incorrect), and Motor response (response, no-response), henceforth referred to as *performance model* (12 regressors of interest); in the other we combined the factors Correctness and Motor response into the factor Target (target, non-target) and thus, this model included the factors Target (target, non-target), Valence (neutral, happy, angry), and Feedback (no-feedback, positive feedback, negative feedback), henceforth referred to as *reward model* (9 regressors of interest). Note that the reward model was only used in this manuscript to define regions of interest for a specific similarity analysis. In both models we added regressors for all the control tasks, the feedback presentation, and the presentation of the total score (which were not analyzed in this manuscript). In all GLMs regressors were included in FLS’s three column format, including their temporal derivates (start time in seconds, trial duration one second). To account for motion, drift, and other nuisance factors, the model included the following regressors calculated by *fMRIPrep*: GS, the six basic motion, and the discrete cosine basis functions. A smoothing of 5 mm full width half maximum (FWHM) and whitening was applied in FEAT when running the model.

#### Second level analyses

For univariate within-subjects analysis, we ran a repeated measures ANOVA model across the whole functional volume using the Multivariate Repeated Measures (MRM) toolbox in Matlab (McFarquhar et al., 2016). The within-subjects factors for the model were Valence (neutral, happy, angry), Motor response (response vs. no response), and Correctness (correct response vs. incorrect response). We assessed all main effects and interactions of these factors.

To compare the similarity analysis results to results obtainable using between-subjects univariate methods, we ran two whole-brain ANCOVAs including the same within-subjects factors specified above and the between participants variables Prospective and Retrospective ELS (separately in their respective models), with mother’s age and SES, participant sex, and ART included as zero weighted covariates to isolate their influence.

In all second-level models we applied a cluster-based thresholding approach using permutation testing with 1000 iterations in each model to address multiple comparisons. We defined clusters at an uncorrected p-value threshold of 0.001. To control for Type I errors, we implemented a cluster-level family-wise error rate (FWER) correction with a threshold of 0.05.

#### Defining Regions of interest

Originally, we planned to define ROIs that were selective for faces with angry valence in the *performance model*. However, this model yielded few ROI candidates. Therefore, we defined our ROI clusters based on the *reward model*. Each significant cluster in the reward model was visually inspected using MRM post-estimations tools and if the ROI showed strongest activation during angry-valence trials it was extracted and converted to a binary mask. This yielded seven clusters included the following regions based on MRM’s automatic labeling: (1) left amygdala, (2) bihemispheric fusiform gyrus, (3) bihemispheric parahippocampal gyrus and left posterior cingulate cortex, (4) bihemispheric medial frontal gyrus, (6) left inferior and medial frontal gyrus, and (7) left anterior cingulate cortex, medial frontal gyrus, and uncus. This approach was grounded in previous research (Saragosa-Harris et al., 2024; Saarinen et al., 2021; Gollier-Briant et al., 2016; Pine et al., 2005) and our third research question regarding how ELS may influence the propensity for individuals to react differently to angry or threat signaling facial expressions.

#### Interindividual similarity analyses

We utilized IS-RSA to assess the associations between ELS, neural representations, and behavioral task performance. IS-RSA is a non-parametric method that examines relationships between variables by comparing interindividual similarities constructed across the sample. The analysis involves computing pairwise differences for each variable of interest, followed by comparing these differences across the variables. Specifically, we generated intersubject dissimilarity matrices (ISDMs) for each variable of interest, where each matrix represents pairwise comparisons between all unique pairs of participants, quantified using a distance metric tailored to the variable type. For single score aggregates or indices, such as the Prospective and Retrospective ELS scores, we employed Euclidean distance. For multidimensional vectors, such as neural representational profiles, we used correlation distance (calculated as 1 - r) to capture structural differences in the data rather than raw value differences.

To establish an effect size for the comparisons between the ISDMs, the following steps were taken: the redundant upper triangle of the ISDM was removed, alongside the diagonal (self-self-comparisons that are always maximally similar), resulting in all possible unique pairwise comparisons. In our sample, this resulted in 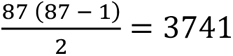unique pairwise comparisons for each ISDM, which were then vectorized. Finally, these vectorized ISDMs were compared using partial spearman correlation to establish the monotonic relationship between the rank orders of the pairwise comparisons between the compared matrices while controlling for confounders variables (mother’s age and SES, participant sex, and ART-status). See Figure 2 for a schematic illustration of the general analysis steps.

**Figure 2:**
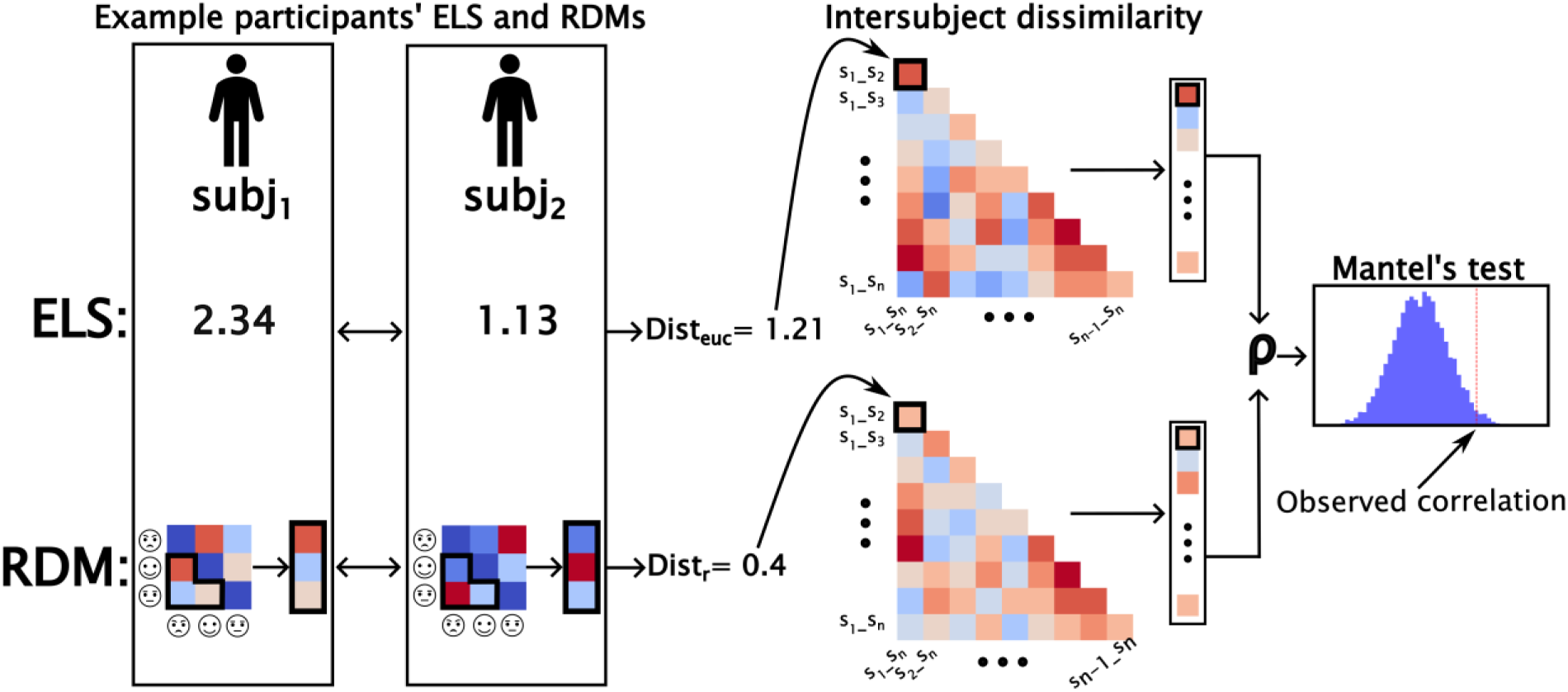
Schematic of the analysis steps for IS-RSA using the ELS ISDM × ROI ISDM analysis for one brain region as an example. Reading from left the right, first, each participant pair’s ELS and RDM are compared. The ELS is compared with simple Euclidean distance (dist_euc_), whereas the RDMs are compared by first vectorizing the non-redundant portion of the RDM, and then using correlation distance (dist_cor_) to attain a dissimilarity value for the pair. Finally, all pairwise dissimilarities are placed in their respective position in the intersubject dissimilarity matrix. The matrix comprises all the unique pairwise distances attainable from the current sample, spanning from comparing subj_1_ to subj_2_, to comparing subj_n-1_ to subj_n_. Then, the ELS and ROI intersubject dissimilarity matrices are vectorized, and a partial spearman correlation is calculated between them, with mother’s age and SES, participant sex, and ART-status partialed out. This attained correlation is finally tested by comparing it to the distribution of resampled correlations. These resampled correlations are attained by shuffling the rows and columns of one of the matrices, vectorizing it, recalculating the correlation, and repeating this process thousands of times. If the observed correlation deviates enough from the expected correlation based on the resampling, it will be deemed statistically significant.

For behavioral data, we used a measure of detection sensitivity (d’). Here, we calculated d’ separately for all the levels of the two within-subjects factors: valence and feedback condition. Even though the feedback conditions are not specifically analyzed in the present study, we included them in the calculations of participants’ d’ profiles to capture more performance related variance between individuals. After calculating the d’ profiles for each participant, we computed the d’ ISDM by calculating the correlation distance between each unique participant pair’s d’ profiles.

For neural representations of the task, we computed representational dissimilarity matrices (RDMs; Kriegeskorte et al., 2008) for every cortical Region of Interest (ROI) for each participant. ROIs were defined using the Human Connectome Project multimodal parcellation 1.0 (HCPMMP1.0, Glasser et al., 2016). The HCPMMP1.0 parcels the brain into 360 modal-specific regions with 180 ROIs per hemisphere. Each RDM comprised comparisons between a total of 12 conditions consisting of all the combinations of emotional expression type (neutral, happy, and angry) and signal detection theory derived response type (hit, correct rejection, false alarm, and miss). To compute the dissimilarities for the elements of the RDM for any given region, we calculated the correlation distances between all the unique pairs of the 12 conditions (e.g., correlation distance between happy-hit and angry-false alarm) using vectorized condition related mean signal changes of every voxel in the region. Finally, we computed the ISDMs separately for each region by calculating the correlation distance for each unique participant pairwise comparison using their vectorized RDMs after removal of the redundant upper triangle and the diagonal of the RDM.

Finally, we tested whether IS-RSA could be used to capture between-subjects variability from volumetric clusters with noticeable within-subjects variability in the repeated measures reward model. Here, we constructed an ISDM comprising correlation distances between participants’ vectors of mean signal changes during commission errors to angry facial expressions in the seven clusters extracted from the multivariate repeated measures results. More specifically, after running the model and inspecting the sources of within-subjects variability, we identified 7 volumetric clusters with the highest reactivity to angry facial expressions. These clusters were then used as ROIs by extracting their average percentual signal changes compared to the mean functional signal in moments where the participant committed a false alarm during an angry facial stimulus presentation (henceforth referred to as “angry FA”). Each participant thus received a profile of seven signal change percentage values, one for each ROI, which was then used in the creation of an ISDM by calculating a correlation distance between the angry FA “profiles” of each unique participant pair. This ISDM was then compared to Prospective and Retrospective ELS ISDM and d’ ISDM (see Figure 3 for a schematic illustration).

**Figure 3:**
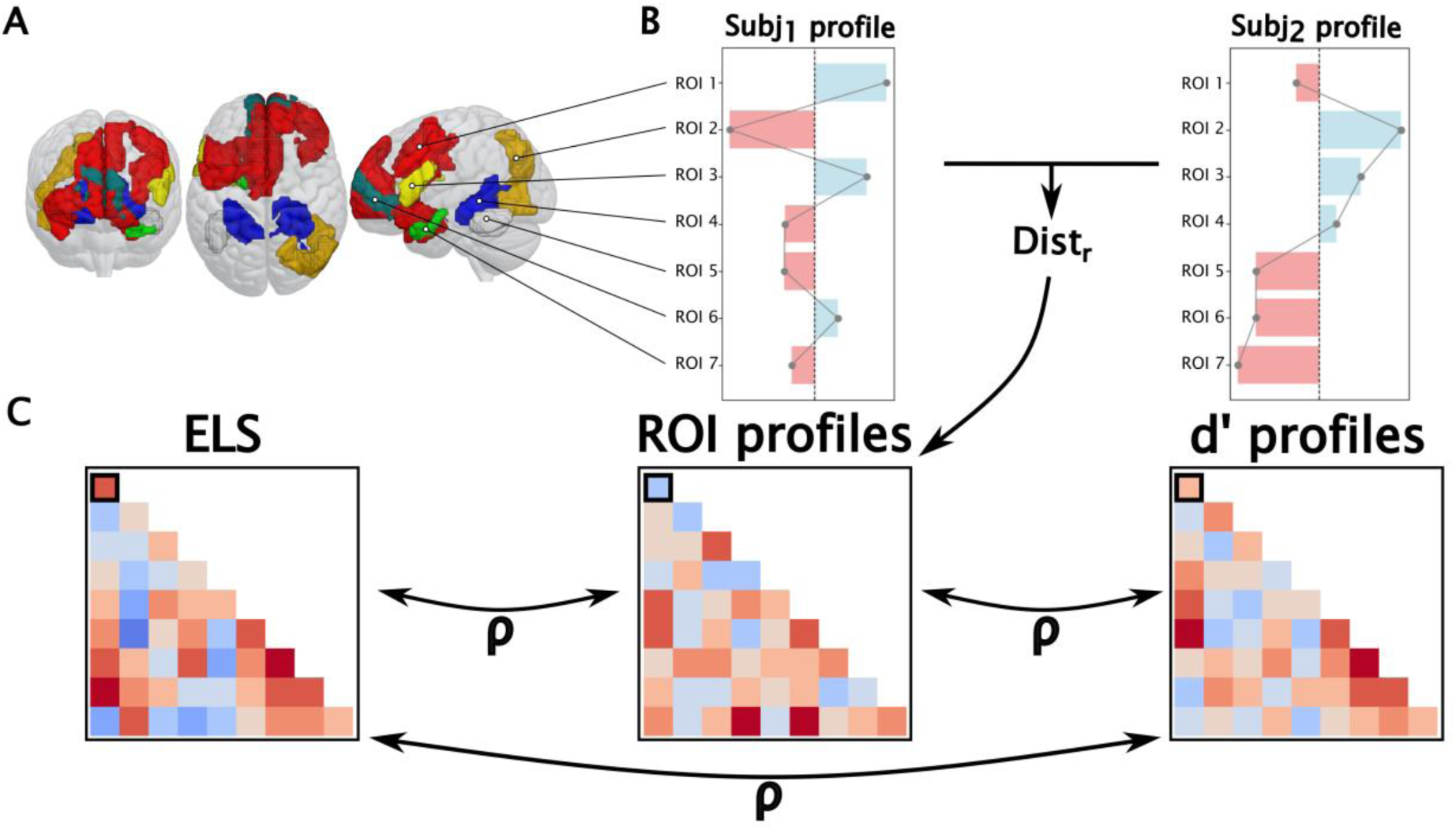
Schematic for the ELS ISDM × Angry FA ISDM analysis steps. (A) Shows the seven extracted volumetric regions (color coded) from the within-subjects linear model. (B) Displays an example profile of the average signal change during commission errors to angry facial expression presentations (compared to the mean functional signal) that was extracted for each cluster for each participant. (C) A pairwise distance between the profiles was then calculated using correlation distance and mapped onto an intersubject dissimilarity matrix, which was compared to ELS ISDM and d’ ISDM with Spearman’s correlation, both separately and in a mediation model, while controlling for mother’s age and SES, participant sex, and ART-status.

Due to the inflation of the amount of datapoints and the potential non-normality in the pairwise comparisons data, Mantel’s test was used to establish *p*-values for the ISDM correlations. In the Mantel’s test, the rows and columns of one of the compared matrices are shuffled and the correlation is recalculated. This procedure is then repeated thousands of times, and the observed correlation is compared to the distribution of correlations attained after reshuffling. For a positive one-tailed test, the p-value is then the amount of correlation values higher than the observed correlation divided by the number of reshufflings (or 1 – percentile of the observed correlation within the distribution of the permuted correlations). The Mantel’s test was performed for 360 ISDM correlations (once for each region from the HCPMMP 1.0 parcellation) for both Prospective and Retrospective ELS separately, each with 10 000 permutations. Here, False discovery rate (FDR) was used for correction of the family-wise error rate due to multiple comparisons.

Additionally, the following ISDM correlations were performed and tested with Mantel’s test: (1) Prospective ELS ISDM × d’ ISDM, (2) Retrospective ELS ISDM × d’ ISDM, (3) Prospective ELS ISDM × angry FA ISDM, and (4) Retrospective ELS ISDM × angry FA ISDM. Finally, using the Lavaan toolbox in Matlab, we tested whether the association between Prospective ELS ISDM × d’ ISDM was mediated by angry FA ISDM while controlling for the mother’s age and SES, participant sex, and ART-status. This was achieved with the Mantel’s test with 5000 iterations by shuffling the Prospective ELS and d’ ISDMs, recalculating the z-values with the vectorized ISDMs, and comparing the observed z-values to the resampled z-values.

#### Result inspection and visualizations

To explore the main result from the Prospective ELS ISDM × ROI ISDM analysis, individual domains of the ELS aggregate were separately tested in the same way as in the main analysis (but only for the significant regions from the main analysis) by constructing ISDMs for each ELS domains (responses during pregnancy (only BDI and GHQ), when the child was 2 months old, when the child was 12 months old, responses from the mother only, responses from the father only, mental health related questionnaires (BDI + GHQ) and family related problems questionnaires (RDAS + PSI)). Each domain was tested with the Mantel’s test with 5000 permutations and FDR correction at the .05 alpha level.

Due to IS-RSA being a simple correlation between two vectorized ISDMs, it is relatively uninformative about what underlies any observed correlation. Thus, after attaining the IS-RSA results, we utilized several novel approaches to verify and inspect the results. To this end, we utilized the correlations between the pairwise distance in each significant ROIs RDM element and the pairwise distance in ELS. This was achieved by subtracting each participant pair’s RDMs from each other, resulting in *difference RDMs* where each element of the matrix denotes the absolute difference between the values of that element for each pair of participants, and calculating Pearson’s correlation between the pair’s difference in ELS and the element-wise absolute differences of the RDMs. This resulted in a matrix (difference RDM) for each significant ROI from the Prospective ELS ISDM × ROI ISDM analysis that is informative about the direction and consistency of change in each element of the ROI specific RDM as a function of change in Prospective ELS. Next, we utilized both multidimensional scaling (MDS; Mead, 1992) and Procrustes alignment by applying 2-dimensional MDS on the RDMs for the ROIs with significant IS-RSA correlations, with ELS as the third dimension (Figure 4A). Procrustes alignment, first introduced by Schönemann (1966), is a low complexity approach to shape correspondence utilized in various scientific fields, including neuroscience (Guntupalli et al., 2016; Haxby et al., 2011). Because the MDS applied to the RDMs results in a variety of 2-dimensional spaces that cannot be directly compared due to their deviations in underlying spatial alignment, Procrustes alignment is utilized to find a common 2-dimensional space. Procrustes alignment takes in a set of points or vectors (also called the *reference*), and applies the optimal transform (scaling, rotations, and reflections) to a second matrix to minimize the sum of the squares of the pointwise differences between the reference and the second matrix. Here the participant with the lowest Prospective ELS was used as the reference. In addition to utilizing MDS and Procrustes alignment on the raw values of the RDMs, we used them to highlight how pairwise differences in Prospective ELS map onto pairwise differences in each RDM element. To achieve this, MDS was applied to the difference RDMs, which were then Procrustes aligned, and displayed against Prospective ELS dissimilarity (Figure 4B). This resulted in a 3-dimensional visualization that displays how the elementwise differences in the RDMs change as a function of increasing dissimilarity in Prospective ELS. Both Procrustes aligned MDS images were smoothed by fitting a polynomial curve (3- and 5-degree polynomial for the raw RDMs and the difference RDMs respectively) through the datapoints of each of the 12 elements in the RDM.

**Figure 4:**
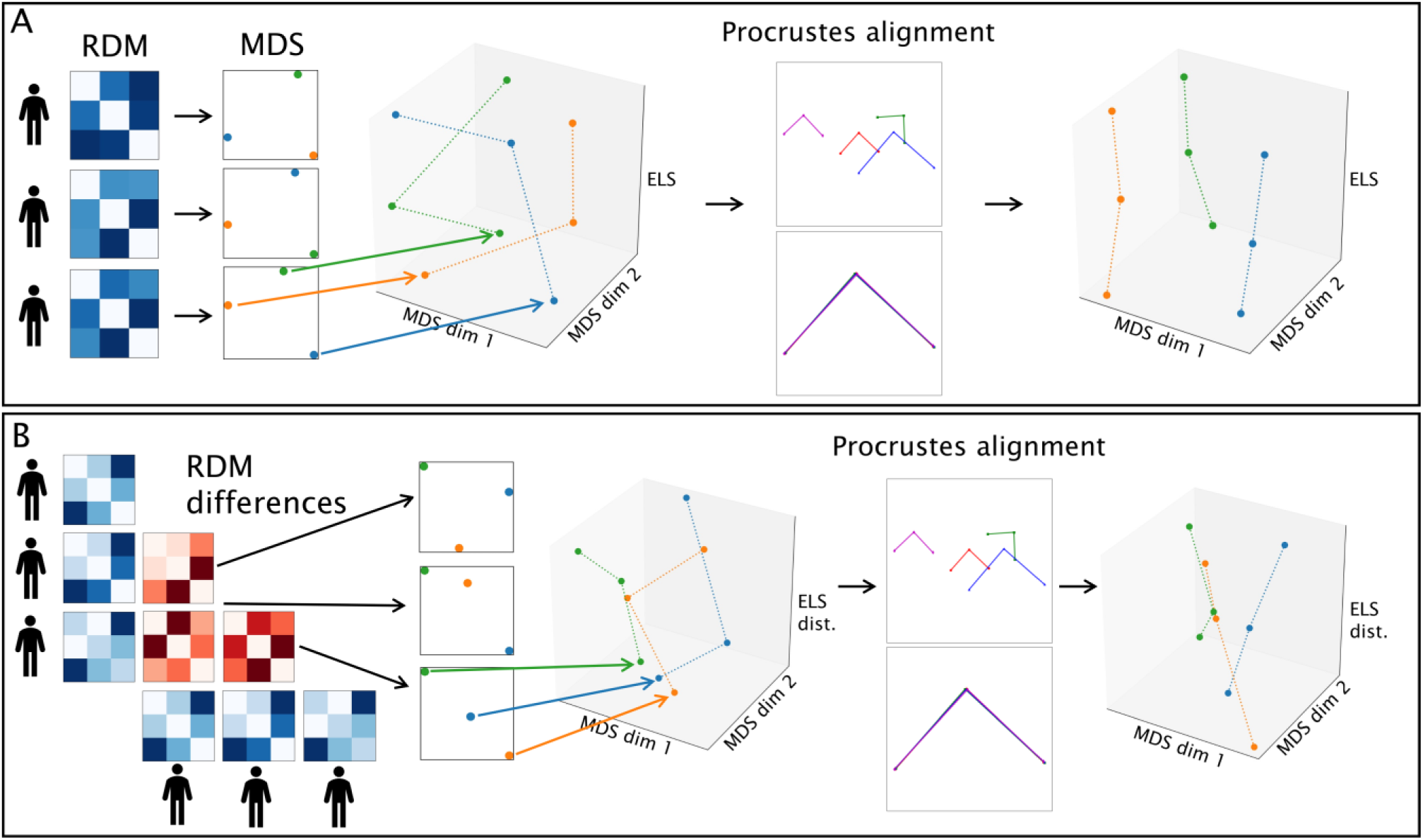
Schematic illustration of process for Procrustes aligned multidimensional scaling. First (A), the raw RDM values were translated into 2-dimensional multidimensional space. Because the multidimensional spaces are not properly aligned when plotting them against ELS, Procrustes alignment was used to align the spaces as closely as possible, using the MDS of the lowest ELS participant as the reference. To inspect how the pairwise differences in the RDMs changed as a function of the pairwise differences in ELS, (B) each participant’s region specific RDM was subtracted from every other region specific RDM in a pairwise manner, resulting in difference RDMs of absolute differences for each unique pairwise comparison for a given region. These difference RDMs were then translated into 2-dimensional multidimensional space. Finally, Procrustes alignment was used to align the multidimensional spaces using the pair with the highest ELS similarity as the reference.

Similarity analyses and visualizations were conducted with Python. The mediation analysis was conducted using the Lavaan toolbox (Rosseel, 2012) in R.

## RESULTS

### Interindividual similarity analysis results

#### Exploratory whole cortex IS-RSA results

We utilized IS-RSA to explore the correlations between pairwise differences in Prospective and Retrospective ELS and pairwise differences in neural representations within the 360 HCPMMP1.0 regions, while controlling for demographic covariates (see “Interindividual similarity analysis” section for details). Prospective and Retrospective ELS were both tested against the 360 regions separately. As presented in Figure 2, the correlations were attained by computing ISDMs for all the variables of interest using either Euclidean or correlation distance and calculating a partial Spearman’s correlation between the Prospective or Retrospective ELS ISDM and each of the ROI specific ISDMs, while controlling for mother’s age and SES, participant sex, and ART-status.

The 360 Prospective ELS ISDM × ROI ISDM tests revealed significant positive correlations for 40 cortical regions after FDR correction at the 0.05 alpha level, with significant correlations ranging from *r_s_* = .1 to *r_s_* = .24. Out of all the observed Prospective ELS ISDM × ROI ISDM correlations, 95.5% were positive, with an average correlation of *r_s_* = .071 (*r_s_*-min = -.045; *r_s_*-max = .245) and an average uncorrected p-value of .13 (*p*-min = .00001; *p*-max = .87). Notable significant regions seen only in the right hemisphere included frontal opercular regions, orbitofrontal cortex, lateral intraparietal cortex, inferior frontal gyrus, dorsolateral prefrontal cortex, and inferior regions around the precentral gyrus, including the prefrontal eye field. Notable significant regions only observed in the left hemisphere included the postcentral gyrus and ventromedial visual areas. Bihemispherically, significant positive correlations were observed in regions such as the anterior insula, anterior cingulate, and anterior regions around the middle and superior temporal gyrus. None of the Retrospective ELS ISDM × ROI ISDM correlations remained significant after FDR correction. Out of all the observed Retrospective ELS ISDM × ROI ISDM correlations, 63.9% were positive, with an average correlation of *r_s_* = .019 (*r_s_*-min = -.107; *r_s_*-max = .153) and an average uncorrected p-value of .378 (*p*-min = .003; *p*-max = 1).

Next, in order to explore which domains of the Prospective ELS aggregate may have driven the significant associations in the Prospective ELS ISDM × ROI ISDM IS-RSA, each significant ROI specific RDM from the Prospective ELS ISDM × ROI ISDM analysis was then correlated against the individual domains of the Prospective ELS aggregate. Each correlation between the domain of the aggregate and the RDMs of the significant regions were also tested using a Mantel’s test with 5000 permutations, and attained p-values were FDR corrected to avoid potentially spurious results. None of the ROIs had a significant correlation with parental problems reported during the pregnancy (T1) (*r_s_*-min = -.009; *r_s_*-max = .166; *r_s_*-mean = .079). Prospective ELS when the child was 2 months old (T2) was significantly associated with 13 regions (*r_s_*-min = .081; *r_s_*-max = .241; *r_s_*-mean = .149), and Prospective ELS when the child was 12 months old (T3) was associated with 16 regions (*r_s_*-min = .08; *r_s_*-max = .223; *r_s_*-mean = .147). Maternal self-reports were associated with 23 regions (*r_s_*-min = .1; *r_s_*-max = .288; *r_s_*-mean = .173), while paternal self-reports were associated with only 1 region (*r_s_*-min = -.01; *r_s_*-max = .164; *r_s_*-mean = .05). Prospective ELS related parental mental health related problems were associated with 2 regions (*r_s_*-min = .05; *r_s_*-max = .204; *r_s_*-mean = .114), while the family relationship problems were associated with 16 regions (*r_s_*-min = .061; *r_s_*-max = .22; *r_s_*-mean = .139). Figure 5 displays the 40 regions with significant Prospective ELS ISDM × ROI ISDM correlations, in addition to which of the regions had significant correlations with which individual domain of the Prospective ELS aggregate. See “supplementary_figure1” for a depiction of the sorted IS-RSA correlation values for the correlations between the different ELS indices and the 360 cortical ROI ISDMs.

**Figure 5:**
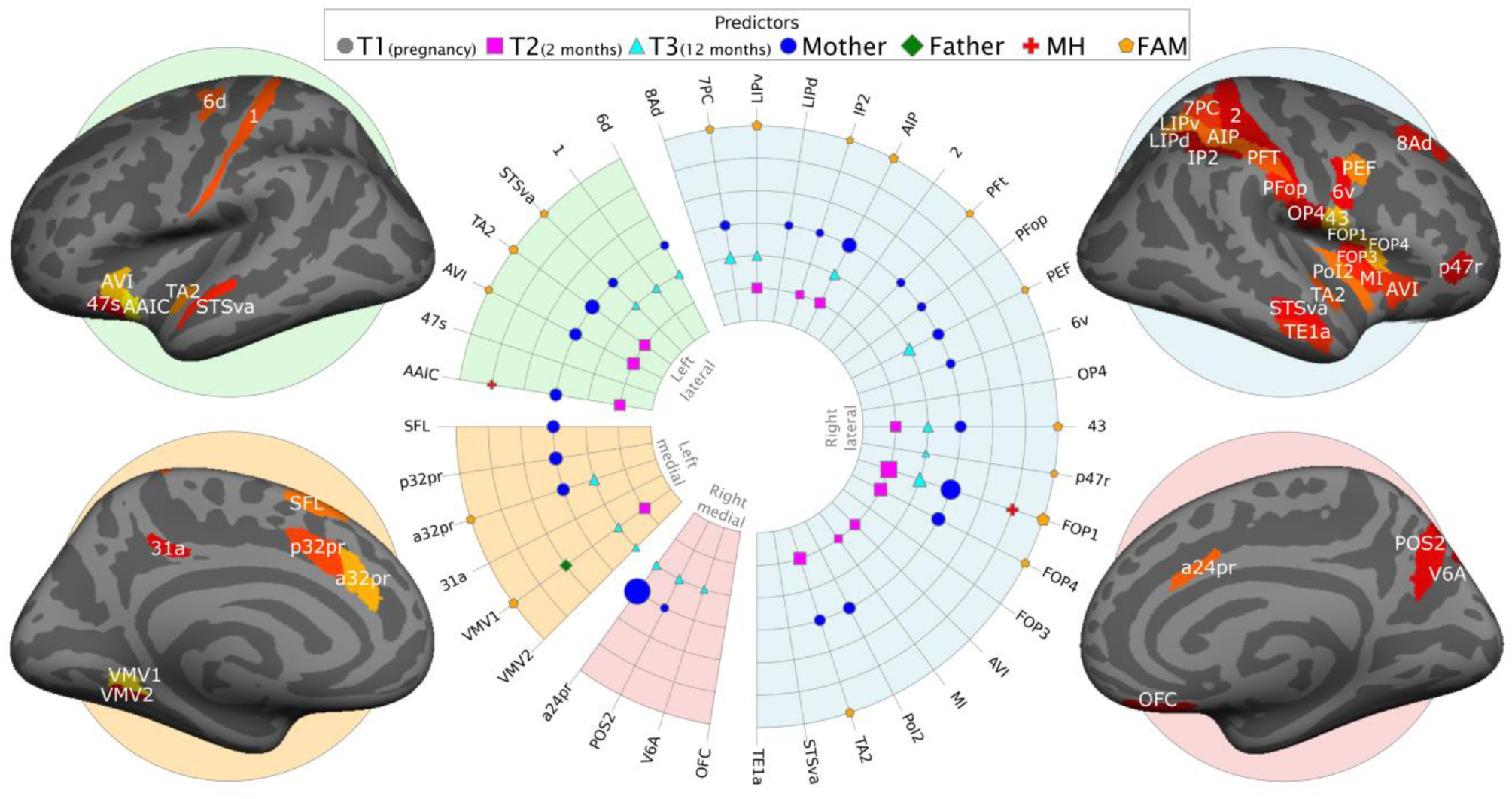
The 40 cortical regions with significant IS-RSA correlations from the Prospective ELS ISDM × ROI ISDM partial Spearman’s correlation highlighted on inflated medial and lateral surfaces for the right and left hemisphere. The radial plot displays which domains of Prospective ELS had significant IS-RSA correlations with the 40 regions. In the radial plot, the size of the marker indicates the size of the correlation, with larger markers denoting larger IS-RSA correlations. The predictors include the domains for timing of ELS (T1: pregnancy; T2: 2 months; T3: 12 months), reporter (mother or father self-reports), mental health problems (MH), and family relationship problems (FAM).

#### Other similarity analyses

Finally, as presented in Figure 3, we utilized the same similarity analysis approach to inspect whether there was a significant IS-RSA correlation between Prospective or Retrospective ELS ISDM, d’ ISDM, and the angry FA ISDM. The angry FA ISDM was constructed from pairwise differences in brain activity during commission errors to angry facial expressions in the seven volumetric clusters extracted from the reward model. Just like in the main analysis, partial Spearman’s correlation was used, with the same covariates controlled as in the main analysis. Out of these analyses, only the Prospective ELS ISDM × d’ ISDM had a significant positive IS-RSA correlation (*r_s_* = .101, *p* = .016). However, while the Prospective ELS ISDM × angry FA ISDM similarity analysis yielded only a small positive correlation that wasn’t statistically significant (*r_s_* = .038, *p* = .121), we explored whether the pairwise differences in the angry FA ISDM mediated the Prospective ELS ISDM × d’ ISDM association. Here, we utilized a combination of IS-RSA and a mediation model by comparing the observed mediatory effect to a distribution of resampled effects, while controlling for same covariates as in the previous analyses. To achieve the resampled effects, both the ELS and d’ ISDMs were shuffled like in the Mantel’s test, and the mediation effect was recalculated 5000 times with the shuffled matrices. A significant mediatory effect of angry FA on the association between Prospective ELS ISDM and d’ ISDM (*z*-direct = 5.029, *p* = .065; *z*-indirect = 5.829, *p* = .003; *z*-total = 6.821, *p* = .023) was observed in the mediation analysis.

#### Interindividual similarity analysis result visualizations and post-inspections

To better understand the Prospective ELS ISDM × ROI ISDM correlations, we utilized several visualizations and post-inspections for the data. First, due to the partial Spearman’s correlation used in the IS-RSA being a rank-based analysis, we rank transformed both the Prospective ELS ISDM and the ROI ISDMs for the 40 significant regions and plotted them against each other in a contour plots. These contour plots highlight how interindividual dissimilarity in each ROI’s RDM increases as a function of increasing interindividual Prospective ELS dissimilarity. Then, to visualize how the neural representations change as a function of Prospective ELS, we used multidimensional scaling and Procrustes alignment to plot the change in the raw values of the RDMs as a function of Prospective ELS (as demonstrated in Figure 4A). Additionally, in line with the pairwise approach utilized in the present study, we calculated difference RDMs where, for each unique pair of participants, the absolute difference in the elements of the pair’s RDMs was calculated. To visualize how the pairwise differences in the elements of the RDMs changed as a function of pairwise differences in Prospective ELS, these difference RDMs were then used in another Procrustes aligned multidimensional scaling plot (as demonstrated in Figure 4B). Finally, despite the IS-RSA results stemming from a multivariate approach, we calculated correlations between the pairwise differences in Prospective ELS and each difference RDM’s element. This approach explored a more linear direction and consistency of change in pairwise differences of the RDMs’ elements as a function of increasing pairwise dissimilarity in Prospective ELS. Figure 6 shows this approach using right frontal operculum region (FOP1), which had the strongest IS-RSA correlation with Prospective ELS. Figure 7 presents all the aforementioned post-inspection approaches for FOP1. For images of the other regions with significant Prospective ELS ISDM × ROI ISDM correlations, see supplementary figures 3-41.

**Figure 6:**
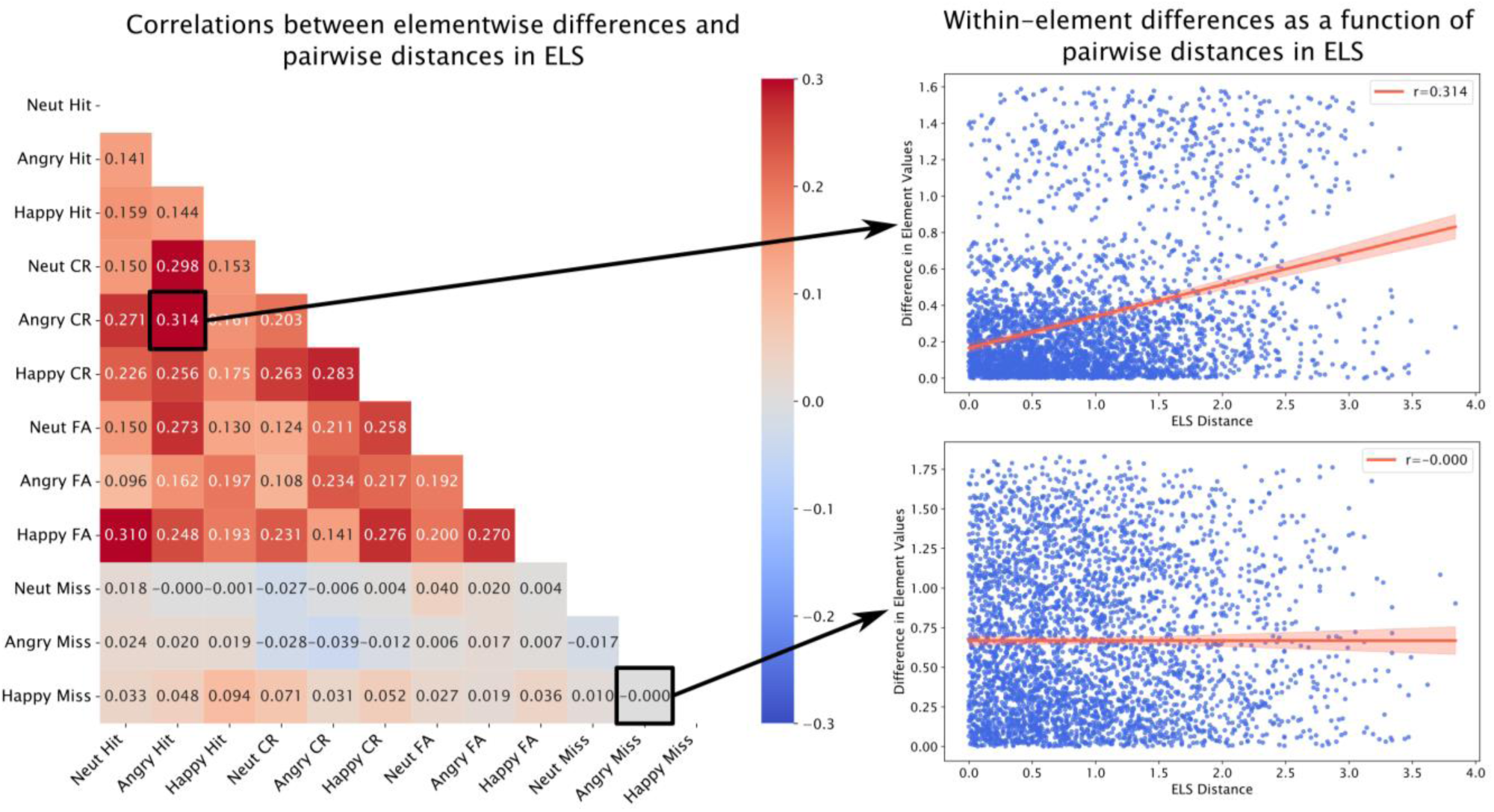
Change in the right frontal opercular region’s (FOP1) RDM’s elementwise differences as a function of ELS dissimilarity. The matrix (left) displays correlations between each element’s absolute differences and ELS dissimilarity, highlighting the direction and consistency of change in pairwise differences of an element as a function of increasing dissimilarity in Prospective ELS. To attain the matrix, the absolute difference between each unique participant pair’s RDM was first calculated, after which each element’s absolute difference was correlated with pairwise differences in Prospective ELS (right).

**Figure 7:**
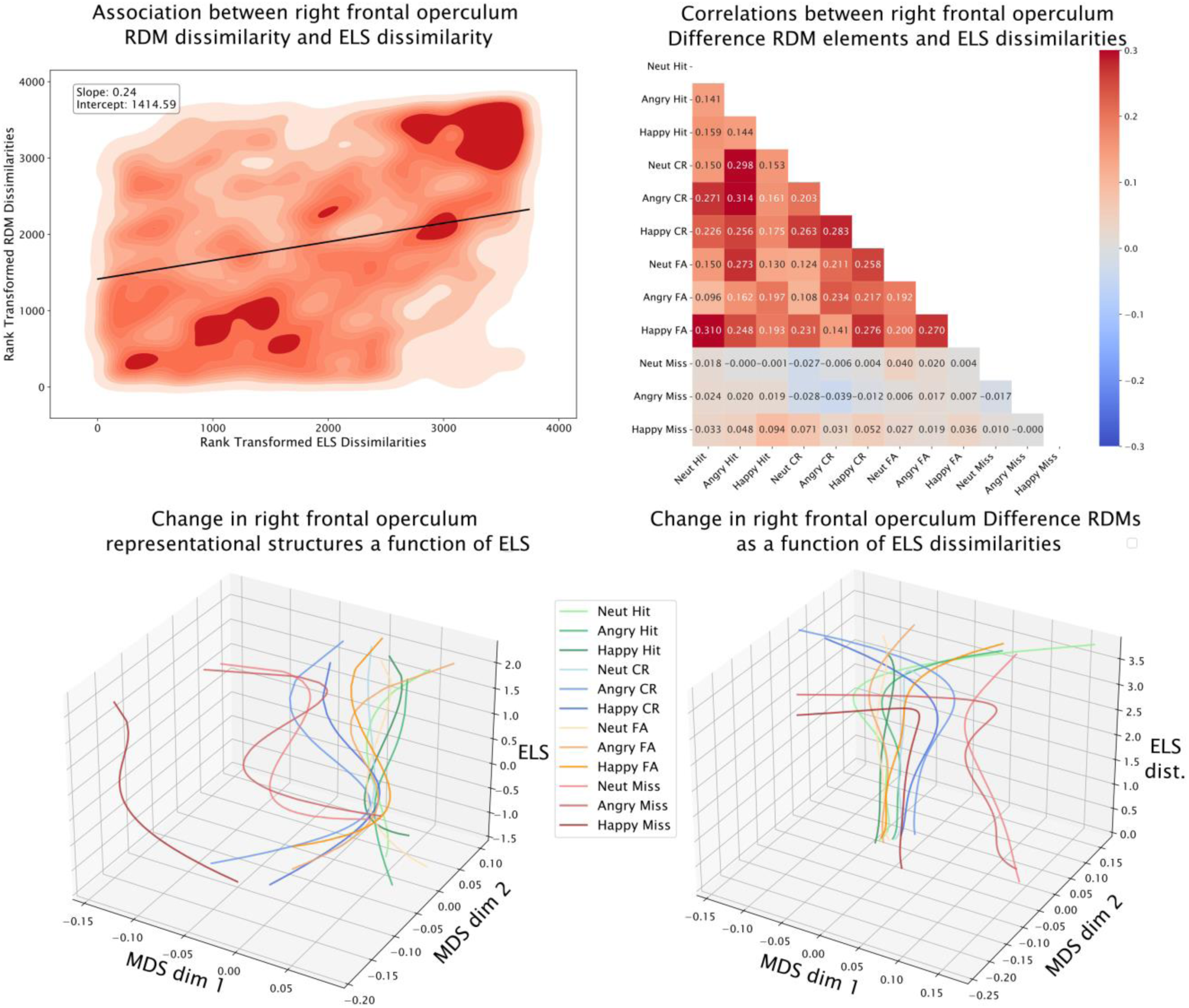
An example of the four visualizations computed for each region with significant Prospective ELS ISDM × ROI ISDM correlations. Here we use the frontal opercular region FOP1 of the HCPMMP1.0 as an example, as it had the highest IS-RSA correlation with Prospective ELS. The contour plot (top left) displays the association between the rank transformed pairwise distances of the FOP1 RDMs and the rank transformed Prospective ELS pairwise distances, highlighting their positive association. The heatmap (top right) displays Pearson’s correlations between pairwise distances in each RDM element and dissimilarity in ELS, highlighting the direction and consistency of change in each element as a function of Prospective ELS. The 3-dimensional MDS plot (bottom left) on the raw values of the FOP1 RDMs shows how the 2-dimensional representational structure, or space, extracted from the RDM changes as a function of increasing Prospective ELS. Here, a 3-degree polynomial curve was used for smoothing the lines. The 3-dimensional MDS plot (bottom right) on the difference RDM values of the FOP1 RDMs against Prospective ELS dissimilarities displays how the differences between each pair’s difference RDMs evolve as a function of Prospective ELS dissimilarity. The plot highlights how pairwise differences in ELS are associated with relatively stable absolute differences in most of the FOP1 RDMs’ elements, except for large dissimilarities in ELS which appear to result in large deviations in how the RDMs’ elements change as a function of ELS dissimilarity.

### Behavioral task performance results

#### D-prime and criterion

To investigate the association between ELS and participants’ d’ and c during the emotional go/no-go task, we first conducted a likelihood ratio test to compare models with and without Prospective or Retrospective ELS. Random intercepts accounted for individual differences in baseline performance. Compared to the null model, the results indicated a significant improvement in model fit when Prospective ELS and its interactions were included (LRT: *χ*^2^(10) = 24.65, *p* ≈ .006), but not when Retrospective ELS and its interactions were included (LRT: *χ*^2^(9) = 11.014, *p* ≈ .275). The Prospective ELS-included model using the Satterthwaite approximation revealed significant main effects (*F*(1,71.06) = 5.215, *p* = .025) on *d’*, where higher Prospective ELS was associated with lower d’ (β = −5.416). Prospective ELS and mother’s age had a statistically significant interaction (F(1,71.03) = 6.770, *p* = .011, β = .160), suggesting that higher Prospective ELS was not associated with as low a *d’* for those with older mothers compared to those with younger mothers.

#### Reaction time

We applied a similar model comparisons approach to reaction time data. Here, neither the model including Prospective ELS and its interactions (LRT: *χ*^2^(10) = 7.272, *p* ≈ .7), nor the model including Retrospective ELS and its interactions (LRT: *χ*^2^(9) = 10.668, *p* ≈ .3), improved the null model. In the null model, emotional facial expression type was a significant predictor of reaction time (*F*(2, 143.36) = 30.74, *p* < .001), with participants responding faster to angry faces (*M* = 474.105 ms) compared to happy (*M* = 493.926 ms) or neutral (*M* = 499.774 ms) faces. Additionally, there was a significant difference (*F*(2, 126.09) = 10.464, *p* = .002) between the reaction speeds of female participants (*M* = 460.634 ms) compared to male participants (*M* = 517.904 ms).

#### Whole brain repeated measures

The primary within-subjects repeated measures analysis revealed significant main effects for the factors of correctness (correct vs. incorrect responses), response type (response vs. nonresponse), and valence (neutral vs. happy vs. angry), indicating robust task-related neural activation across multiple brain regions (see “supplementary_figure42” for cortical surface projected effects). For correct vs. incorrect responses: Significant clusters were observed bilaterally in the postcentral gyrus and precentral gyrus, highlighting motor and sensory processing differences between correct and incorrect trials. Additional significant clusters for correct vs. incorrect were detected in the insula, cuneus, middle occipital gyrus, and superior temporal gyrus, among others. For response vs. nonresponse: Significant clusters were found in the bilateral inferior temporal gyrus, precuneus, postcentral gyrus, and fusiform gyrus. For valence: significant clusters were observed in the right inferior frontal gyrus, the left superior temporal gyrus, and the right middle frontal gyrus.

In the models incorporating between-subjects covariates—including either Prospective or Retrospective ELS, in addition to participant sex, mother’s age and SES, and ART history—no significant clusters were identified after FWE correction (*p* > .05).

## DISCUSSION

Uncovering the neurodevelopmental sequelae of early life stress (ELS) remains a challenging yet critical avenue of psychological and neuroscientific research. While numerous studies have documented associations between ELS and altered brain function, findings are yet to converge into clear, generalizable outcomes (Kraaijenvanger et al., 2020). This inconsistency likely reflects the complexity of the underlying neurodevelopmental processes, which may not be adequately captured by traditional univariate analyses that rely on mean activation differences or binary exposure categorizations. Such paradigms often inadvertently overlook the nuanced and gradual effects of chronic ELS.

To expand on previous research, we utilized a novel combination of representational similarity analysis (RSA) and its intersubject variant (IS-RSA) to investigate whether interindividual similarity in ELS (assessed both prospectively and retrospectively) exposure aligns with interindividual similarity in cortical neural representations during an emotional go/no-go task. Additionally, we explored whether Prospective and Retrospective ELS influence self-regulatory behavior and brain responses to negativity- or threat-related social stimuli (e.g., angry facial expressions). This study also sought to examine how operational definitions of ELS and advanced multivariate approaches contribute to variability in findings, thereby highlighting methodological considerations in neurodevelopmental research

The interindividual similarity analyses revealed significant positive correlations between interindividual similarity in Prospective ELS and neural representational similarity in 40 different cortical regions. However, none of the correlations between interindividual similarity in Retrospective ELS, assessed with a typical adverse childhood experiences (ACE) questionnaire, and cortical neural representations were significant. Most of the regions with significant correlations between similarities in Prospective ELS and neural representational similarity were in the lateral right hemisphere. Notable observed significant regions include the anterior insula, superior temporal sulcus, anterior cingulate cortex, and the right frontal operculum.

Many of the significant regions have been implicated in previous research on neurodevelopment, emotions, cognitive control, and facial emotion recognition. For example, the frontal operculum is related to facial emotion control (Caruana et al., 2016), processing (Bayer et al., 2018), and discrimination (Iarrobino et al., 2021). The insula, a laterally obscured cortical region beneath the sylvian fissure that is surrounded by the frontal, temporal, and parietal operculum, has been associated with a variety of different functions, such as interoception, pain processing, vestibular functions, attention, speech, and importantly, emotional experiencing and social cognition (Uddin et al., 2017). Bidirectional insular connectivity extends especially to regions such as the orbitofrontal, anterior cingulate, supplementary motor, primary and secondary somatosensory, and temporal cortices (Gasquoine, 2014). Many of these connected regions overlap with several regions with significant IS-RSA correlations in the current study. There is also brain region related agreement between the current study and evidence originating from research spanning from over four decades about the importance of the right hemisphere’s role in the recognition of emotional facial expressions (Ley & Bryden, 1979). A regional overlap can also be seen between the current study and a study by Adolphs et al. in 1996 that investigated cortical regions with the largest association with facial emotion recognition impairment in lesion patients. This overlapping of regions suggests both that the IS-RSA methodology captures functional alterations, and that Prospecive ELS, thus, likely influences how these specific regions represent facial emotional expressions.

Importantly, that interindividual similarity in Prospective ELS, but not Retrospective ELS, had significant positive correlations with neural representations highlights how moderate but chronic ELS might have a larger, or more systematic, influence on brain development than previously thought. Some previous research has explicitly focused on more moderate forms of ELS (Shapero et al., 2015; Edge et al., 2009), even though a consensus on the proper definition of moderate ELS is yet to emerge. This observation underscores the necessity of re-evaluating how moderate ELS is conceptualized and measured in both research and clinical settings. The lack of a clear consensus on its definition may contribute to variability in findings across studies, highlighting the need for standardized criteria that capture the nuanced, cumulative impacts of chronic but moderate ELS.

IS-RSA correlations between the 40 significant regions and the individual domains of the Prospective ELS index were also inspected. Parental mental health and relationship problems during the child’s infancy (when the child was 2 and 12 months old), problems reported by the mother in general (over all timepoints and questionnaire types), and family relationship problems in general (over both parents and all timepoints) had the highest IS-RSA correlations with the 40 regions. Surprisingly, interindividual similarity in ELS during the pregnancy (2^nd^ trimester) was not strongly associated with interindividual similarity in neural representations in any of the 40 regions, despite the contemporary scientific evidence strongly suggesting the importance of maternal prenatal distress (i.e., prenatal programming) for structural and functional brain development of the child (Wu et al., 2024). The discrepancy in findings may be caused confounding factors, such as (a) the nature of the emotional go/no-go task focusing on relatively narrow emotional-cognitive self-regulation, (b) the fact that few expecting mothers and fathers in the current study reported extremely high levels of distress symptoms, or (c) the inclusion of the father’s responses in the aggregate. Nevertheless, the results of the current study suggest that ELS relating especially to maternal and family related problems during the first year of life might play a large role in shaping cortical representations of facial emotion processing, emotion regulation, and cognitive control.

Finally, because emotion regulation problems and impulsive behavior have been implicated in those with experiences of ELS (Sanchez & Bangasser, 2022; McMullin et al., 2021), we investigate whether seven clusters displaying significantly different activity during presentations of angry facial expressions (extracted from a within-subjects model) would also display similar activation during commission errors to angry faces in those with similar ELS exposure. However, similarity in neither Prospective nor Retrospective ELS was significantly associated with similarity in activation of these regions. Nevertheless, because a significant association between similarities in Prospective ELS and detection sensitivity (d’) was observed, we additionally explored whether similarity in the activation profiles mediated this association. We found that similarity in the activation profiles did mediate the relationship between similarity in Prospective ELS and similarity in detection sensitivity. This finding suggests shared variance in interindividual similarities in Prospective ELS, brain activity, and behavioral responses, highlighting again the potential long-term influences that even more moderate but chronic forms of ELS may carry into early adulthood.

The most straightforward interpretation of these IS-RSA results is that individuals who are more similar in terms of Prospective ELS are also more similar in terms of how their brain represents socio-emotional information and related self-regulation. The results thus suggest that experiences during the first year of life related to parental mental health and family relationship problems may influence the development of a wide range of cortical regions in how they process socio-emotional information and coordinate cognitive-emotional self-regulation. However, due to the multidimensional nature of the analysis approach utilized in the present study, the aspects of the representational structure underlying the observed effects remain relatively obscured.

Nevertheless, the visualizations created using Procrustes aligned multidimensional scaling on the raw values of the RDMs (bottom left image in Figure 7; see also supplementary figures 3-41) demonstrate that there are some clear patterns in the neural representational spaces as a function of Prospective ELS. First, almost all the different conditions in the representational space seem to converge in those with moderate amounts of ELS, while those with low or high ELS seem to have a more varied neural representational structure. The exception to this pattern is another pattern, which is that omission errors are consistently dissimilar to most other conditions across the different regions, and they do not display consistent patterns as a function of ELS across different regions. The pairwise differences in the RDM elements for omission errors do not show any statistically clear relationship with pairwise differences in ELS, either (top right image in Figure 7; see also supplementary figures 3-41). This discrepancy in neural representations during omission errors might reflect attentional lapses during the task, which makes the lack of systematic interindividual variation in brain activity during these moments unsurprising. Another interesting pattern emerges in the multidimensional spaces of the pairwise differences in the RDMs as a function of pairwise differences in ELS across all significant regions (bottom right image in Figure 7; see also supplementary figures 3-41): Small pairwise differences in ELS seem to have a relatively consistent association with the pairwise differences in the representational structure, while large interindividual dissimilarities in ELS seem to relate to rapidly increasing variability in dissimilarities in neural representations.

Unlike the similarity analyses, the univariate whole-brain analyses did not yield significant ELS-related between-subjects differences. However, clear main effects for response correctness, motor response, valence of the stimulus, and the two-way interactions of correctness and valence with motor response were observed within-subjects. The difference in results between the whole-brain repeated measures approach and the IS-RSA approach suggests that univariate analyses might not be sensitive enough to capture complex differences between individual brains.

However, in linear models for the behavioral data, we observed that Prospective ELS was significantly associated with slightly lower detection sensitivity, suggesting that those with higher Prospective ELS were less sensitive to detecting and reacting accordingly to target and non-target facial expressions in the emotional go/no-go task. An interaction effect also emerged between Prospective ELS and the mother’s age, which suggested that this lowering of detection sensitivity due to ELS was not as extreme in those with older mothers. This interaction suggests that the potentially decremental effect of ELS on the ability to accurately recognize facial expressions and interpret social-emotional information might be attenuated in those with older mothers. As briefly mentioned earlier, a significant positive correlation was also observed between interindividual similarity in Prospective ELS and interindividual similarity in detection sensitivity using IS-RSA. No association between ACEs and detection sensitivity was observed in any of the models employed. Also, no differences in reaction time or criterion due to Prospective or Retrospective ELS were observed, suggesting that the cognitive faculties coordinating these behavioral aspects are not as strongly influenced by ELS as those coordinating detection sensitivity.

One of the narratives underlying the developmental alterations following ELS is that exposure to prolonged or extreme stress during early childhood results in biological cascades stemming from overt activation of the stress response system, disrupting the individual’s homeodynamic balance (Agorastos et al., 2019). Despite the generally grim narratives underlying the deleterious outcomes of ELS, these outcomes can also be viewed as expectable adaptations that help the individual survive in their early environment, shaping rather than overtly impairing cognition (Frankenhuis & de Weerth, 2013). In line with this, the IS-RSA methodology and the post-inspections utilized in the present study do not confer inferences about performance or health related normativity. Instead, they merely allow the inspection of within-sample similarity structures free from judgements about what should or should not be. That interindividual similarity in Prospective ELS was associated with interindividual similarity in neural representations does not inform us about whether a specific representational structure is inherently desirable. Instead, the result highlights a link between prospectively measured moderate-chronic ELS and brain function in an analytically robust way due to being less constrained by assumptions about the underlying nature of brain function. This approach is contrasted to frameworks such as the univariate activation analysis framework, with which no significant between-subjects differences could be uncovered in the current study. Thus, the current study demonstrates how a combination of RSA and IS-RSA, when paired with spatial and shape analytical approaches like multidimensional scaling and Procrustes alignment, offers legitimate aid in untangling the complex brain function related outcomes of ELS. Importantly, these approaches are not limited to developmental neuroscience and ELS research but can be extended to almost any variables of interest.

The findings of this study have broader implications for understanding the systematic influences of ELS on brain development without assigning normative judgments to specific neural or behavioral outcomes. The association between Prospective ELS and neural representational similarity demonstrates how chronic, moderate ELS exposure shapes brain function in a systematic and robust manner. The observed systematic interindividual differences in the neural representational structures may reflect adaptive responses to early environments and could therefore serve protective functions. This perspective leans on the notions of resilience and plasticity of the human brain, emphasizing its ability to organize itself in response to chronic stress inducing environments that pose unique demands for the child. The results also underscore the importance of considering moderate but chronic ELS domains, such as parental mental health and family relationship problems, in clinical settings. While extreme but transient ELS events remain central developmental disruptors, the results of the current study highlight the potential long-term developmental influence of chronic but moderate ELS.

Nevertheless, the need for more precise research undoubtedly remains to uncover how ELS influences brain development. For example, in the current study, family relationship problems were not evaluated during pregnancy due to the Prospective ELS questionnaires including questions about the parent-child relationship. While the sample was balanced for Prospective ELS due to the stratified sampling, few individuals displayed high ACEs in the Retrospective ELS measurement. This lack of high Retrospective ELS participants might at least partly explain why no significant results were obtained for it. Alternatively, the ACEs questionnaire concerns adverse events that have occurred also during middle and late childhood. The lack of systematic effects relating to Retrospective ELS in the current study might thus be due to (a) large within-sample variation in timing of adverse events over the whole childhood, (b) a lack of participants with adverse events during the early sensitive periods, or (c) a lack of variability in ACEs between participants. Future research would benefit from varied and rigorous operationalizations of ELS, perhaps by examining multiple aspects of distress (e.g. neglect vs. threat; McLaughlin & Sheridan, 2016) over various timepoints. Finally, as with many tasks utilized in psychological experiments, the emotional go/no-go task captures complex processes, some of which are likely yet to even be conceptualized. The structure employed for the representational dissimilarity matrices in the current study included both the quality of the emotional stimuli and the signal-detection theory-derived behavioral outcomes and is thus a complex outcome measure irreducible to singular conceptualizations like emotion regulation or cognitive control. Future research would benefit from disambiguation of neural representational structures (and ELS-related changes within them) by utilizing more simple tasks that reliably capture the underlying psychological construct of interest.

In conclusion, we utilized a combination of RSA and IS-RSA alongside spatial and shape analytical approaches to demonstrate complex associations between prospectively measured moderate ELS and relatively widespread cortical neural representations during an emotional go/no-go task in early adulthood. Spatial and shape analytical approaches revealed systematic patterns in neural representational spaces as a function of ELS, demonstrating their potential utility for future neuroscientific and -developmental research. The unique sample of Finnish families followed since pregnancyth enabled a bridging between the prenatal period and early adulthood of the participants, elucidating the far-reaching implications ELS may have on adult brain function. The utilization of different measurements of ELS informs future research about the use of various operationalizations and how they might influence observable outcomes. The difference in results between univariate activation analyses and IS-RSA demonstrated in the current study also highlights the need for researchers to adopt multivariate approaches that take into consideration the complexities of brain function.

## Data availability

Within-subjects fMRI result images, cluster masks, and analysis codes are available on the Open Science Framework (https://osf.io/j6bh2/). The raw data from the ongoing longitudinal study project are not readily available because participant privacy and ethical permissions do not allow public sharing of the data. Requests to access the datasets should be directed to R-LP.

## Author contributions

MI: Conceptualization, methodology, software, validation, formal analysis, data curation, writing – original draft, writing – review & editing, visualization, funding acquisition. JL: Conceptualization, investigation, data curation, writing – review & editing, supervision, project administration, funding acquisition. MF: investigation, data curation, writing – review & editing. MV: investigation, data curation, writing – review & editing. R-LP: conceptualization, investigation, resources, writing – review & editing, supervision, project administration, funding acquisition. PW: conceptualization, methodology, software, validation, investigation, data curation, writing – review & editing, supervision, project administration.

## Acknowledgements

The authors would like to thank Viljami Salmela and Tom Rosenström for methodological consultation, and Alessandra Bombino for assistance with the mediation analyses using R. This study was a part of the Miracles of Development research project supported by the Academy of Finland (R-LP, #3266413), and an individual grant from the Academy of Finland for JL (#323845). MI has received individual grants from the Finnish Brain Foundation, Signe and Ane Gyllenberg Foundation, and funding from the Doctoral programme in Human Behavior of the University of Helsinki.

## REFERENCES

Abidin, R. R. (1997). “Parenting stress index: A measure of the parent–child system,” in Evaluating Stress: A Book of Resources, eds C. P. Zalaquett and R. J. Wood (Scarecrow Education).

Adolphs, R., Damasio, H., Tranel, D., & Damasio, A. R. (1996). Cortical systems for the recognition of emotion in facial expressions. The Journal of neuroscience: the official journal of the Society for Neuroscience, 16(23), 7678–7687. 10.1523/JNEUROSCI.16-23-07678.1996

Agorastos, A., Pervanidou, P., Chrousos, G. P., & Baker, D. G. (2019). Developmental Trajectories of Early Life Stress and Trauma: A Narrative Review on Neurobiological Aspects Beyond Stress System Dysregulation. Frontiers in psychiatry, 10, 118. 10.3389/fpsyt.2019.00118

Ahmed, S. P., Bittencourt-Hewitt, A., & Sebastian, C. L. (2015). Neurocognitive bases of emotion regulation development in adolescence. Developmental cognitive neuroscience, 15, 11–25. 10.1016/j.dcn.2015.07.006

Ancelin, M. L., Carrière, I., Artero, S., Maller, J. J., Meslin, C., Dupuy, A. M., Ritchie, K., Ryan, J., & Chaudieu, I. (2021). Structural brain alterations in older adults exposed to early-life adversity. Psychoneuroendocrinology, 129, 105272. 10.1016/j.psyneuen.2021.105272

Avants, B.B., C.L. Epstein, M. Grossman, and J.C. Gee. (2008). “Symmetric Diffeomorphic Image Registration with Cross-Correlation: Evaluating Automated Labeling of Elderly and Neurodegenerative Brain.” Medical Image Analysis, 12(1), 26–41. 10.1016/j.media.2007.06.004

Baldwin, J. R., Wang, B., Karwatowska, L., Schoeler, T., Tsaligopoulou, A., Munafò, M. R., & Pingault, J. B. (2023). Childhood Maltreatment and Mental Health Problems: A Systematic Review and Meta-Analysis of Quasi-Experimental Studies. The American journal of psychiatry, 180(2), 117–126. 10.1176/appi.ajp.20220174

Barack, D. L., & Krakauer, J. W. (2021). Two views on the cognitive brain. Nature reviews. Neuroscience, 22(6), 359–371. 10.1038/s41583-021-00448-6

Bayer, M., Rubens, M. T., & Johnstone, T. (2018). Simultaneous EEG-fMRI reveals attention-dependent coupling of early face processing with a distributed cortical network. Biological psychology, 132, 133–142. 10.1016/j.biopsycho.2017.12.002

Beck, A. T., Ward, C. H., Mendelson, M., Mock, J., and Erbaugh, J. (1961). An inventory for measuring depression. Archives of general psychiatry, 4, 561–571. 10.1001/archpsyc.1961.01710120031004

Behzadi, Yashar, Khaled Restom, Joy Liau, and Thomas T. Liu. (2007). A Component Based Noise Correction Method (CompCor) for BOLD and Perfusion Based fMRI. NeuroImage, 37(1), 90–101. 10.1016/j.neuroimage.2007.04.042

Bell, D. C., & Bell, L. G. (2018). Accuracy of retrospective reports of family environment. Journal of child and family studies, 27, 1029–1040. 10.1007/s10826-017-0948-5

Buthmann, J. L., Miller, J. G., Uy, J. P., Coury, S. M., Jo, B., & Gotlib, I. H. (2024). Early life stress predicts trajectories of emotional problems and hippocampal volume in adolescence. European child & adolescent psychiatry, 33(7), 2331–2342. 10.1007/s00787-023-02331-4

Brennan, A. M., Stewart, H. A., Jamhour, N., Businelle, M. S., & Gouvier, Wm. D. (2006). An Examination of the Retrospective Recall of Psychological Distress. Journal of forensic neuropsychology, 4(4), 99–110. 10.1300/J151v04n04_06

Brieant, A., Clinchard, C., Deater-Deckard, K., Lee, J., King-Casas, B., & Kim-Spoon, J. (2023). Differential Associations of Adversity Profiles with Adolescent Cognitive Control and Psychopathology. Research on child and adolescent psychopathology, 51(12), 1725–1738. 10.1007/s10802-022-00972-8

Caruana, F., Gozzo, F., Pelliccia, V., Cossu, M., & Avanzini, P. (2016). Smile and laughter elicited by electrical stimulation of the frontal operculum. Neuropsychologia, 89, 364–370. 10.1016/j.neuropsychologia.2016.07.001

Cohodes, E. M., Kitt, E. R., Baskin-Sommers, A., and Gee, D. G. (2020). Influences of early-life stress on frontolimbic circuitry: Harnessing a dimensional approach to elucidate the effects of heterogeneity in stress exposure. Developmental psychobiology, 63(2), 157–172. 10.1002/dev.21969

Cox, R. W., & Hyde, J. S. (1997). Software tools for analysis and visualization of fMRI data. NMR in biomedicine, 10(4-5), 171–178. 10.1002/(sici)1099-1492(199706/08)10:4/5<171::aid-nbm453>3.0.co;2-l

Decharms, R. C., & Zador, A. (2000). Neural representation and the cortical code. Annual review of neuroscience, 23(1), 613–647. 10.1146/annurev.neuro.23.1.613

Doretto, V., & Scivoletto, S. (2018). Effects of early neglect experience on recognition and processing of facial expressions: A systematic review. Brain sciences, 8(1), 10. 10.3390/brainsci8010010

Edge, M. D., Ramel, W., Drabant, E. M., Kuo, J. R., Parker, K. J., & Gross, J. J. (2009). For better or worse? Stress inoculation effects for implicit but not explicit anxiety. Depression and anxiety, 26(9), 831–837. 10.1002/da.20592

Ellonen, N., Kääriäinen, J., Salmi, V., and Sariola, H. (2008). Lasten ja nuorten väkivaltakokemukset [Children’s and adolescents’ experiences of violence]. Oikeuspoliittinen Tutkimuslaitos [Finland’s Natl. Res. Instit. Legal Policy] 2008:152529.

Ettekal, I., Eiden, R. D., Nickerson, A. B., & Schuetze, P. (2019). Comparing alternative methods of measuring cumulative risk based on multiple risk indicators: Are there differential effects on children’s externalizing problems? PLoS one, 14(7), e0219134. 10.1371/journal.pone.0219134

Esteban, Oscar, Ross Blair, Christopher J. Markiewicz, Shoshana L. Berleant, Craig Moodie, Feilong Ma, Ayse Ilkay Isik, et al. 2018. “FMRIPrep.” Software. Zenodo. 10.5281/zenodo.852659

Esteban, Oscar, Christopher Markiewicz, Ross W Blair, Craig Moodie, Ayse Ilkay Isik, Asier Erramuzpe Aliaga, James Kent, et al. 2018. “fMRIPrep: A Robust Preprocessing Pipeline for Functional MRI.” Nature Methods. 10.1038/s41592-018-0235-4

da Silva Ferreira, G. C., Crippa, J. A., & de Lima Osório, F. (2014). Facial emotion processing and recognition among maltreated children: a systematic literature review. Frontiers in psychology, 5, 1460. 10.3389/fpsyg.2014.01460

FeldmanHall, O., Dunsmoor, J. E., Tompary, A., Hunter, L. E., Todorov, A., & Phelps, E. A. (2018). Stimulus generalization as a mechanism for learning to trust. Proceedings of the national academy of sciences of the United States of America, 115(7), E1690–E1697. 10.1073/pnas.1715227115

Finn, E. S., Glerean, E., Khojandi, A. Y., Nielson, D. M., Molfese, P. J., Handwerker, D. A., … & Bandettini, P. A. (2020). Idiosynchrony: from shared responses to individual differences during naturalistic neuroimaging. NeuroImage, 215, 116828. 10.1016/j.neuroimage.2020.116828

Finkelhor, D., Shattuck, A., Turner, H., & Hamby, S. (2015). A revised inventory of Adverse Childhood Experiences. Child abuse & neglect, 48, 13–21. 10.1016/j.chiabu.2015.07.011

Flykt, M., Vänskä, M., Punamäki, R.-L., Heikkilä, L., Tiitinen, A., Poikkeus, P., et al. (2021). Adolescent attachment profiles are associated with mental health and risk-taking behavior. Frontiers in Psychology, 12:761864. 10.3389/fpsyg.2021.761864

Frankenhuis, W. E., & de Weerth, C. (2013). Does Early-Life Exposure to Stress Shape or Impair Cognition? Current directions in psychological science, 22(5), 407–412. 10.1177/0963721413484324

Freund, M. C., Etzel, J. A., & Braver, T. S. (2021). Neural coding of cognitive control: the representational similarity analysis approach. Trends in cognitive sciences, 25(7), 622–638. 10.1016/j.tics.2021.03.011

Gasquoine P. G. (2014). Contributions of the insula to cognition and emotion. Neuropsychology review, 24(2), 77–87. 10.1007/s11065-014-9246-9

Gershon, A., Sudheimer, K., Tirouvanziam, R., Williams, L. M., & O’Hara, R. (2013). The long-term impact of early adversity on late-life psychiatric disorders. Current psychiatry reports, 15, 1–9. 10.1007/s11920-013-0352-9

Glasser, M. F., Coalson, T. S., Robinson, E. C., Hacker, C. D., Harwell, J., Yacoub, E., … & Van Essen, D. C. (2016). A multi-modal parcellation of human cerebral cortex. Nature, 536(7615), 171–178. 10.1038/nature18933

Goldberg, D. P., & Hillier, V. F. (1979). A scaled version of the General Health Questionnaire. Psychological medicine, 9(1), 139–145. 10.1017/s0033291700021644

Gollier-Briant, F., Paillere-Martinot, M. L., Lemaitre, H., Miranda, R., Vulser, H., Goodman, R., … & Artiges, E. (2016). Neural correlates of three types of negative life events during angry face processing in adolescents. Social cognitive and affective neuroscience, 11(12), 1961–1969. 10.1093/scan/nsw100

Gratton, G., Cooper, P., Fabiani, M., Carter, C. S., & Karayanidis, F. (2018). Dynamics of cognitive control: Theoretical bases, paradigms, and a view for the future. Psychophysiology, 55(3), e13016. 10.1111/psyp.13016

Greve, D. N., & Fischl, B. (2009). Accurate and robust brain image alignment using boundary-based registration. NeuroImage, 48(1), 63–72. 10.1016/j.neuroimage.2009.06.060

Guntupalli, J. S., Hanke, M., Halchenko, Y. O., Connolly, A. C., Ramadge, P. J., & Haxby, J. V. (2016). A Model of Representational Spaces in Human Cortex. Cerebral cortex, 26(6), 2919–2934. 10.1093/cercor/bhw068

Hare, T. A., Tottenham, N., Galvan, A., Voss, H. U., Glover, G. H., & Casey, B. J. (2008). Biological substrates of emotional reactivity and regulation in adolescence during an emotional go-nogo task. Biological psychiatry, 63(10), 927–934. 10.1016/j.biopsych.2008.03.015

Hardt, J., & Rutter, M. (2004). Validity of adult retrospective reports of adverse childhood experiences: review of the evidence. Journal of child psychology and psychiatry, 45(2), 260–273. 10.1111/j.1469-7610.2004.00218.x

Haxby, J. V., Guntupalli, J. S., Connolly, A. C., Halchenko, Y. O., Conroy, B. R., Gobbini, M. I., Hanke, M., & Ramadge, P. J. (2011). A common, high-dimensional model of the representational space in human ventral temporal cortex. Neuron, 72(2), 404–416. 10.1016/j.neuron.2011.08.026

Herzberg, M. P., & Gunnar, M. R. (2020). Early life stress and brain function: Activity and connectivity associated with processing emotion and reward. NeuroImage, 209, 116493. 10.1016/j.neuroimage.2019.116493

Hsiao, P. A., Kim, M. J., Chou, F. B., & Chen, P. A. (2024). Intersubject representational similarity analysis uncovers the impact of state anxiety on brain activation patterns in the human extrastriate cortex. Brain imaging and behavior, 18(2), 412–420. 10.1007/s11682-024-00854-1

Huntenburg, J. M. (2014). Evaluating Nonlinear Coregistration of BOLD EPI and T1w Images. Berlin: Master Thesis, Freie Universität.

Iarrobino, I., Bongiardina, A., Dal Monte, O., Sarasso, P., Ronga, I., Neppi-Modona, M., … & Ricci, R. (2021). Right and left inferior frontal opercula are involved in discriminating angry and sad facial expressions. Brain stimulation, 14(3), 607–615. 10.1016/j.brs.2021.03.014

Ilomäki, M., Lindblom, J., Salmela, V., Flykt, M., Vänskä, M., Salmi, J., Tolonen, T., Alho, K., Punamäki, R. L., & Wikman, P. (2022). Early life stress is associated with the default mode and fronto-limbic network connectivity among young adults. Frontiers in behavioral neuroscience, 16, 958580. 10.3389/fnbeh.2022.958580

Jenkinson, Mark, Peter Bannister, Michael Brady, and Stephen Smith. 2002. “Improved Optimization for the Robust and Accurate Linear Registration and Motion Correction of Brain Images.” NeuroImage 17 (2): 825–41. 10.1006/nimg.2002.1132

Kaiser, R. H., Clegg, R., Goer, F., Pechtel, P., Beltzer, M., Vitaliano, G., Olson, D. P., Teicher, M. H., & Pizzagalli, D. A. (2018). Childhood stress, grown-up brain networks: corticolimbic correlates of threat-related early life stress and adult stress response. Psychological medicine, 48(7), 1157–1166. 10.1017/S0033291717002628

Kraaijenvanger, E. J., Pollok, T. M., Monninger, M., Kaiser, A., Brandeis, D., Banaschewski, T., & Holz, N. E. (2020). Impact of early life adversities on human brain functioning: A coordinate-based meta-analysis. Neuroscience and biobehavioral reviews, 113, 62–76. 10.1016/j.neubiorev.2020.03.008

Kriegeskorte, N., Mur, M., & Bandettini, P. (2008). Representational similarity analysis - connecting the branches of systems neuroscience. Frontiers in systems neuroscience, 2, 4. 10.3389/neuro.06.004.2008

Kriegeskorte, N., & Kievit, R. A. (2013). Representational geometry: integrating cognition, computation, and the brain. Trends in cognitive sciences, 17(8), 401–412. 10.1016/j.tics.2013.06.007

Lalande, K. M., & Bonanno, G. A. (2011). Retrospective memory bias for the frequency of potentially traumatic events: A prospective study. Psychological trauma: theory, research, practice, and policy, 3(2), 165–170. 10.1037/a0020847

Lanczos, C. 1964. “Evaluation of Noisy Data.” Journal of the Society for industrial and applied mathematics series b numerical analysis, 1(1): 76–85. 10.1137/0701007

Ley, R. G., & Bryden, M. P. (1979). Hemispheric differences in processing emotions and faces. Brain and language, 7(1), 127–138. 10.1016/0093-934x(79)90010-5

McFarquhar, M., McKie, S., Emsley, R., Suckling, J., Elliott, R., & Williams, S. (2016). Multivariate and repeated measures (MRM): A new toolbox for dependent and multimodal group-level neuroimaging data. NeuroImage, 132, 373–389. 10.1016/j.neuroimage.2016.02.053

McLaughlin, K. A., & Sheridan, M. A. (2016). Beyond Cumulative Risk: A Dimensional Approach to Childhood Adversity. Current directions in psychological science, 25(4), 239–245. 10.1177/0963721416655883

McLaughlin, K. A., Weissman, D., & Bitrán, D. (2019). Childhood Adversity and Neural Development: A Systematic Review. Annual review of developmental psychology, 1, 277–312. 10.1146/annurev-devpsych-121318-084950

McMullin, S. D., Shields, G. S., Slavich, G. M., & Buchanan, T. W. (2021). Cumulative lifetime stress exposure predicts greater impulsivity and addictive behaviors. Journal of health psychology, 26(14), 2921– 2936. 10.1177/1359105320937055

Mead, A. (1992). Review of the development of multidimensional scaling methods. Journal of the Royal Statistical Society: Series D (The Statistician), 41(1), 27–39. 10.2307/2348634

Miu, A. C., Szentágotai-Tătar, A., Balazsi, R., Nechita, D., Bunea, I., & Pollak, S. D. (2022). Emotion regulation as mediator between childhood adversity and psychopathology: A meta-analysis. Clinical psychology review, 93, 102141. 10.1016/j.cpr.2022.102141

Mueller, S. C., Maheu, F. S., Dozier, M., Peloso, E., Mandell, D., Leibenluft, E., Pine, D. S., & Ernst, M. (2010). Early-life stress is associated with impairment in cognitive control in adolescence: an fMRI study. Neuropsychologia, 48(10), 3037–3044. 10.1016/j.neuropsychologia.2010.06.013

Parsons, V. L. (2017). “Stratified Sampling,” in Wiley statsref: Statistics reference online, eds N. Balakrishnan, T. Colton, B. Everitt, W. Piegorsch, F. Ruggeri, and J. L. Teugels (Hoboken, NJ: John Wiley & Sons). 10.1002/9781118445112.stat05999.pub2

Phelps E. A. (2004). Human emotion and memory: interactions of the amygdala and hippocampal complex. Current opinion in neurobiology, 14(2), 198–202. 10.1016/j.conb.2004.03.015

Pechtel, P., & Pizzagalli, D.A. (2011). Effects of early life stress on cognitive and affective function: an integrated review of human literature. Psychopharmacology 214, 55–70. 10.1007/s00213-010-2009-2

Pine, D. S., Mogg, K., Bradley, B. P., Montgomery, L., Monk, C. S., McClure, E., Guyer, A. E., Ernst, M., Charney, D. S., & Kaufman, J. (2005). Attention bias to threat in maltreated children: implications for vulnerability to stress-related psychopathology. The American journal of psychiatry, 162(2), 291–296. 10.1176/appi.ajp.162.2.291

Power, Jonathan D., Anish Mitra, Timothy O. Laumann, Abraham Z. Snyder, Bradley L. Schlaggar, and Steven E. Petersen. 2014. “Methods to Detect, Characterize, and Remove Motion Artifact in Resting State fMRI.” NeuroImage 84 (Supplement C): 320–41. 10.1016/j.neuroimage.2013.08.048

Reuben, A., Moffitt, T. E., Caspi, A., Belsky, D. W., Harrington, H., Schroeder, F., Hogan, S., Ramrakha, S., Poulton, R., & Danese, A. (2016). Lest we forget: comparing retrospective and prospective assessments of adverse childhood experiences in the prediction of adult health. Journal of child psychology and psychiatry, and allied disciplines, 57(10), 1103–1112. 10.1111/jcpp.12621

Rhoads, S. A., Cardinale, E. M., O’Connell, K., Palmer, A., VanMeter, J. W., & Marsh, A. A. (2020). Mapping neural activity patterns to contextualized fearful facial expressions onto callous-unemotional (cu) traits: intersubject representational similarity analysis reveals less variation among high-cu adolescents. Personality neuroscience, 3. 10.1017/pen.2020.13

Ross, M. C., Heilicher, M., & Cisler, J. M. (2021). Functional imaging correlates of childhood trauma: A qualitative review of past research and emerging trends. Pharmacology, biochemistry, and behavior, 211, 173297. 10.1016/j.pbb.2021.173297

Rosseel Y (2012). lavaan: An R Package for Structural Equation Modeling. Journal of Statistical Software, 48(2), 1–36. 10.18637/jss.v048.i02

Saarinen, A., Keltikangas-Järvinen, L., Jääskeläinen, E., Huhtaniska, S., Pudas, J., Tovar-Perdomo, S., Penttilä, M., Miettunen, J., & Lieslehto, J. (2021). Early Adversity and Emotion Processing From Faces: A Meta-analysis on Behavioral and Neurophysiological Responses. Biological psychiatry. Cognitive neuroscience and neuroimaging, 6(7), 692–705. 10.1016/j.bpsc.2021.01.002

Samuelsson, H., Jarnvik, K., Henningsson, H., Andersson, J., & Carlbring, P. (2012). The Umeå University Database of Facial Expressions: a validation study. Journal of medical Internet research, 14(5), e136. 10.2196/jmir.2196

Sanchez, E. O., & Bangasser, D. A. (2022). The effects of early life stress on impulsivity. Neuroscience and biobehavioral reviews, 137, 104638. 10.1016/j.neubiorev.2022.104638

Saragosa-Harris, N. M., Guassi Moreira, J. F., Waizman, Y., Sedykin, A., Peris, T. S., & Silvers, J. A. (2024). Early life adversity is associated with greater similarity in neural representations of ambiguous and threatening stimuli. Development and psychopathology, 1–13. 10.1017/S0954579424000683

Schulz, K. P., Fan, J., Magidina, O., Marks, D. J., Hahn, B., & Halperin, J. M. (2007). Does the emotional go/no-go task really measure behavioral inhibition?: Convergence with measures on a non-emotional analog. Archives of clinical neuropsychology, 22(2), 151–160. 10.1016/j.acn.2006.12.001

Schweizer, S., Walsh, N. D., Stretton, J., Dunn, V. J., Goodyer, I. M., & Dalgleish, T. (2016). Enhanced emotion regulation capacity and its neural substrates in those exposed to moderate childhood adversity. Social cognitive and affective neuroscience, 11(2), 272–281. 10.1093/scan/nsv109

Shapero, B. G., Hamilton, J. L., Stange, J. P., Liu, R. T., Abramson, L. Y., & Alloy, L. B. (2015). Moderate Childhood Stress Buffers Against Depressive Response to Proximal Stressors: A Multi-Wave Prospective Study of Early Adolescents. Journal of abnormal child psychology, 43(8), 1403–1413. 10.1007/s10802-015-0021-z

Sheng, J., Wang, S., Zhang, L., Liu, C., Shi, L., Zhou, Y., … & Xue, G. (2023). Intersubject similarity in neural representations underlies shared episodic memory content. Proceedings of the national academy of sciences, 120(35). 10.1073/pnas.2308951120

Schönemann, P.H. (1966). A generalized solution of the orthogonal Procrustes problem. Psychometrika, 31, 1–10. 10.1007/BF02289451

Spanier, G. B. (1976). Measuring dyadic adjustment: New scales for assessing the quality of marriage and similar dyads. Journal of marriage and the family, 38(1), 15–28. 10.2307/350547

Tashjian, S. M., Guassi Moreira, J. F., & Galván, A. (2019). Multivoxel Pattern Analysis Reveals a Neural Phenotype for Trust Bias in Adolescents. Journal of cognitive neuroscience, 31(11), 1726–1741. 10.1162/jocn_a_01448

Treiber, J. M., White, N. S., Steed, T. C., Bartsch, H., Holland, D., Farid, N., McDonald, C. R., Carter, B. S., Dale, A. M., & Chen, C. C. (2016). Characterization and Correction of Geometric Distortions in 814 Diffusion Weighted Images. PloS one, 11(3), e0152472. 10.1371/journal.pone.0152472

Tustison, N. J., Avants, B. B., Cook, P. A., Zheng, Y., Egan, A., Yushkevich, P. A., & Gee, J. C. (2010). N4ITK: improved N3 bias correction. IEEE transactions on medical imaging, 29(6), 1310–1320. 10.1109/TMI.2010.2046908

Uddin, L. Q., Nomi, J. S., Hébert-Seropian, B., Ghaziri, J., & Boucher, O. (2017). Structure and Function of the Human Insula. Journal of clinical neurophysiology: official publication of the American Electroencephalographic Society, 34(4), 300–306. 10.1097/WNP.0000000000000377

Vänskä, M., Punamäki, R.-L., Tolvanen, A., Lindblom, J., Flykt, M., Unkila-Kallio, L., Tiitinen, A., Repokari, L., Sinkkonen, J., & Tulppala, M. (2011). Maternal pre- and postnatal mental health trajectories and child mental health and development: Prospective study in a normative and formerly infertile sample. International Journal of behavioral development, 35(6), 517–531. 10.1177/0165025411417505

Wade, M., Wright, L., & Finegold, K. E. (2022). The effects of early life adversity on children’s mental health and cognitive functioning. Translational psychiatry, 12(1), 244. 10.1038/s41398-022-02001-0

Wang, S., Peterson, D. J., Gatenby, J. C., Li, W., Grabowski, T. J., & Madhyastha, T. M. (2017). Evaluation of Field Map and Nonlinear Registration Methods for Correction of Susceptibility Artifacts in Diffusion MRI. Frontiers in neuroinformatics, 11, 17. 10.3389/fninf.2017.00017

Wikman, P., Moisala, M., Ylinen, A., Lindblom, J., Leikas, S., Salmela-Aro, K., Lonka, K., Güroğlu, B., & Alho, K. (2022). Brain Responses to Peer Feedback in Social Media Are Modulated by Valence in Late Adolescence. Frontiers in behavioral neuroscience, 16, 790478. 10.3389/fnbeh.2022.790478

Wu, Y., De Asis-Cruz, J. & Limperopoulos, C. (2024). Brain structural and functional outcomes in the offspring of women experiencing psychological distress during pregnancy. Molecular Psychiatry, 29, 2223– 2240. 10.1038/s41380-024-02449-0

Zhang, Y., M. Brady, and S. Smith. (2001). “Segmentation of Brain MR Images Through a Hidden Markov Random Field Model and the Expectation-Maximization Algorithm.” IEEE Transactions on Medical Imaging 20 (1): 45–57. 10.1109/42.906424

