## Supplementary figures and images for "Similarity in Early Life Stress Exposure is Associated with Similarity in Neural Representations in Early Adulthood"

### Supplemental Figure 1

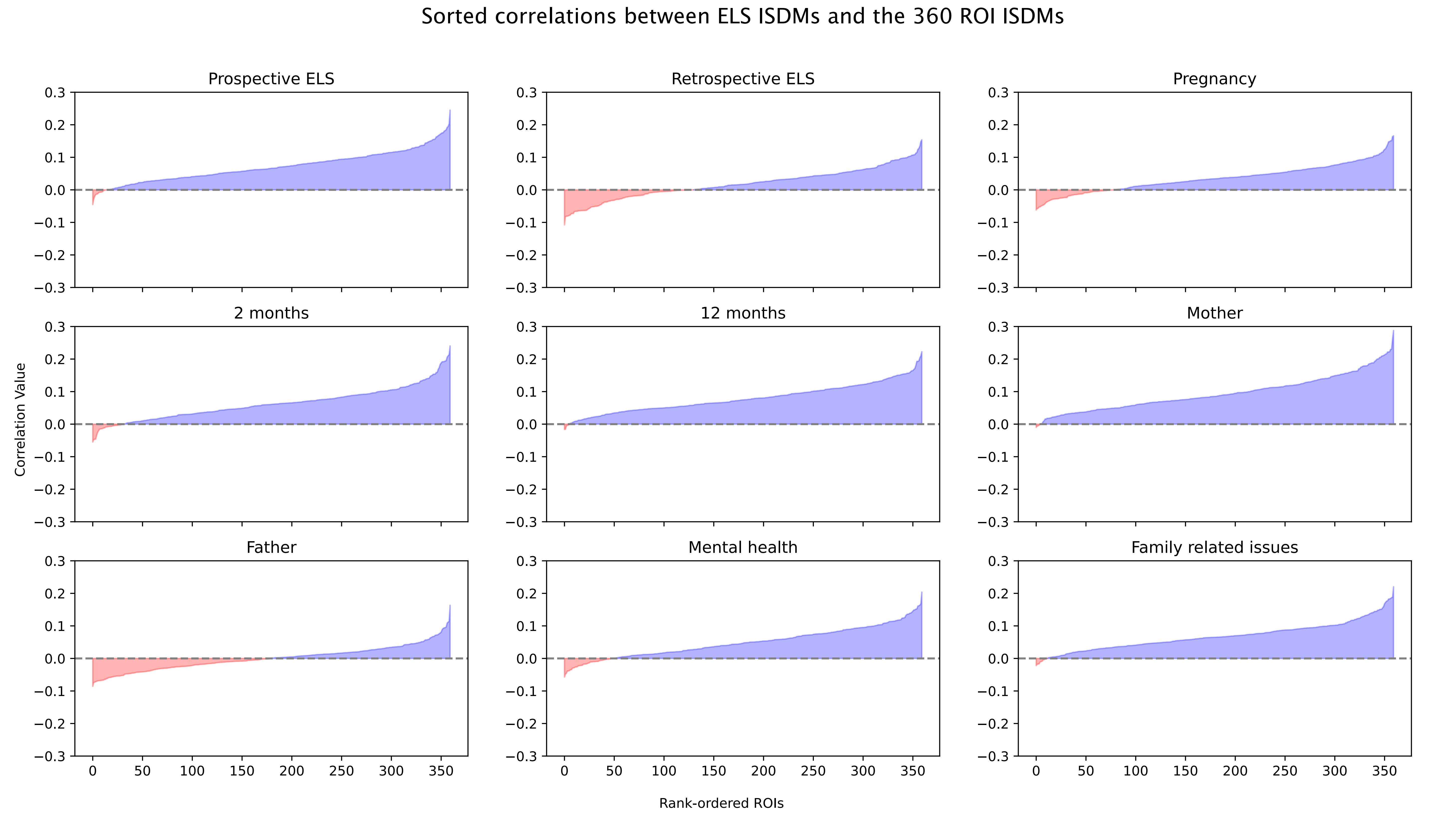

### Supplemental Figure 2

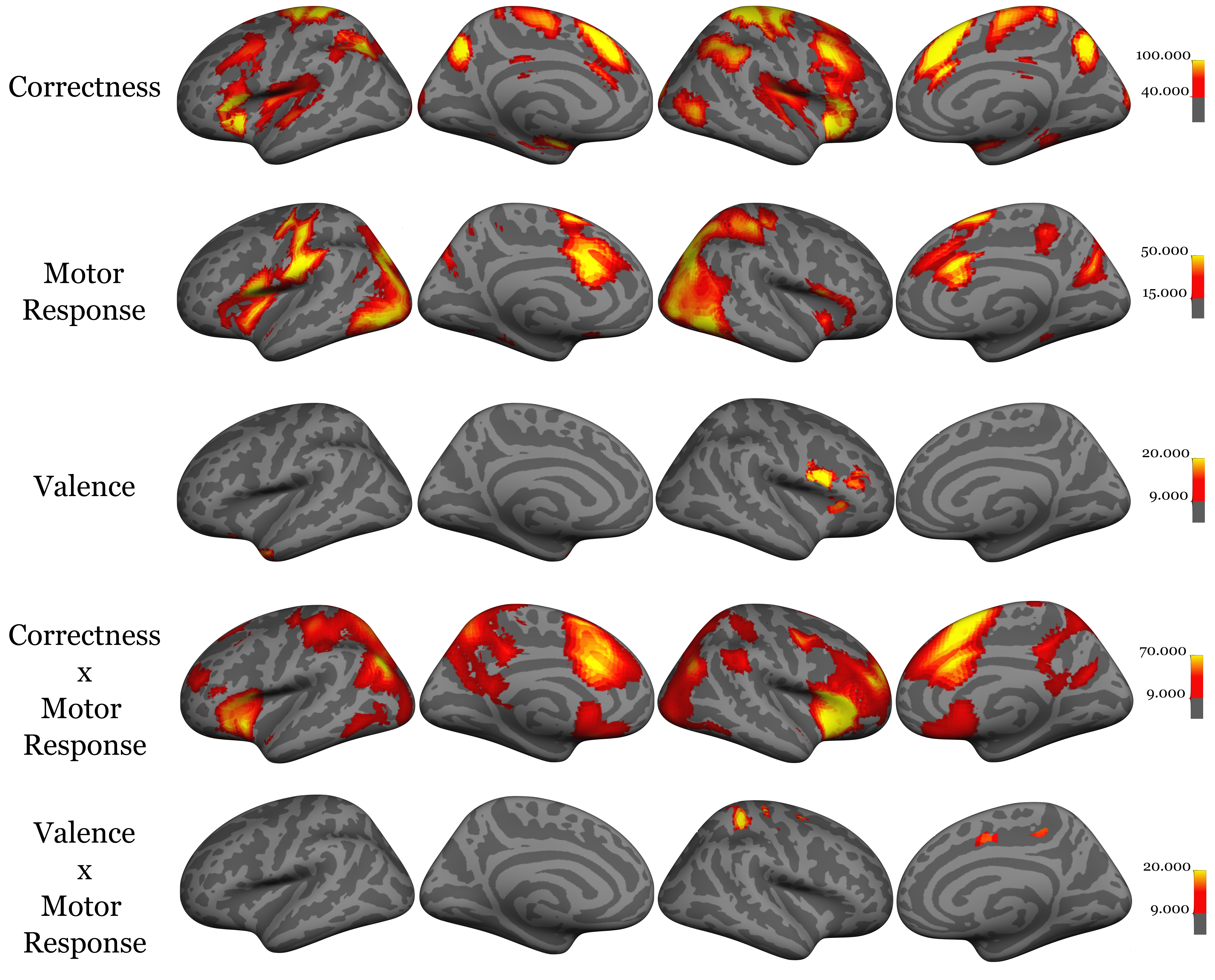
