## Supplemental Figures 3-41 for "Similarity in Early Life Stress Exposure is Associated with Similarity in Neural Representations in Early Adulthood"

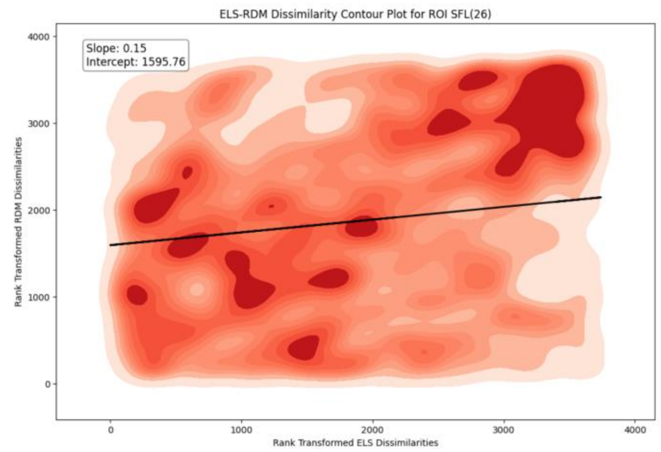
ELS-RDM Dissimilarity Contour Plot for ROI SFL(26)
Slope: 0.15
Intercept: 1595.76
Rank Transformed RDM Dissimilarities
Rank Transformed ELS Dissimilarities

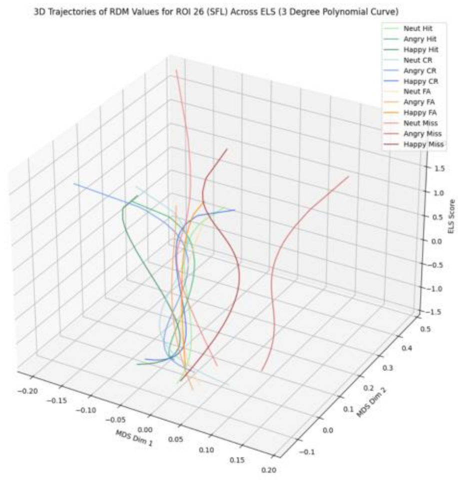
3D Trajectories of RDM Values for ROI 26 (SFL) Across ELS (3 Degree Polynomial Curve)
Neut Hit
Angry Hit
Happy Hit
Neut CR
Angry CR
Happy CR
Neut FA
Angry FA
Happy FA
Neut Miss
Angry Miss
Happy Miss
MDS Dim 1
MDS Dim 2
ELS Score

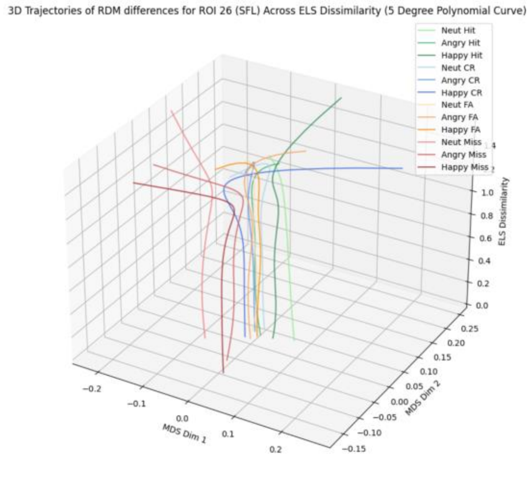
3D Trajectories of RDM differences for ROI 26 (SFL) Across ELS Dissimilarity (5 Degree Polynomial Curve)
Neut Hit
Angry Hit
Happy Hit
Neut CR
Angry CR
Happy CR
Neut FA
Angry FA
Happy FA
Neut Miss
Angry Miss
Happy Miss
MDS Dim 1
MDS Dim 2
ELS Dissimilarity

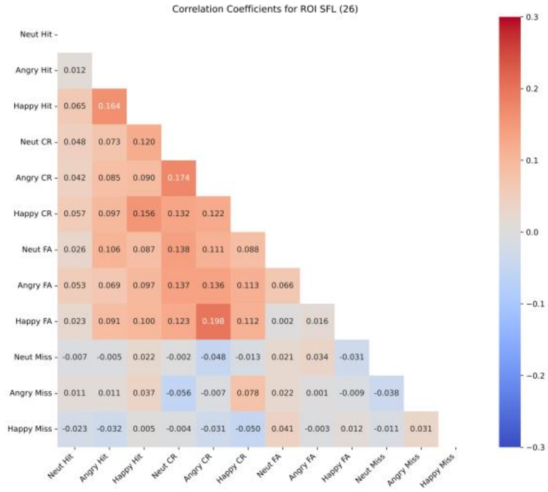
Correlation Coefficients for ROI SFL (26)
Neut Hit
Angry Hit
Happy Hit
Neut CR
Angry CR
Happy CR
Neut FA
Angry FA
Happy FA
Neut Miss
Angry Miss
Happy Miss

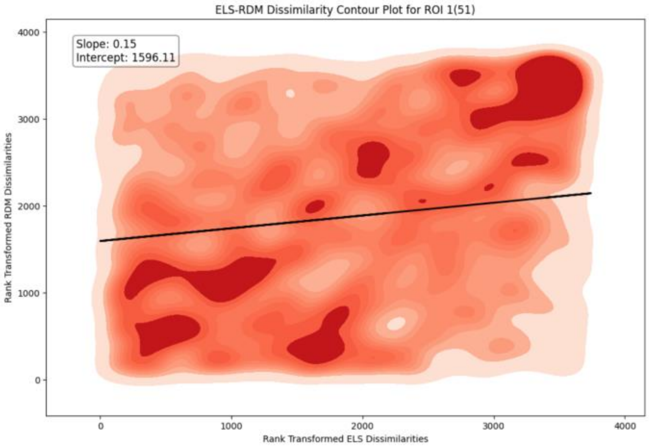
ELS-RDM Dissimilarity Contour Plot for ROI 1(51)
Slope: 0.15
Intercept: 1596.11
Rank Transformed RDM Dissimilarities
Rank Transformed ELS Dissimilarities

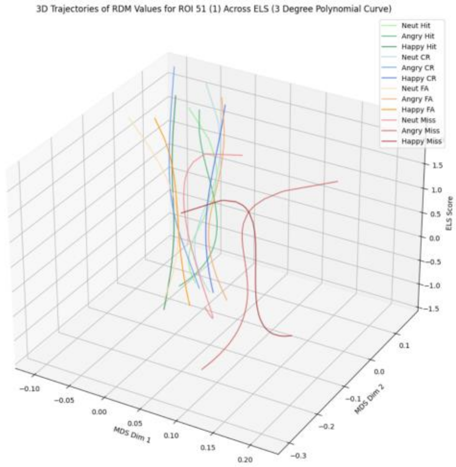
3D Trajectories of RDM Values for ROI 51 (1) Across ELS (3 Degree Polynomial Curve)
Neut Hit
Angry Hit
Happy Hit
Neut CR
Angry CR
Happy CR
Neut FA
Angry FA
Happy FA
Neut Miss
Angry Miss
Happy Miss
MDS Dim 1
MDS Dim 2
ELS Score

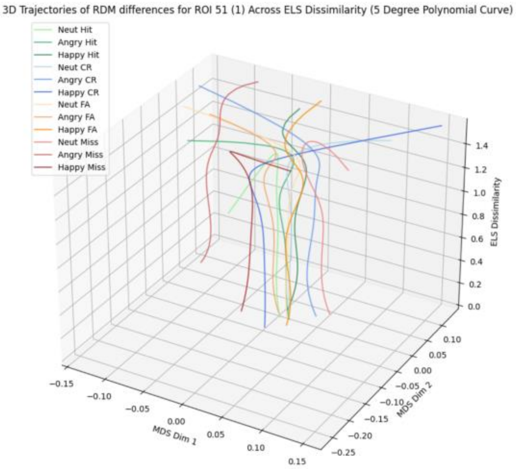
3D Trajectories of RDM differences for ROI 51 (1) Across ELS Dissimilarity (5 Degree Polynomial Curve)
Neut Hit
Angry Hit
Happy Hit
Neut CR
Angry CR
Happy CR
Neut FA
Angry FA
Happy FA
Neut Miss
Angry Miss
Happy Miss
MDS Dim 1
MDS Dim 2
ELS Dissimilarity

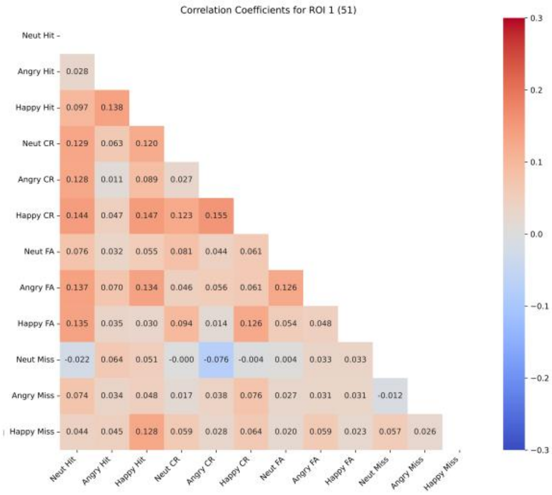
Correlation Coefficients for ROI 1 (51)
Neut Hit
Angry Hit 0.028
Happy Hit 0.097 0.138
Neut CR 0.129 0.063 0.120
Angry CR 0.128 0.011 0.089 0.027
Happy CR 0.144 0.047 0.147 0.123 0.155
Neut FA 0.076 0.032 0.055 0.081 0.044 0.061
Angry FA 0.137 0.070 0.134 0.046 0.056 0.061 0.126
Happy FA 0.135 0.035 0.030 0.094 0.014 0.126 0.054 0.048
Neut Miss -0.022 0.064 0.051 -0.000 -0.076 -0.004 0.004 0.033 0.033
Angry Miss 0.074 0.034 0.048 0.017 0.038 0.076 0.027 0.031 0.031 -0.012
Happy Miss 0.044 0.045 0.128 0.059 0.028 0.064 0.020 0.059 0.023 0.057 0.026
Neut Hit Angry Hit Happy Hit Neut CR Angry CR Happy CR Neut FA Angry FA Happy FA Neut Miss Angry Miss Happy Miss

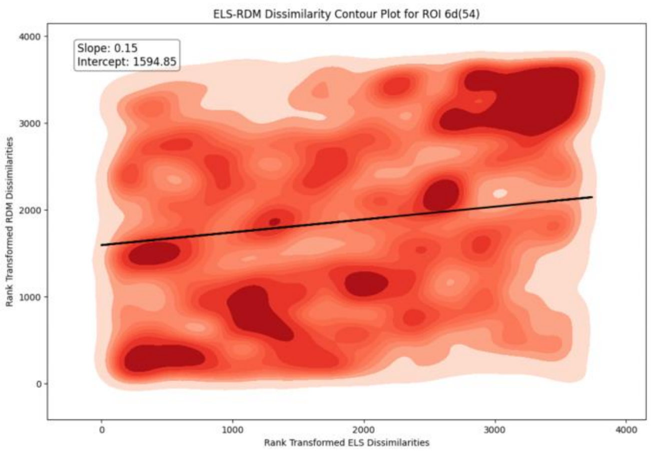
ELS-RDM Dissimilarity Contour Plot for ROI 6d(54)
Slope: 0.15
Intercept: 1594.85
Rank Transformed RDM Dissimilarities
Rank Transformed ELS Dissimilarities

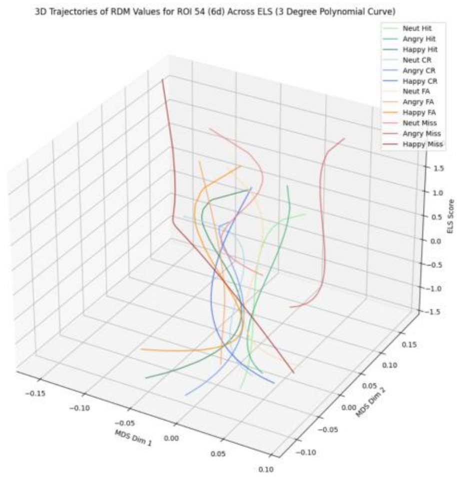
3D Trajectories of RDM Values for ROI 54 (6d) Across ELS (3 Degree Polynomial Curve)
Neut Hit
Angry Hit
Happy Hit
Neut CR
Angry CR
Happy CR
Neut FA
Angry FA
Happy FA
Neut Miss
Angry Miss
Happy Miss
ELS Score
MDS Dim 1
MDS Dim 2

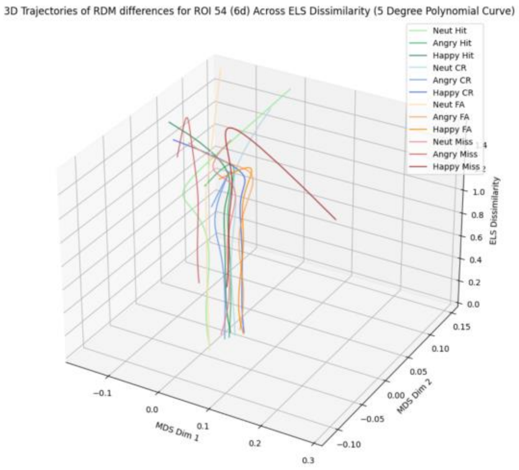
3D Trajectories of RDM differences for ROI 54 (6d) Across ELS Dissimilarity (5 Degree Polynomial Curve)
Neut Hit
Angry Hit
Happy Hit
Neut CR
Angry CR
Happy CR
Neut FA
Angry FA
Happy FA
Neut Miss
Angry Miss
Happy Miss
ELS Dissimilarity
MDS Dim 1
MDS Dim 2

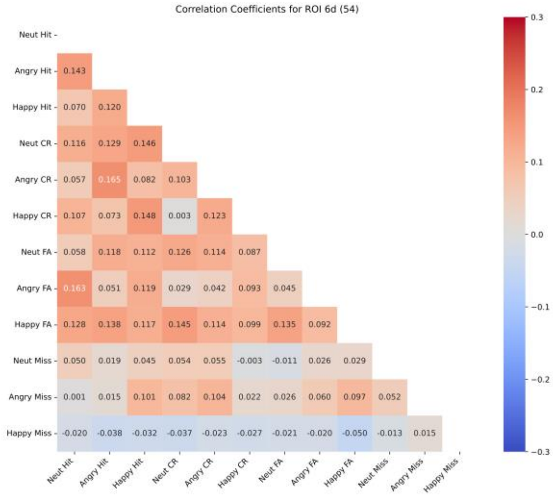
Correlation Coefficients for ROI 6d (54)
Neut Hit
Angry Hit 0.143
Happy Hit 0.070 0.120
Neut CR 0.116 0.129 0.146
Angry CR 0.057 0.165 0.082 0.103
Happy CR 0.107 0.073 0.148 0.003 0.123
Neut FA 0.058 0.118 0.112 0.126 0.114 0.087
Angry FA 0.163 0.051 0.119 0.029 0.042 0.093 0.045
Happy FA 0.128 0.138 0.117 0.145 0.114 0.099 0.135 0.092
Neut Miss 0.050 0.019 0.045 0.054 0.055 -0.003 -0.011 0.026 0.029
Angry Miss 0.001 0.015 0.101 0.082 0.104 0.022 0.026 0.060 0.097 0.052
Happy Miss -0.020 -0.038 -0.032 -0.037 -0.023 -0.027 -0.021 -0.020 -0.050 -0.013 0.015
Neut Hit Angry Hit Happy Hit Neut CR Angry CR Happy CR Neut FA Angry FA Happy FA Neut Miss Angry Miss Happy Miss

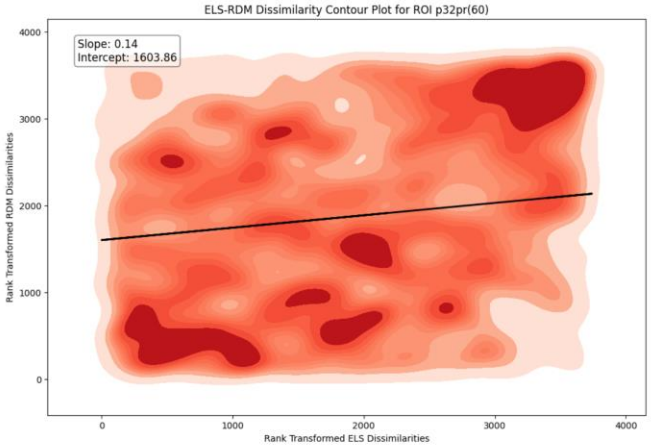
ELS-RDM Dissimilarity Contour Plot for ROI p32pr(60)
Slope: 0.14
Intercept: 1603.86
Rank Transformed RDM Dissimilarities
Rank Transformed ELS Dissimilarities

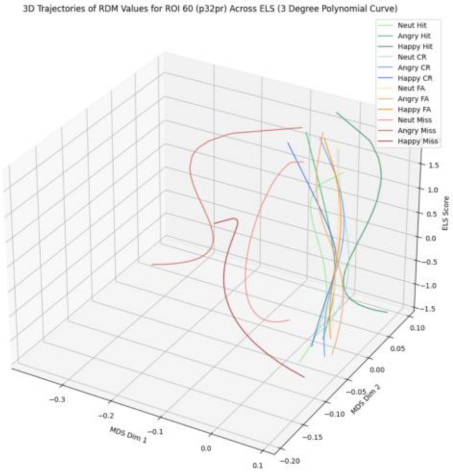
3D Trajectories of RDM Values for ROI 60 (p32pr) Across ELS (3 Degree Polynomial Curve)
Neut Hit
Angry Hit
Happy Hit
Neut CR
Angry CR
Happy CR
Neut FA
Angry FA
Happy FA
Neut Miss
Angry Miss
Happy Miss
ELS Score
MDS Dim 1
MDS Dim 2

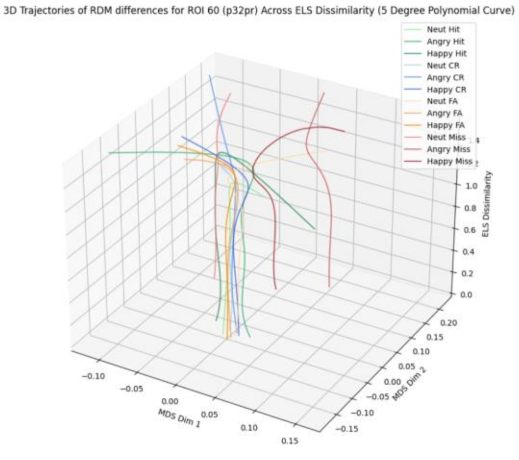
3D Trajectories of RDM differences for ROI 60 (p32pr) Across ELS Dissimilarity (5 Degree Polynomial Curve)
Neut Hit
Angry Hit
Happy Hit
Neut CR
Angry CR
Happy CR
Neut FA
Angry FA
Happy FA
Neut Miss
Angry Miss
Happy Miss
ELS Dissimilarity
MDS Dim 1
MDS Dim 2

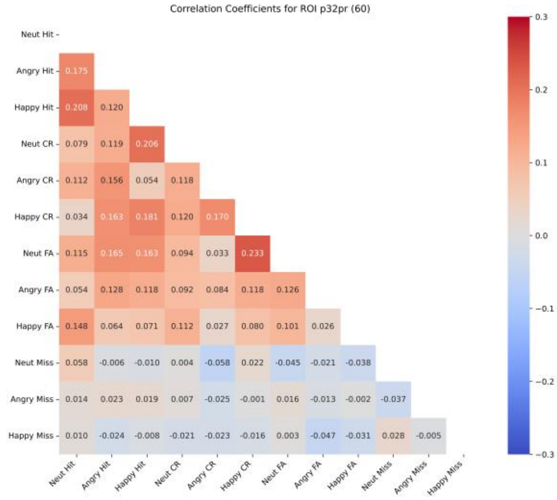
Correlation Coefficients for ROI p32pr (60)
Neut Hit
Angry Hit 0.175
Happy Hit 0.208 0.120
Neut CR 0.079 0.119 0.206
Angry CR 0.112 0.156 0.054 0.118
Happy CR 0.034 0.163 0.181 0.120 0.170
Neut FA 0.115 0.165 0.163 0.094 0.033 0.233
Angry FA 0.054 0.128 0.118 0.092 0.084 0.118 0.126
Happy FA 0.148 0.064 0.071 0.112 0.027 0.080 0.101 0.026
Neut Miss 0.058 -0.006 -0.010 0.004 -0.058 0.022 -0.045 -0.021 -0.038
Angry Miss 0.014 0.023 0.019 0.007 -0.025 -0.001 0.016 -0.013 -0.002 -0.037
Happy Miss 0.010 -0.024 -0.008 -0.021 -0.023 -0.016 0.003 -0.047 -0.031 0.028 -0.005
Neut Hit Angry Hit Happy Hit Neut CR Angry CR Happy CR Neut FA Angry FA Happy FA Neut Miss Angry Miss Happy Miss

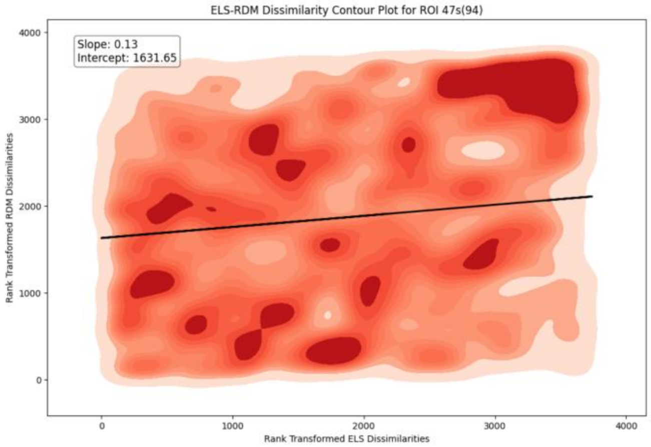
ELS-RDM Dissimilarity Contour Plot for ROI 47s(94)
Slope: 0.13
Intercept: 1631.65
Rank Transformed RDM Dissimilarities
Rank Transformed ELS Dissimilarities

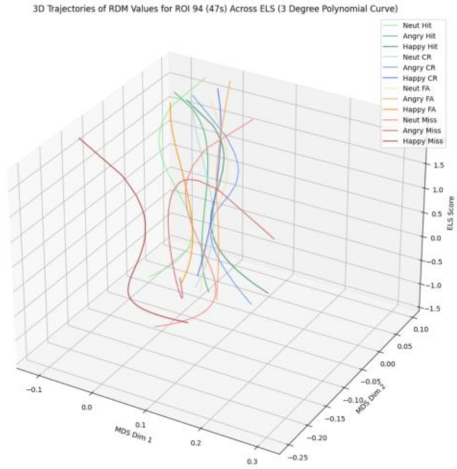
3D Trajectories of RDM Values for ROI 94 (47s) Across ELS (3 Degree Polynomial Curve)
Neut Hit
Angry Hit
Happy Hit
Neut CR
Angry CR
Happy CR
Neut FA
Angry FA
Happy FA
Neut Miss
Angry Miss
Happy Miss
ELS Score
MDS Dim 1
MDS Dim 2

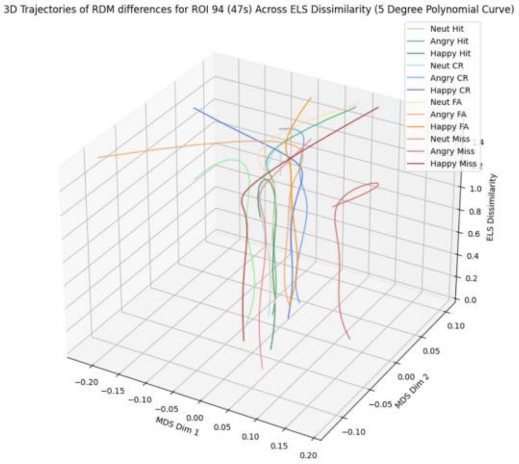
3D Trajectories of RDM differences for ROI 94 (47s) Across ELS Dissimilarity (5 Degree Polynomial Curve)
Neut Hit
Angry Hit
Happy Hit
Neut CR
Angry CR
Happy CR
Neut FA
Angry FA
Happy FA
Neut Miss
Angry Miss
Happy Miss
ELS Dissimilarity
MDS Dim 1
MDS Dim 2

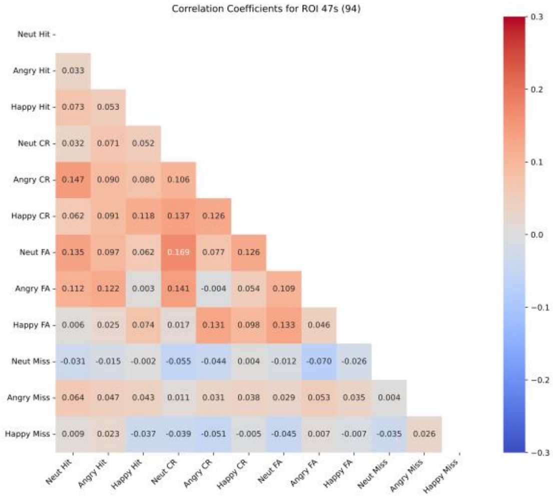
Correlation Coefficients for ROI 47s (94)
Neut Hit
Angry Hit 0.033
Happy Hit 0.073 0.053
Neut CR 0.032 0.071 0.052
Angry CR 0.147 0.090 0.080 0.106
Happy CR 0.062 0.091 0.118 0.137 0.126
Neut FA 0.135 0.097 0.062 0.169 0.077 0.126
Angry FA 0.112 0.122 0.003 0.141 -0.004 0.054 0.109
Happy FA 0.006 0.025 0.074 0.017 0.131 0.098 0.133 0.046
Neut Miss -0.031 -0.015 -0.002 -0.055 -0.044 0.004 -0.012 -0.070 -0.026
Angry Miss 0.064 0.047 0.043 0.011 0.031 0.038 0.029 0.053 0.035 0.004
Happy Miss 0.009 0.023 -0.037 -0.039 -0.051 -0.005 -0.045 0.007 -0.007 -0.035 0.026
Neut Hit Angry Hit Happy Hit Neut CR Angry CR Happy CR Neut FA Angry FA Happy FA Neut Miss Angry Miss Happy Miss

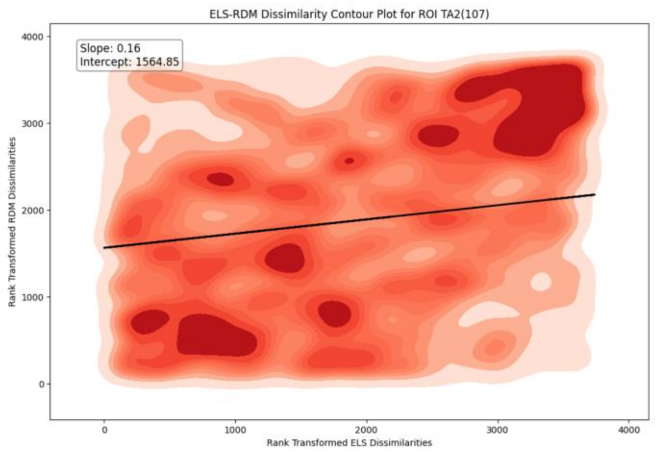
ELS-RDM Dissimilarity Contour Plot for ROI TA2(107)
Slope: 0.16
Intercept: 1564.85
Rank Transformed RDM Dissimilarities
Rank Transformed ELS Dissimilarities

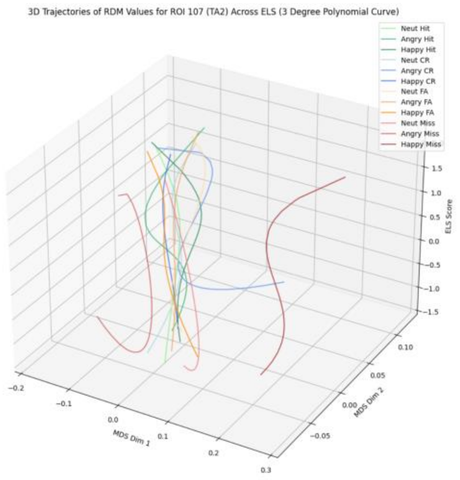
3D Trajectories of RDM Values for ROI 107 (TA2) Across ELS (3 Degree Polynomial Curve)
Neut Hit
Angry Hit
Happy Hit
Neut CR
Angry CR
Happy CR
Neut FA
Angry FA
Happy FA
Neut Miss
Angry Miss
Happy Miss
ELS Score
MDS Dim 1
MDS Dim 2

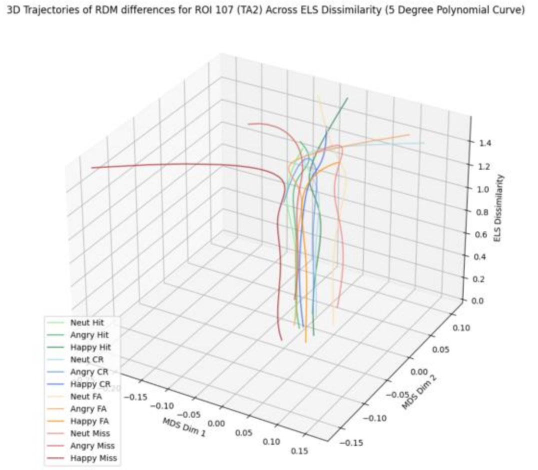
3D Trajectories of RDM differences for ROI 107 (TA2) Across ELS Dissimilarity (5 Degree Polynomial Curve)
ELS Dissimilarity
Neut Hit
Angry Hit
Happy Hit
Neut CR
Angry CR
Happy CR
Neut FA
Angry FA
Happy FA
Neut Miss
Angry Miss
Happy Miss
MDS Dim 1
MDS Dim 2

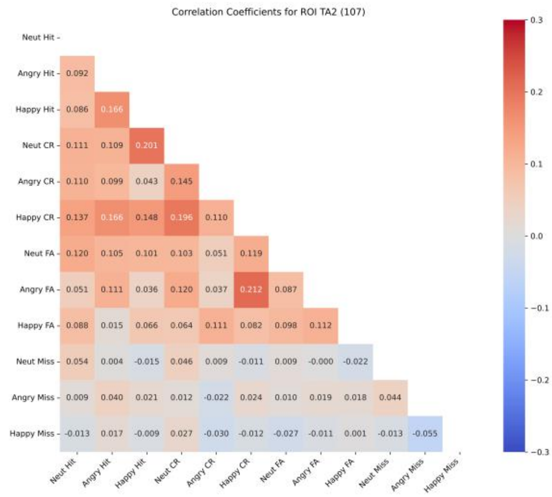
Correlation Coefficients for ROI TA2 (107)
Neut Hit
Angry Hit 0.092
Happy Hit 0.086 0.166
Neut CR 0.111 0.109 0.201
Angry CR 0.110 0.099 0.043 0.145
Happy CR 0.137 0.166 0.148 0.196 0.110
Neut FA 0.120 0.105 0.101 0.103 0.051 0.119
Angry FA 0.051 0.111 0.036 0.120 0.037 0.212 0.087
Happy FA 0.088 0.015 0.066 0.064 0.111 0.082 0.098 0.112
Neut Miss 0.054 0.004 -0.015 0.046 0.009 -0.011 0.009 -0.000 -0.022
Angry Miss 0.009 0.040 0.021 0.012 -0.022 0.024 0.010 0.019 0.018 0.044
Happy Miss -0.013 0.017 -0.009 0.027 -0.030 -0.012 -0.027 -0.011 0.001 -0.013 -0.055
Neut Hit Angry Hit Happy Hit Neut CR Angry CR Happy CR Neut FA Angry FA Happy FA Neut Miss Angry Miss Happy Miss

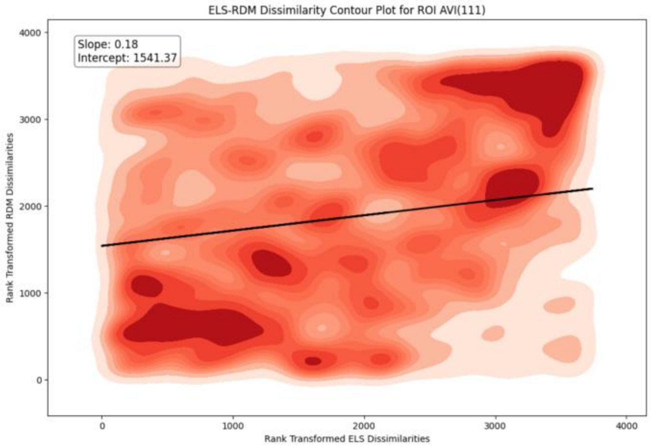
ELS-RDM Dissimilarity Contour Plot for ROI AVI(111)
Slope: 0.18
Intercept: 1541.37
Rank Transformed RDM Dissimilarities
Rank Transformed ELS Dissimilarities
0
1000
2000
3000
4000

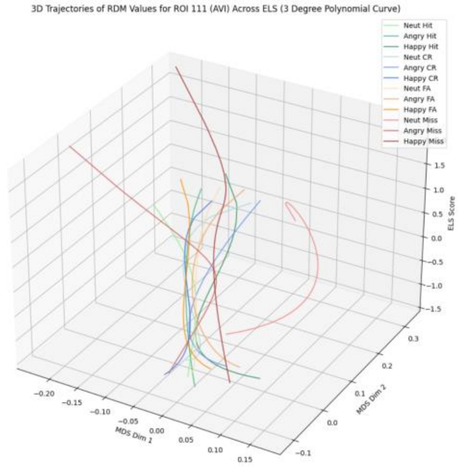
3D Trajectories of RDM Values for ROI 111 (AVI) Across ELS (3 Degree Polynomial Curve)
Neut Hit
Angry Hit
Happy Hit
Neut CR
Angry CR
Happy CR
Neut FA
Angry FA
Happy FA
Neut Miss
Angry Miss
Happy Miss
ELS Score
MDS Dim 1
MDS Dim 2

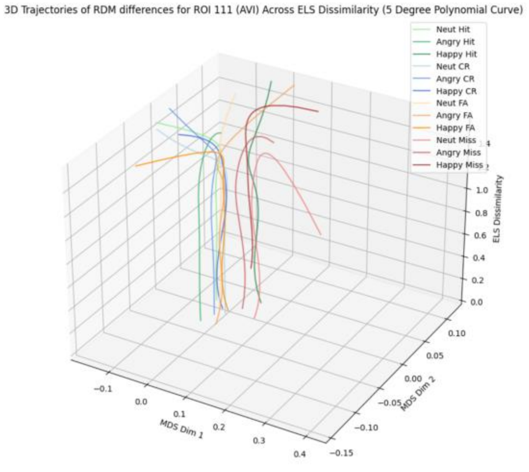
3D Trajectories of RDM differences for ROI 111 (AVI) Across ELS Dissimilarity (5 Degree Polynomial Curve)
Neut Hit
Angry Hit
Happy Hit
Neut CR
Angry CR
Happy CR
Neut FA
Angry FA
Happy FA
Neut Miss
Angry Miss
Happy Miss
ELS Dissimilarity
MDS Dim 1
MDS Dim 2

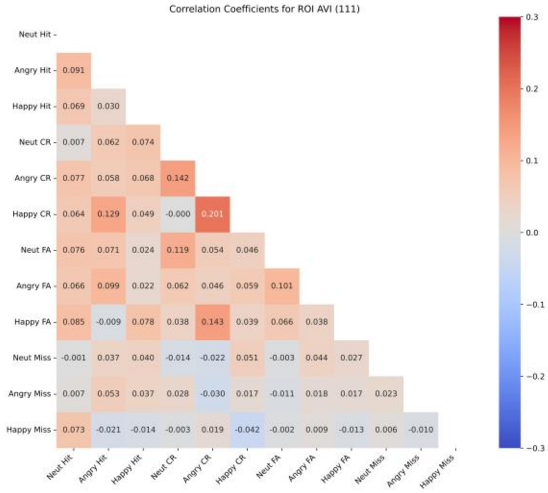
Correlation Coefficients for ROI AVI (111)
Neut Hit
Angry Hit 0.091
Happy Hit 0.069 0.030
Neut CR 0.007 0.062 0.074
Angry CR 0.077 0.058 0.068 0.142
Happy CR 0.064 0.129 0.049 -0.000 0.201
Neut FA 0.076 0.071 0.024 0.119 0.054 0.046
Angry FA 0.066 0.099 0.022 0.062 0.046 0.059 0.101
Happy FA 0.085 -0.009 0.078 0.038 0.143 0.039 0.066 0.038
Neut Miss -0.001 0.037 0.040 -0.014 -0.022 0.051 -0.003 0.044 0.027
Angry Miss 0.007 0.053 0.037 0.028 -0.030 0.017 -0.011 0.018 0.017 0.023
Happy Miss 0.073 -0.021 -0.014 -0.003 0.019 -0.042 -0.002 0.009 -0.013 0.006 -0.010
Neut Hit Angry Hit Happy Hit Neut CR Angry CR Happy CR Neut FA Angry FA Happy FA Neut Miss Angry Miss Happy Miss

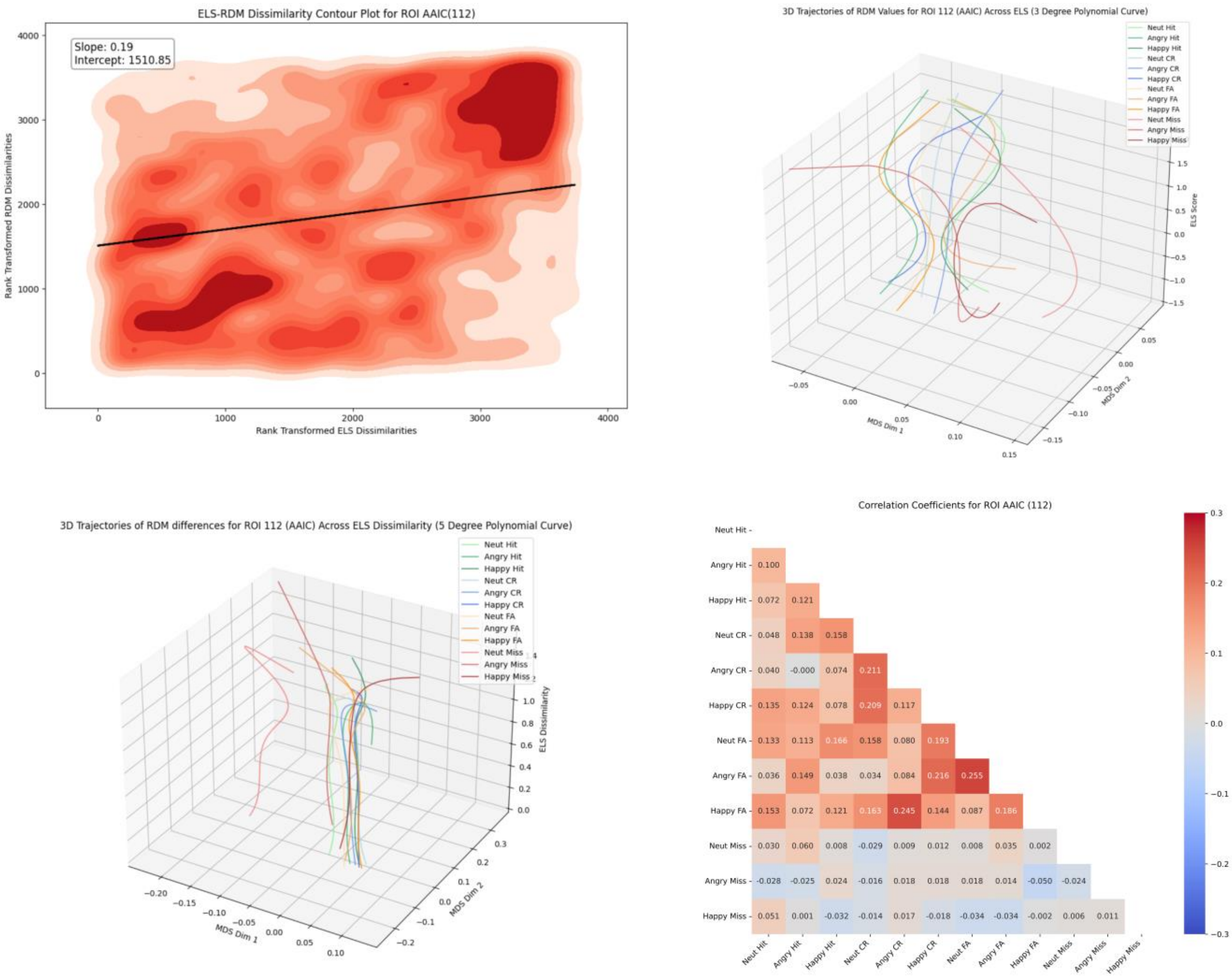
ELS-RDM Dissimilarity Contour Plot for ROI AAIC(112)
Slope: 0.19
Intercept: 1510.85
Rank Transformed RDM Dissimilarities
Rank Transformed ELS Dissimilarities
3D Trajectories of RDM Values for ROI 112 (AAIC) Across ELS (3 Degree Polynomial Curve)
Neut Hit
Angry Hit
Happy Hit
Neut CR
Angry CR
Happy CR
Neut FA
Angry FA
Happy FA
Neut Miss
Angry Miss
Happy Miss
ELS Score
MDS Dim 1
MDS Dim 2
3D Trajectories of RDM differences for ROI 112 (AAIC) Across ELS Dissimilarity (5 Degree Polynomial Curve)
ELS Dissimilarity
Correlation Coefficients for ROI AAIC (112)

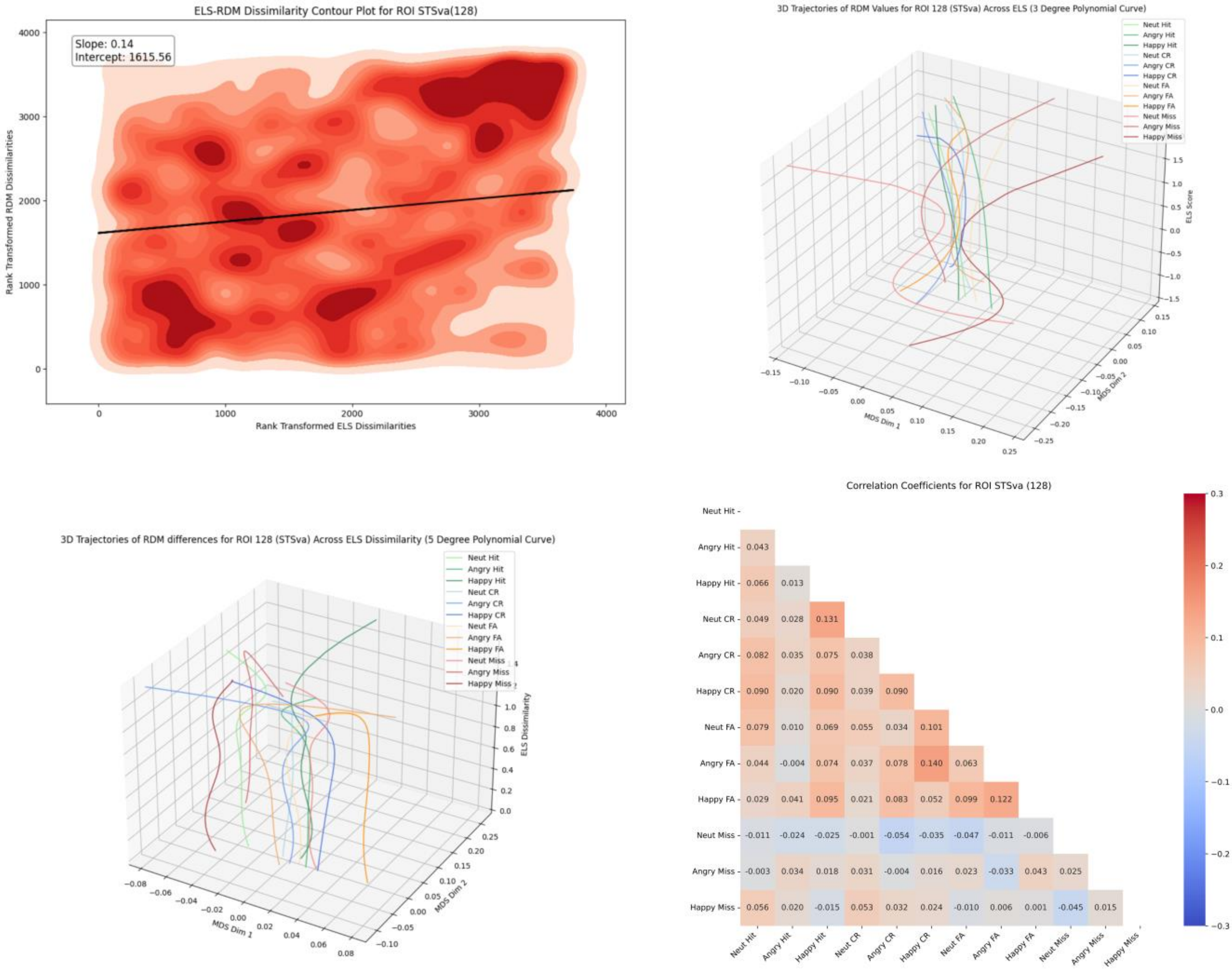

ELS-RDM Dissimilarity Contour Plot for ROI STSva(128)
Slope: 0.14
Intercept: 1615.56
Rank Transformed RDM Dissimilarities
Rank Transformed ELS Dissimilarities
3D Trajectories of RDM Values for ROI 128 (STSva) Across ELS (3 Degree Polynomial Curve)
ELS Score
MDS Dim 1
MDS Dim 2
Neut Hit
Angry Hit
Happy Hit
Neut CR
Angry CR
Happy CR
Neut FA
Angry FA
Happy FA
Neut Miss
Angry Miss
Happy Miss
3D Trajectories of RDM differences for ROI 128 (STSva) Across ELS Dissimilarity (5 Degree Polynomial Curve)
ELS Dissimilarity
MDS Dim 1
MDS Dim 2
Correlation Coefficients for ROI STSva (128)
Neut Hit
Angry Hit 0.043
Happy Hit 0.066 0.013
Neut CR 0.049 0.028 0.131
Angry CR 0.082 0.035 0.075 0.038
Happy CR 0.090 0.020 0.090 0.039 0.090
Neut FA 0.079 0.010 0.069 0.055 0.034 0.101
Angry FA 0.044 -0.004 0.074 0.037 0.078 0.140 0.063
Happy FA 0.029 0.041 0.095 0.021 0.083 0.052 0.099 0.122
Neut Miss -0.011 -0.024 -0.025 -0.001 -0.054 -0.035 -0.047 -0.011 -0.006
Angry Miss -0.003 0.034 0.018 0.031 -0.004 0.016 0.023 -0.033 0.043 0.025
Happy Miss 0.056 0.020 -0.015 0.053 0.032 0.024 -0.010 0.006 0.001 -0.045 0.015

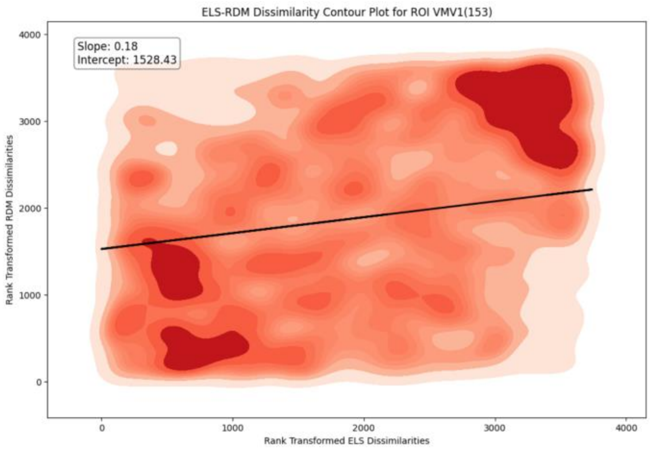
ELS-RDM Dissimilarity Contour Plot for ROI VMV1(153)
Slope: 0.18
Intercept: 1528.43
Rank Transformed RDM Dissimilarities
Rank Transformed ELS Dissimilarities

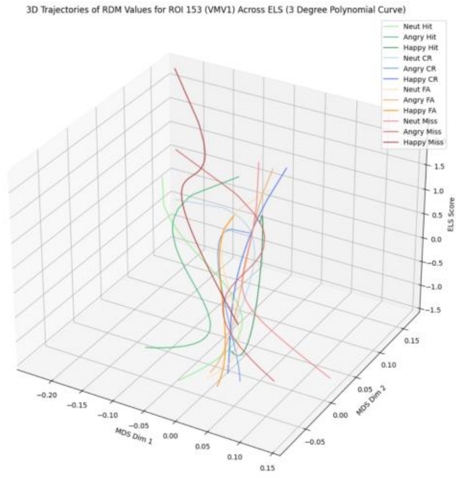
3D Trajectories of RDM Values for ROI 153 (VMV1) Across ELS (3 Degree Polynomial Curve)
Neut Hit
Angry Hit
Happy Hit
Neut CR
Angry CR
Happy CR
Neut FA
Angry FA
Happy FA
Neut Miss
Angry Miss
Happy Miss
ELS Score
MDS Dim 1
MDS Dim 2

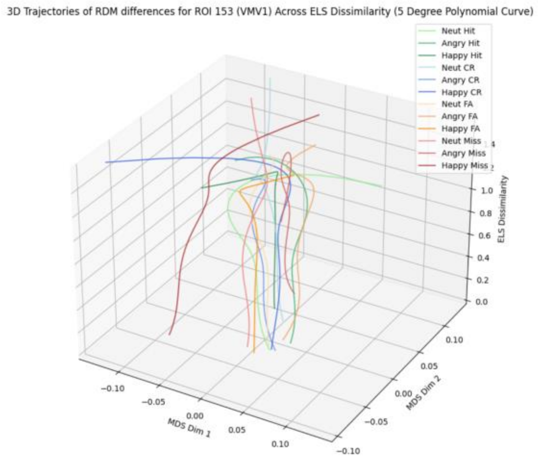
3D Trajectories of RDM differences for ROI 153 (VMV1) Across ELS Dissimilarity (5 Degree Polynomial Curve)
Neut Hit
Angry Hit
Happy Hit
Neut CR
Angry CR
Happy CR
Neut FA
Angry FA
Happy FA
Neut Miss
Angry Miss
Happy Miss
ELS Dissimilarity
MDS Dim 1
MDS Dim 2

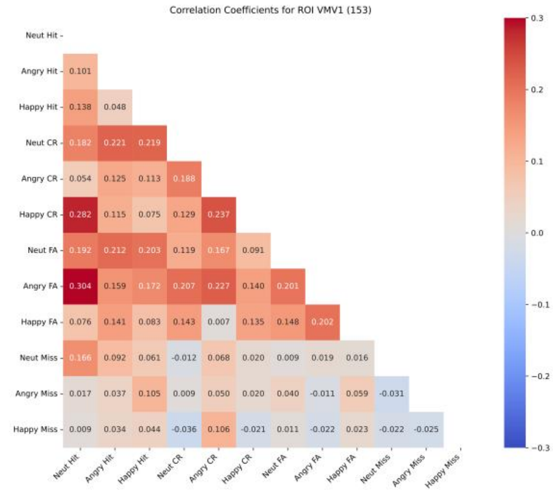
Correlation Coefficients for ROI VMV1 (153)
Neut Hit
Angry Hit 0.101
Happy Hit 0.138 0.048
Neut CR 0.182 0.221 0.219
Angry CR 0.054 0.125 0.113 0.188
Happy CR 0.282 0.115 0.075 0.129 0.237
Neut FA 0.192 0.212 0.203 0.119 0.167 0.091
Angry FA 0.304 0.159 0.172 0.207 0.227 0.140 0.201
Happy FA 0.076 0.141 0.083 0.143 0.007 0.135 0.148 0.202
Neut Miss 0.166 0.092 0.061 -0.012 0.068 0.020 0.009 0.019 0.016
Angry Miss 0.017 0.037 0.105 0.009 0.050 0.020 0.040 -0.011 0.059 -0.031
Happy Miss 0.009 0.034 0.044 -0.036 0.106 -0.021 0.011 -0.022 0.023 -0.022 -0.025
Neut Hit
Angry Hit
Happy Hit
Neut CR
Angry CR
Happy CR
Neut FA
Angry FA
Happy FA
Neut Miss
Angry Miss
Happy Miss

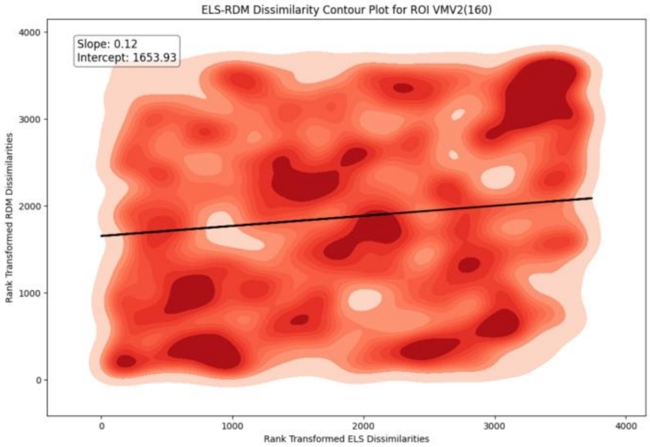
ELS-RDM Dissimilarity Contour Plot for ROI VMV2(160)
Slope: 0.12
Intercept: 1653.93
Rank Transformed RDM Dissimilarities
Rank Transformed ELS Dissimilarities

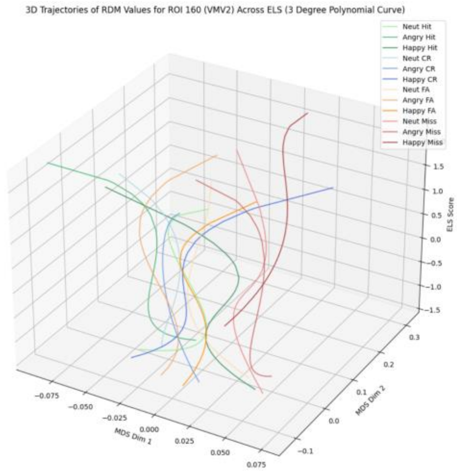
3D Trajectories of RDM Values for ROI 160 (VMV2) Across ELS (3 Degree Polynomial Curve)
Neut Hit
Angry Hit
Happy Hit
Neut CR
Angry CR
Happy CR
Neut FA
Angry FA
Happy FA
Neut Miss
Angry Miss
Happy Miss
ELS Score
MDS Dim 1
MDS Dim 2

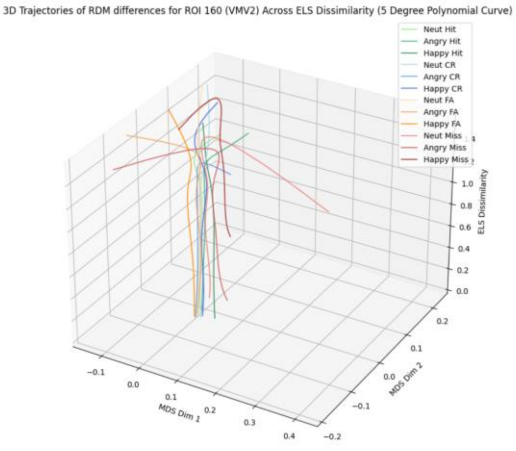
3D Trajectories of RDM differences for ROI 160 (VMV2) Across ELS Dissimilarity (5 Degree Polynomial Curve)
Neut Hit
Angry Hit
Happy Hit
Neut CR
Angry CR
Happy CR
Neut FA
Angry FA
Happy FA
Neut Miss
Angry Miss
Happy Miss
ELS Dissimilarity
MDS Dim 1
MDS Dim 2

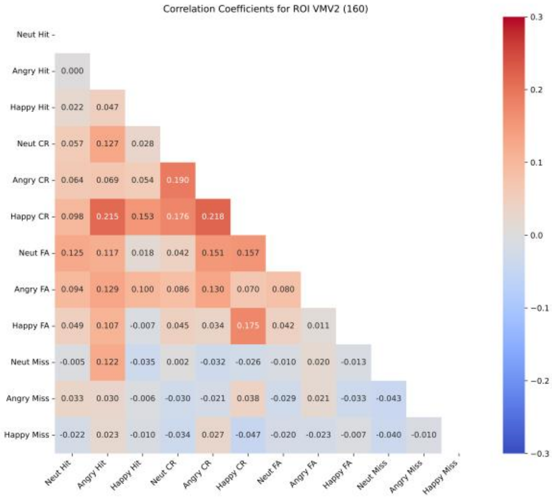
Correlation Coefficients for ROI VMV2 (160)
Neut Hit
Angry Hit 0.000
Happy Hit 0.022 0.047
Neut CR 0.057 0.127 0.028
Angry CR 0.064 0.069 0.054 0.190
Happy CR 0.098 0.215 0.153 0.176 0.218
Neut FA 0.125 0.117 0.018 0.042 0.151 0.157
Angry FA 0.094 0.129 0.100 0.086 0.130 0.070 0.080
Happy FA 0.049 0.107 -0.007 0.045 0.034 0.175 0.042 0.011
Neut Miss -0.005 0.122 -0.035 0.002 -0.032 -0.026 -0.010 0.020 -0.013
Angry Miss 0.033 0.030 -0.006 -0.030 -0.021 0.038 -0.029 0.021 -0.033 -0.043
Happy Miss -0.022 0.023 -0.010 -0.034 0.027 -0.047 -0.020 -0.023 -0.007 -0.040 -0.010
Neut Hit Angry Hit Happy Hit Neut CR Angry CR Happy CR Neut FA Angry FA Happy FA Neut Miss Angry Miss Happy Miss

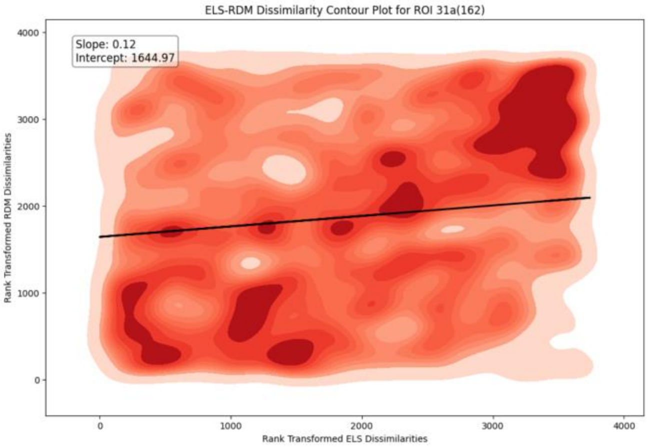
ELS-RDM Dissimilarity Contour Plot for ROI 31a(162)
Slope: 0.12
Intercept: 1644.97
Rank Transformed RDM Dissimilarities
Rank Transformed ELS Dissimilarities

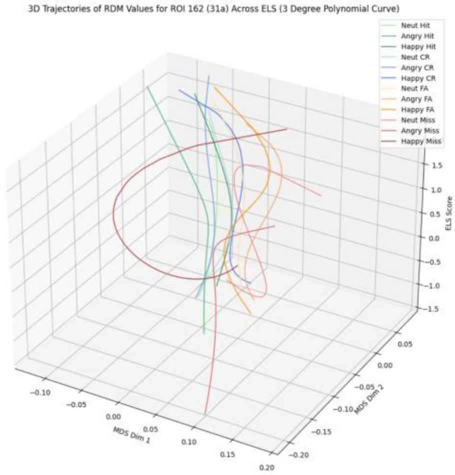
3D Trajectories of RDM Values for ROI 162 (31a) Across ELS (3 Degree Polynomial Curve)
Neut Hit
Angry Hit
Happy Hit
Neut CR
Angry CR
Happy CR
Neut FA
Angry FA
Happy FA
Neut Miss
Angry Miss
Happy Miss
ELS Score
MDS Dim 1
MDS Dim 2

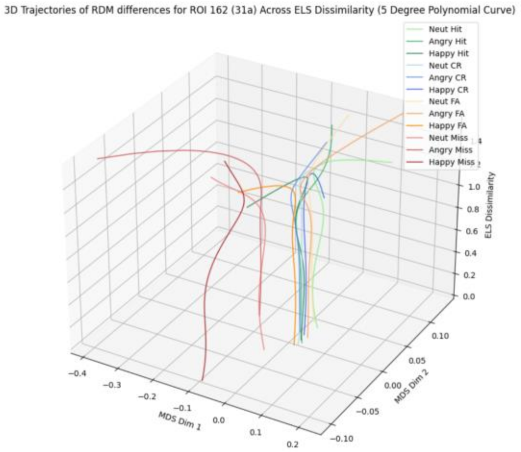
3D Trajectories of RDM differences for ROI 162 (31a) Across ELS Dissimilarity (5 Degree Polynomial Curve)
Neut Hit
Angry Hit
Happy Hit
Neut CR
Angry CR
Happy CR
Neut FA
Angry FA
Happy FA
Neut Miss
Angry Miss
Happy Miss
ELS Dissimilarity
MDS Dim 1
MDS Dim 2

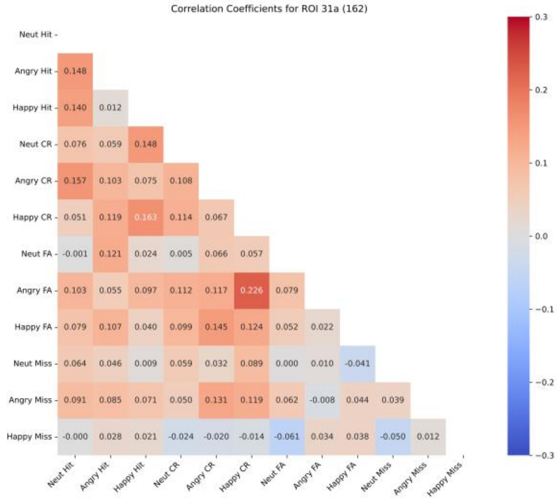
Correlation Coefficients for ROI 31a (162)
Neut Hit
Angry Hit 0.148
Happy Hit 0.140 0.012
Neut CR 0.076 0.059 0.148
Angry CR 0.157 0.103 0.075 0.108
Happy CR 0.051 0.119 0.163 0.114 0.067
Neut FA -0.001 0.121 0.024 0.005 0.066 0.057
Angry FA 0.103 0.055 0.097 0.112 0.117 0.226 0.079
Happy FA 0.079 0.107 0.040 0.099 0.145 0.124 0.052 0.022
Neut Miss 0.064 0.046 0.009 0.059 0.032 0.089 0.000 0.010 -0.041
Angry Miss 0.091 0.085 0.071 0.050 0.131 0.119 0.062 -0.008 0.044 0.039
Happy Miss -0.000 0.028 0.021 -0.024 -0.020 -0.014 -0.061 0.034 0.038 -0.050 0.012
Neut Hit Angry Hit Happy Hit Neut CR Angry CR Happy CR Neut FA Angry FA Happy FA Neut Miss Angry Miss Happy Miss

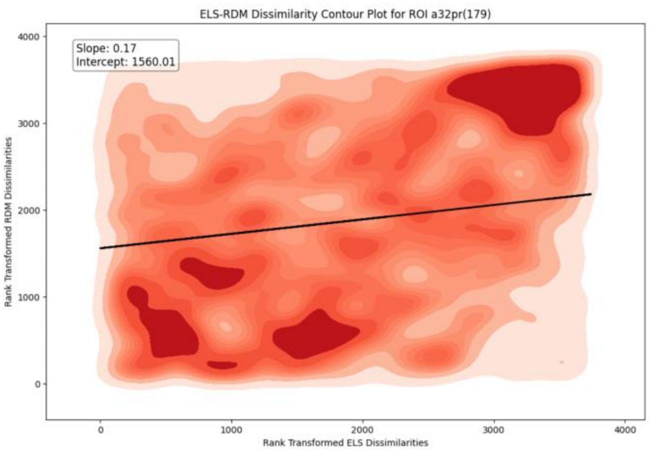
ELS-RDM Dissimilarity Contour Plot for ROI a32pr(179)
Slope: 0.17
Intercept: 1560.01
Rank Transformed RDM Dissimilarities
Rank Transformed ELS Dissimilarities

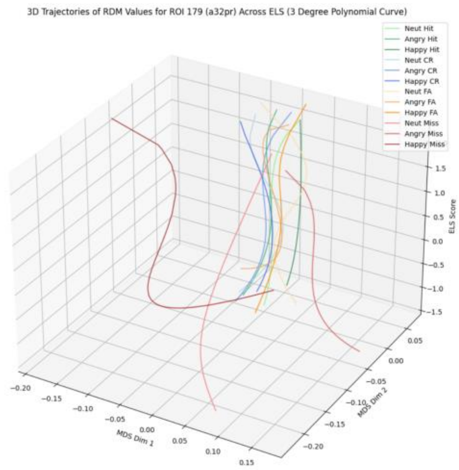
3D Trajectories of RDM Values for ROI 179 (a32pr) Across ELS (3 Degree Polynomial Curve)
Neut Hit
Angry Hit
Happy Hit
Neut CR
Angry CR
Happy CR
Neut FA
Angry FA
Happy FA
Neut Miss
Angry Miss
Happy Miss
ELS Score
MDS Dim 1
MDS Dim 2

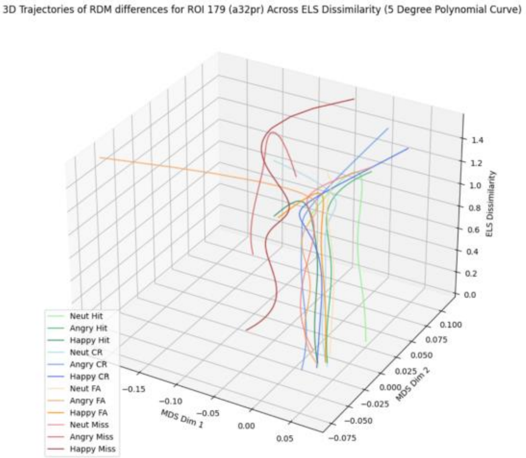
3D Trajectories of RDM differences for ROI 179 (a32pr) Across ELS Dissimilarity (5 Degree Polynomial Curve)
ELS Dissimilarity
MDS Dim 1
MDS Dim 2
Neut Hit
Angry Hit
Happy Hit
Neut CR
Angry CR
Happy CR
Neut FA
Angry FA
Happy FA
Neut Miss
Angry Miss
Happy Miss

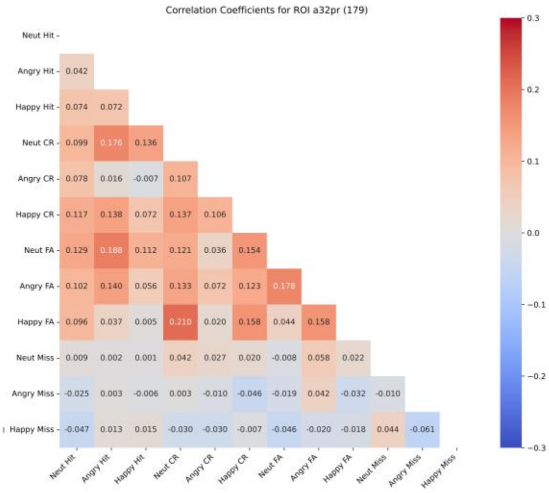
Correlation Coefficients for ROI a32pr (179)
Neut Hit
Angry Hit 0.042
Happy Hit 0.074 0.072
Neut CR 0.099 0.176 0.136
Angry CR 0.078 0.016 -0.007 0.107
Happy CR 0.117 0.138 0.072 0.137 0.106
Neut FA 0.129 0.188 0.112 0.121 0.036 0.154
Angry FA 0.102 0.140 0.056 0.133 0.072 0.123 0.178
Happy FA 0.096 0.037 0.005 0.210 0.020 0.158 0.044 0.158
Neut Miss 0.009 0.002 0.001 0.042 0.027 0.020 -0.008 0.058 0.022
Angry Miss -0.025 0.003 -0.006 0.003 -0.010 -0.046 -0.019 0.042 -0.032 -0.010
Happy Miss -0.047 0.013 0.015 -0.030 -0.030 -0.007 -0.046 -0.020 -0.018 0.044 -0.061
Neut Hit Angry Hit Happy Hit Neut CR Angry CR Happy CR Neut FA Angry FA Happy FA Neut Miss Angry Miss Happy Miss

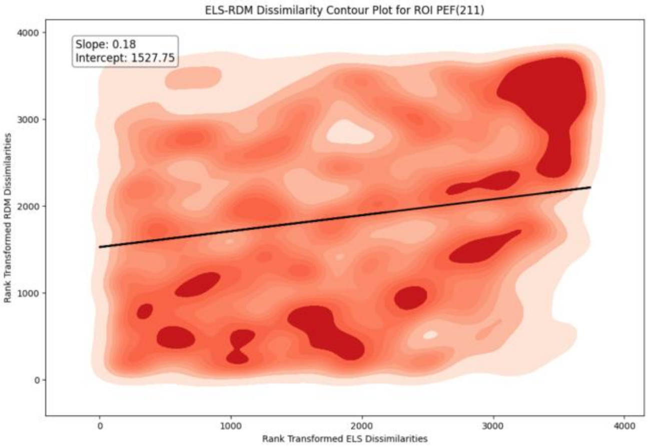
ELS-RDM Dissimilarity Contour Plot for ROI PEF(211)
Slope: 0.18
Intercept: 1527.75
Rank Transformed RDM Dissimilarities
Rank Transformed ELS Dissimilarities
0
1000
2000
3000
4000

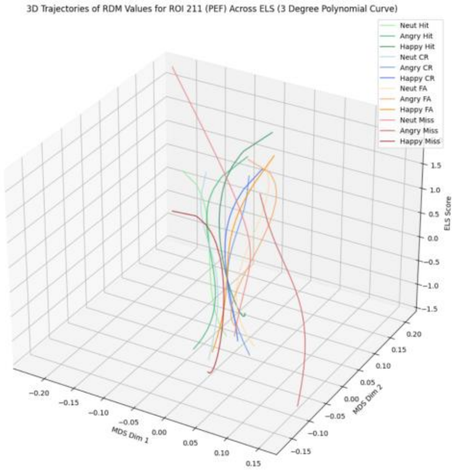
3D Trajectories of RDM Values for ROI 211 (PEF) Across ELS (3 Degree Polynomial Curve)
Neut Hit
Angry Hit
Happy Hit
Neut CR
Angry CR
Happy CR
Neut FA
Angry FA
Happy FA
Neut Miss
Angry Miss
Happy Miss
MDS Dim 1
MDS Dim 2
ELS Score

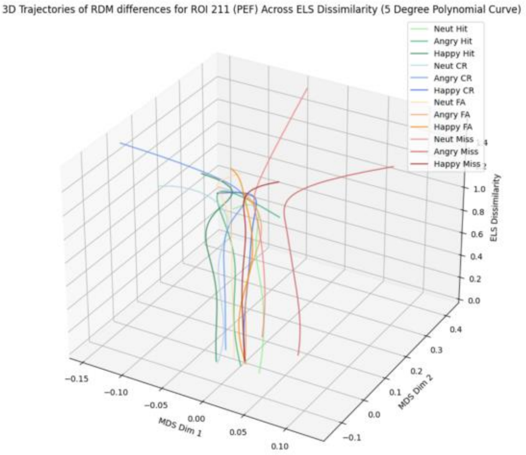
3D Trajectories of RDM differences for ROI 211 (PEF) Across ELS Dissimilarity (5 Degree Polynomial Curve)
Neut Hit
Angry Hit
Happy Hit
Neut CR
Angry CR
Happy CR
Neut FA
Angry FA
Happy FA
Neut Miss
Angry Miss
Happy Miss
MDS Dim 1
MDS Dim 2
ELS Dissimilarity

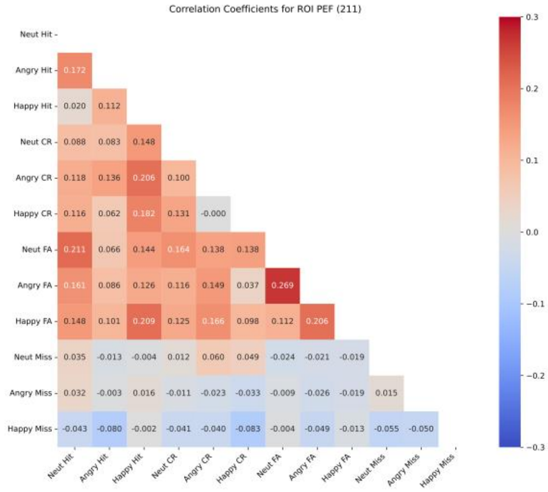
Correlation Coefficients for ROI PEF (211)
Neut Hit
Angry Hit 0.172
Happy Hit 0.020 0.112
Neut CR 0.088 0.083 0.148
Angry CR 0.118 0.136 0.206 0.100
Happy CR 0.116 0.062 0.182 0.131 -0.000
Neut FA 0.211 0.066 0.144 0.164 0.138 0.138
Angry FA 0.161 0.086 0.126 0.116 0.149 0.037 0.269
Happy FA 0.148 0.101 0.209 0.125 0.166 0.098 0.112 0.206
Neut Miss 0.035 -0.013 -0.004 0.012 0.060 0.049 -0.024 -0.021 -0.019
Angry Miss 0.032 -0.003 0.016 -0.011 -0.023 -0.033 -0.009 -0.026 -0.019 0.015
Happy Miss -0.043 -0.080 -0.002 -0.041 -0.040 -0.083 -0.004 -0.049 -0.013 -0.055 -0.050
Neut Hit Angry Hit Happy Hit Neut CR Angry CR Happy CR Neut FA Angry FA Happy FA Neut Miss Angry Miss Happy Miss

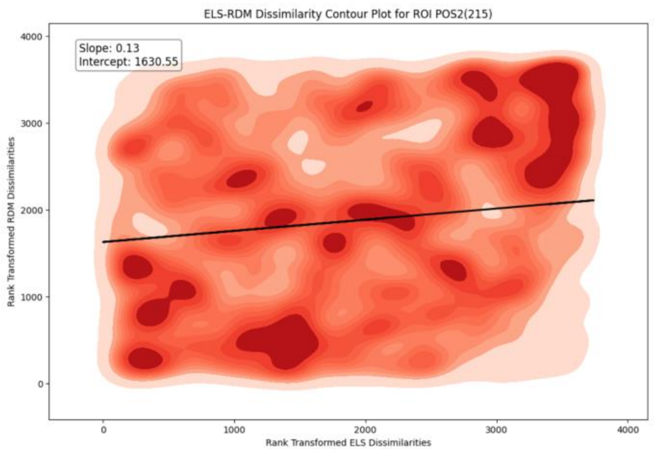
ELS-RDM Dissimilarity Contour Plot for ROI POS2(215)
Slope: 0.13
Intercept: 1630.55
Rank Transformed RDM Dissimilarities
Rank Transformed ELS Dissimilarities

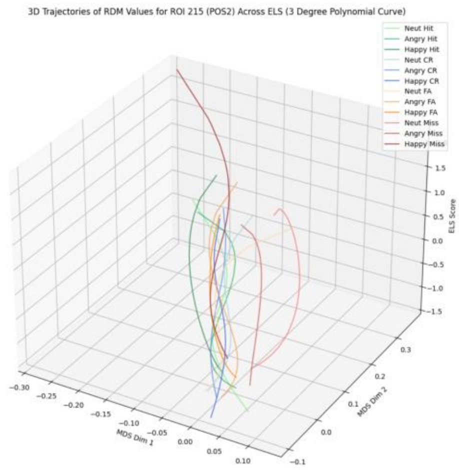
3D Trajectories of RDM Values for ROI 215 (POS2) Across ELS (3 Degree Polynomial Curve)
Neut Hit
Angry Hit
Happy Hit
Neut CR
Angry CR
Happy CR
Neut FA
Angry FA
Happy FA
Neut Miss
Angry Miss
Happy Miss
ELS Score
MDS Dim 1
MDS Dim 2

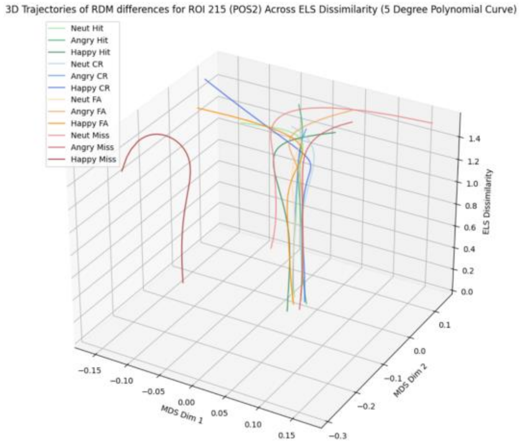
3D Trajectories of RDM differences for ROI 215 (POS2) Across ELS Dissimilarity (5 Degree Polynomial Curve)
Neut Hit
Angry Hit
Happy Hit
Neut CR
Angry CR
Happy CR
Neut FA
Angry FA
Happy FA
Neut Miss
Angry Miss
Happy Miss
ELS Dissimilarity
MDS Dim 1
MDS Dim 2

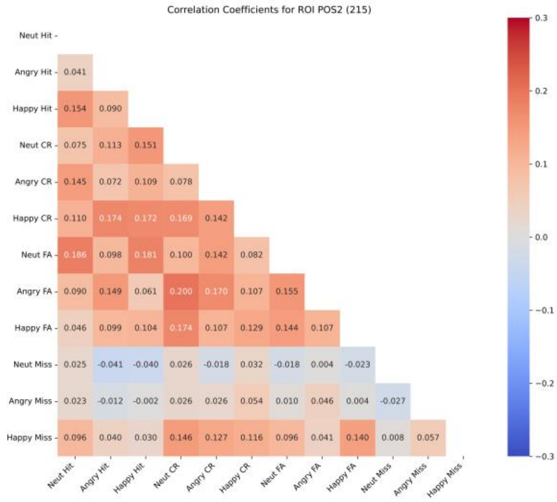
Correlation Coefficients for ROI POS2 (215)
Neut Hit
Angry Hit 0.041
Happy Hit 0.154 0.090
Neut CR 0.075 0.113 0.151
Angry CR 0.145 0.072 0.109 0.078
Happy CR 0.110 0.174 0.172 0.169 0.142
Neut FA 0.186 0.098 0.181 0.100 0.142 0.082
Angry FA 0.090 0.149 0.061 0.200 0.170 0.107 0.155
Happy FA 0.046 0.099 0.104 0.174 0.107 0.129 0.144 0.107
Neut Miss 0.025 -0.041 -0.040 0.026 -0.018 0.032 -0.018 0.004 -0.023
Angry Miss 0.023 -0.012 -0.002 0.026 0.026 0.054 0.010 0.046 0.004 -0.027
Happy Miss 0.096 0.040 0.030 0.146 0.127 0.116 0.096 0.041 0.140 0.008 0.057
Neut Hit Angry Hit Happy Hit Neut CR Angry CR Happy CR Neut FA Angry FA Happy FA Neut Miss Angry Miss Happy Miss

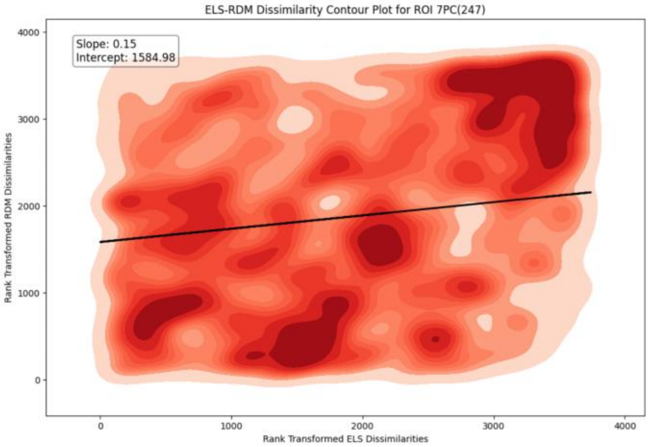
ELS-RDM Dissimilarity Contour Plot for ROI 7PC(247)
Slope: 0.15
Intercept: 1584.98
Rank Transformed RDM Dissimilarities
Rank Transformed ELS Dissimilarities
0
1000
2000
3000
4000

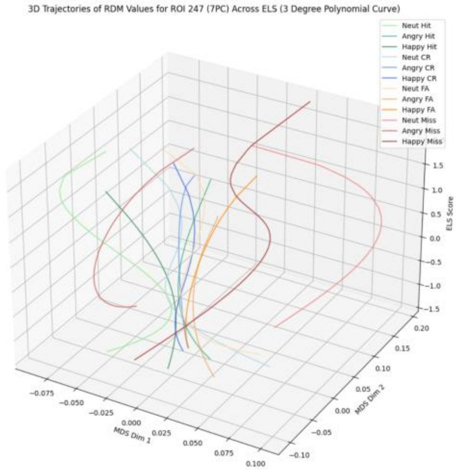
3D Trajectories of RDM Values for ROI 247 (7PC) Across ELS (3 Degree Polynomial Curve)
Neut Hit
Angry Hit
Happy Hit
Neut CR
Angry CR
Happy CR
Neut FA
Angry FA
Happy FA
Neut Miss
Angry Miss
Happy Miss
MDS Dim 1
MDS Dim 2
ELS Score

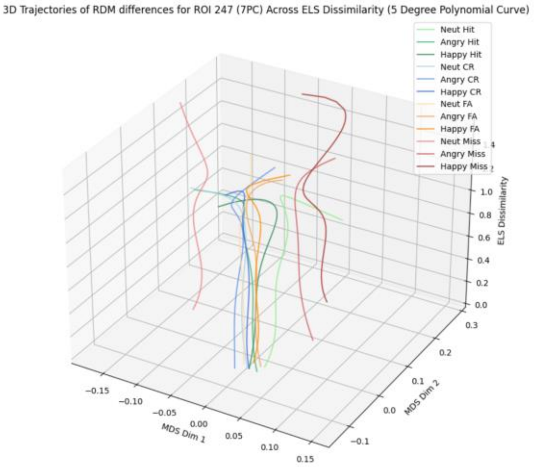
3D Trajectories of RDM differences for ROI 247 (7PC) Across ELS Dissimilarity (5 Degree Polynomial Curve)
Neut Hit
Angry Hit
Happy Hit
Neut CR
Angry CR
Happy CR
Neut FA
Angry FA
Happy FA
Neut Miss
Angry Miss
Happy Miss
MDS Dim 1
MDS Dim 2
ELS Dissimilarity

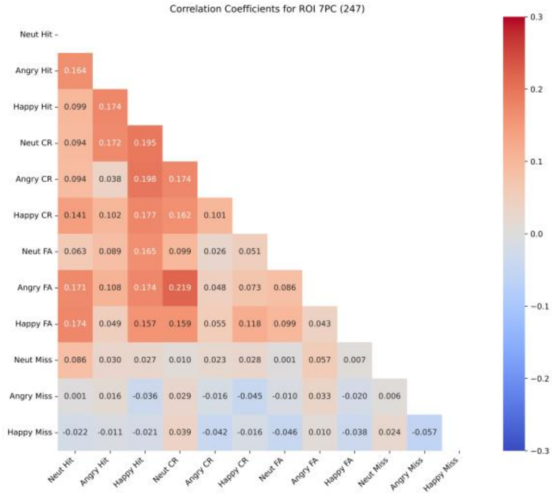
Correlation Coefficients for ROI 7PC (247)
Neut Hit
Angry Hit 0.164
Happy Hit 0.099 0.174
Neut CR 0.094 0.172 0.195
Angry CR 0.094 0.038 0.198 0.174
Happy CR 0.141 0.102 0.177 0.162 0.101
Neut FA 0.063 0.089 0.165 0.099 0.026 0.051
Angry FA 0.171 0.108 0.174 0.219 0.048 0.073 0.086
Happy FA 0.174 0.049 0.157 0.159 0.055 0.118 0.099 0.043
Neut Miss 0.086 0.030 0.027 0.010 0.023 0.028 0.001 0.057 0.007
Angry Miss 0.001 0.016 -0.036 0.029 -0.016 -0.045 -0.010 0.033 -0.020 0.006
Happy Miss -0.022 -0.011 -0.021 0.039 -0.042 -0.016 -0.046 0.010 -0.038 0.024 -0.057
Neut Hit Angry Hit Happy Hit Neut CR Angry CR Happy CR Neut FA Angry FA Happy FA Neut Miss Angry Miss Happy Miss

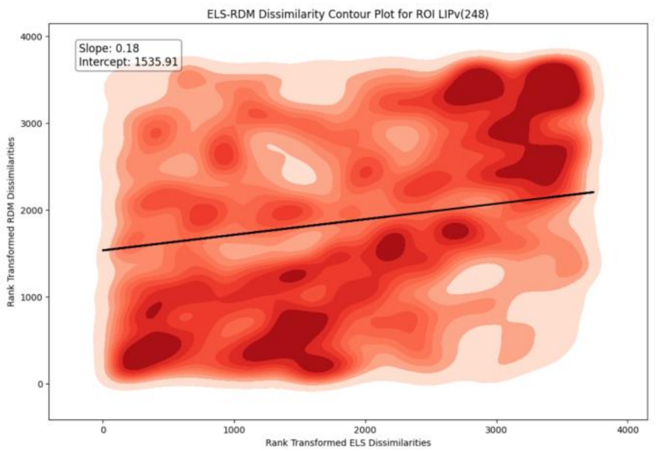
ELS-RDM Dissimilarity Contour Plot for ROI LIPv(248)
Slope: 0.18
Intercept: 1535.91
Rank Transformed RDM Dissimilarities
Rank Transformed ELS Dissimilarities
0
1000
2000
3000
4000

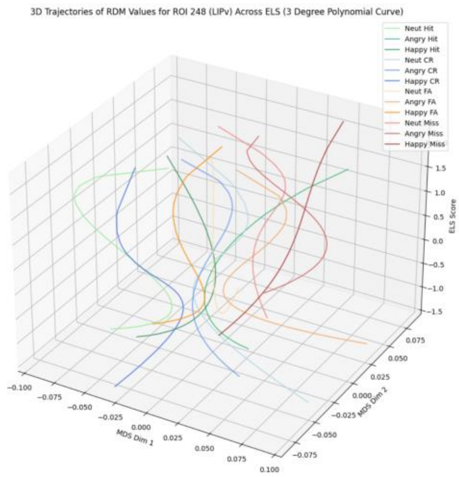
3D Trajectories of RDM Values for ROI 248 (LIPv) Across ELS (3 Degree Polynomial Curve)
Neut Hit
Angry Hit
Happy Hit
Neut CR
Angry CR
Happy CR
Neut FA
Angry FA
Happy FA
Neut Miss
Angry Miss
Happy Miss
ELS Score
MDS Dim 1
MDS Dim 2

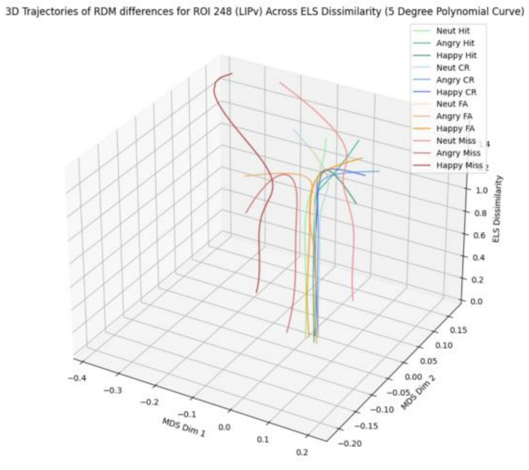
3D Trajectories of RDM differences for ROI 248 (LIPv) Across ELS Dissimilarity (5 Degree Polynomial Curve)
Neut Hit
Angry Hit
Happy Hit
Neut CR
Angry CR
Happy CR
Neut FA
Angry FA
Happy FA
Neut Miss
Angry Miss
Happy Miss
ELS Dissimilarity
MDS Dim 1
MDS Dim 2

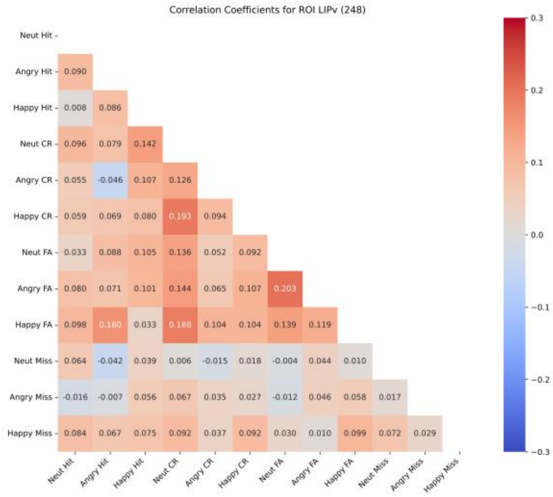
Correlation Coefficients for ROI LIPv (248)
Neut Hit
Angry Hit 0.090
Happy Hit 0.008 0.086
Neut CR 0.096 0.079 0.142
Angry CR 0.055 -0.046 0.107 0.126
Happy CR 0.059 0.069 0.080 0.193 0.094
Neut FA 0.033 0.088 0.105 0.136 0.052 0.092
Angry FA 0.080 0.071 0.101 0.144 0.065 0.107 0.203
Happy FA 0.098 0.160 0.033 0.188 0.104 0.104 0.139 0.119
Neut Miss 0.064 -0.042 0.039 0.006 -0.015 0.018 -0.004 0.044 0.010
Angry Miss -0.016 -0.007 0.056 0.067 0.035 0.027 -0.012 0.046 0.058 0.017
Happy Miss 0.084 0.067 0.075 0.092 0.037 0.092 0.030 0.010 0.099 0.072 0.029
Neut Hit Angry Hit Happy Hit Neut CR Angry CR Happy CR Neut FA Angry FA Happy FA Neut Miss Angry Miss Happy Miss
0.3
0.2
0.1
0.0
−0.1
−0.2
−0.3

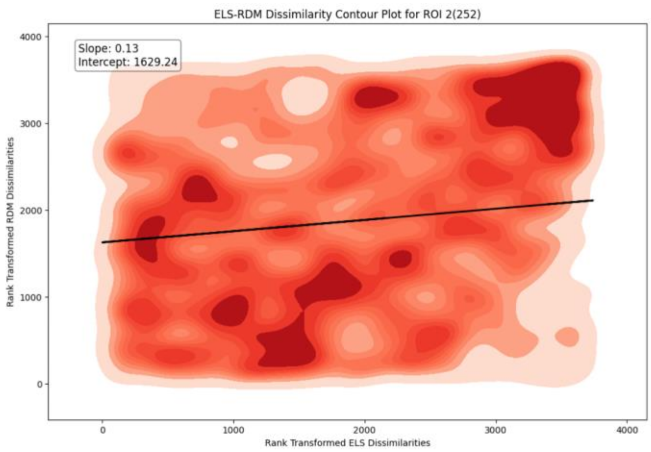
ELS-RDM Dissimilarity Contour Plot for ROI 2(252)
Slope: 0.13
Intercept: 1629.24
Rank Transformed RDM Dissimilarities
Rank Transformed ELS Dissimilarities

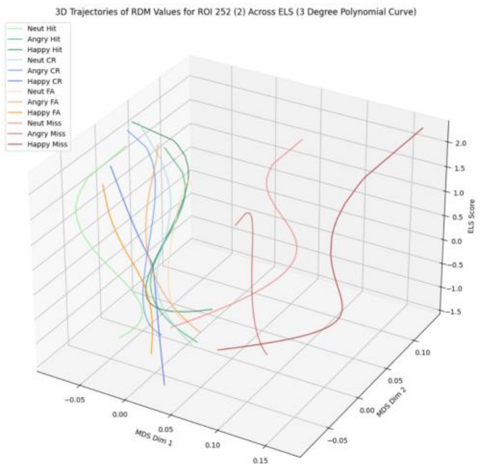
3D Trajectories of RDM Values for ROI 252 (2) Across ELS (3 Degree Polynomial Curve)
Neut Hit
Angry Hit
Happy Hit
Neut CR
Angry CR
Happy CR
Neut FA
Angry FA
Happy FA
Neut Miss
Angry Miss
Happy Miss
MDS Dim 1
MDS Dim 2
ELS Score

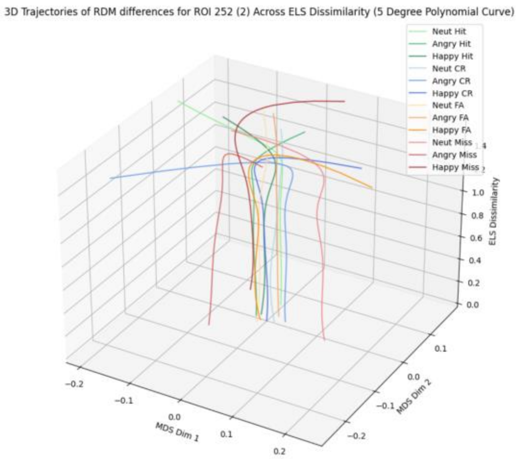
3D Trajectories of RDM differences for ROI 252 (2) Across ELS Dissimilarity (5 Degree Polynomial Curve)
Neut Hit
Angry Hit
Happy Hit
Neut CR
Angry CR
Happy CR
Neut FA
Angry FA
Happy FA
Neut Miss
Angry Miss
Happy Miss
MDS Dim 1
MDS Dim 2
ELS Dissimilarity

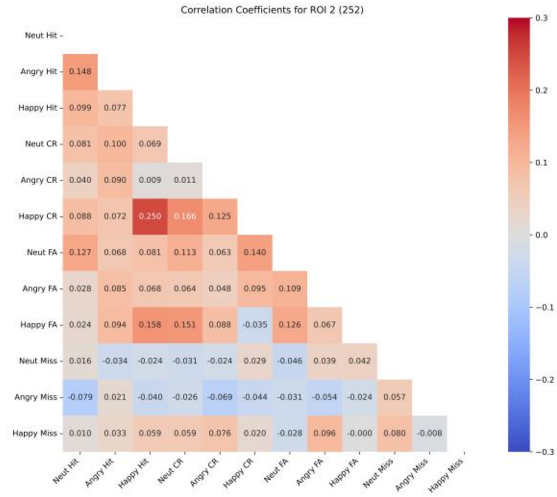
Correlation Coefficients for ROI 2 (252)
Neut Hit
Angry Hit 0.148
Happy Hit 0.099 0.077
Neut CR 0.081 0.100 0.069
Angry CR 0.040 0.090 0.009 0.011
Happy CR 0.088 0.072 0.250 0.166 0.125
Neut FA 0.127 0.068 0.081 0.113 0.063 0.140
Angry FA 0.028 0.085 0.068 0.064 0.048 0.095 0.109
Happy FA 0.024 0.094 0.158 0.151 0.088 -0.035 0.126 0.067
Neut Miss 0.016 -0.034 -0.024 -0.031 -0.024 0.029 -0.046 0.039 0.042
Angry Miss -0.079 0.021 -0.040 -0.026 -0.069 -0.044 -0.031 -0.054 -0.024 0.057
Happy Miss 0.010 0.033 0.059 0.059 0.076 0.020 -0.028 0.096 -0.000 0.080 -0.008
Neut Hit
Angry Hit
Happy Hit
Neut CR
Angry CR
Happy CR
Neut FA
Angry FA
Happy FA
Neut Miss
Angry Miss
Happy Miss

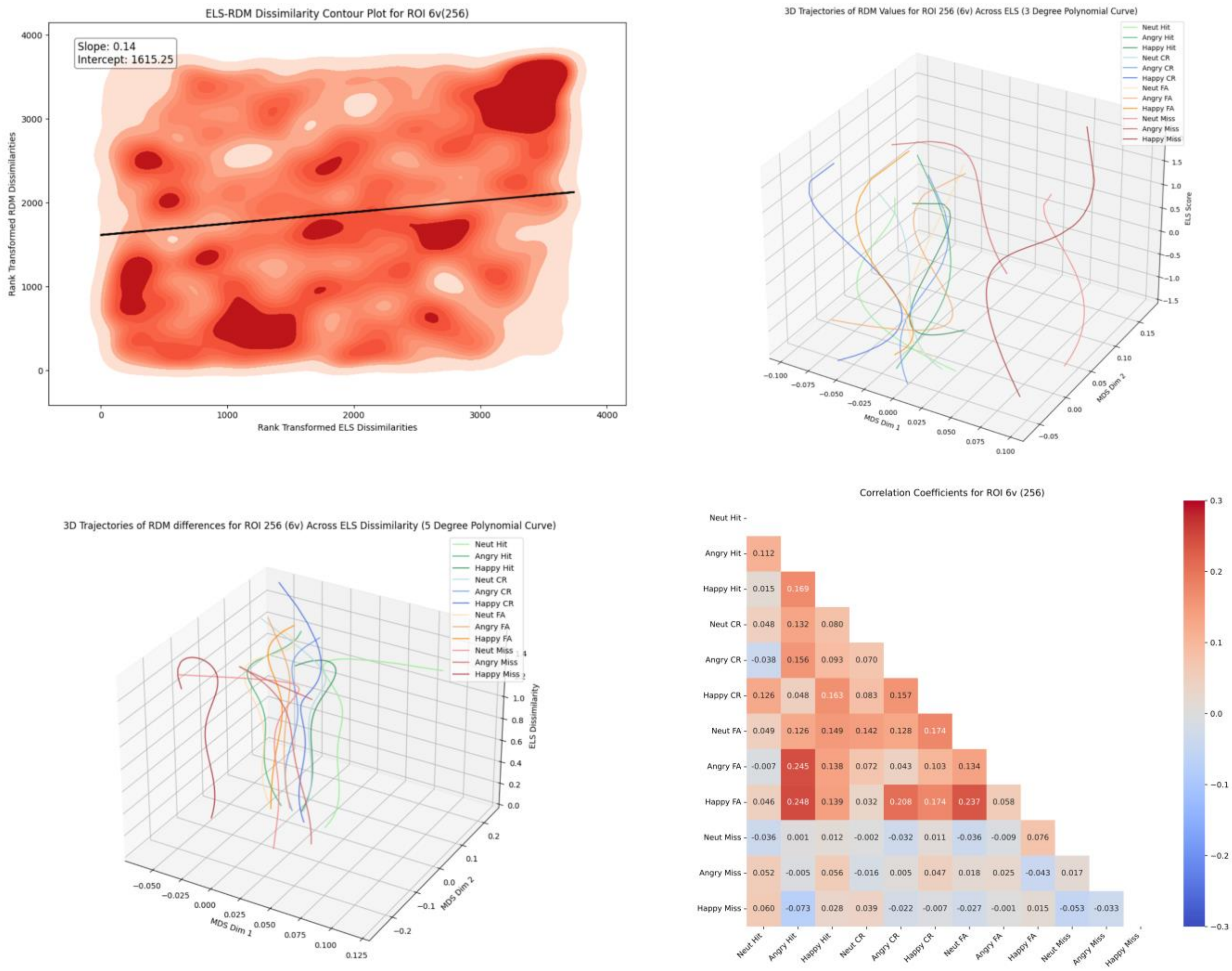
ELS-RDM Dissimilarity Contour Plot for ROI 6v(256)
Slope: 0.14
Intercept: 1615.25
Rank Transformed RDM Dissimilarities
Rank Transformed ELS Dissimilarities
3D Trajectories of RDM Values for ROI 256 (6v) Across ELS (3 Degree Polynomial Curve)
MDS Dim 1
MDS Dim 2
ELS Score
Neut Hit
Angry Hit
Happy Hit
Neut CR
Angry CR
Happy CR
Neut FA
Angry FA
Happy FA
Neut Miss
Angry Miss
Happy Miss
3D Trajectories of RDM differences for ROI 256 (6v) Across ELS Dissimilarity (5 Degree Polynomial Curve)
ELS Dissimilarity
Correlation Coefficients for ROI 6v (256)
Angry Hit 0.112
Happy Hit 0.015 0.169
Neut CR 0.048 0.132 0.080
Angry CR -0.038 0.156 0.093 0.070
Happy CR 0.126 0.048 0.163 0.083 0.157
Neut FA 0.049 0.126 0.149 0.142 0.128 0.174
Angry FA -0.007 0.245 0.138 0.072 0.043 0.103 0.134
Happy FA 0.046 0.248 0.139 0.032 0.208 0.174 0.237 0.058
Neut Miss -0.036 0.001 0.012 -0.002 -0.032 0.011 -0.036 -0.009 0.076
Angry Miss 0.052 -0.005 0.056 -0.016 0.005 0.047 0.018 0.025 -0.043 0.017
Happy Miss 0.060 -0.073 0.028 0.039 -0.022 -0.007 -0.027 -0.001 0.015 -0.053 -0.033

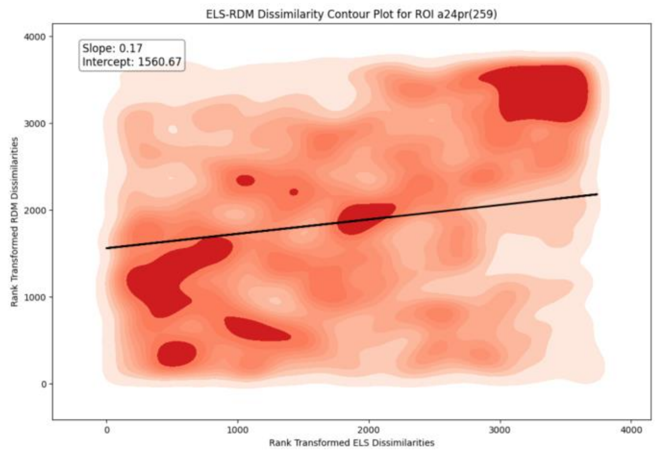
ELS-RDM Dissimilarity Contour Plot for ROI a24pr(259)
Slope: 0.17
Intercept: 1560.67
Rank Transformed RDM Dissimilarities
Rank Transformed ELS Dissimilarities

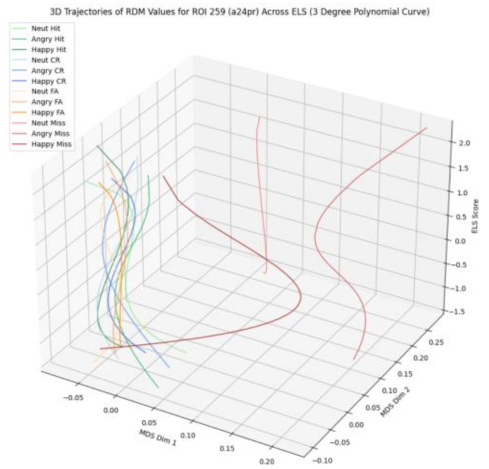
3D Trajectories of RDM Values for ROI 259 (a24pr) Across ELS (3 Degree Polynomial Curve)
Neut Hit
Angry Hit
Happy Hit
Neut CR
Angry CR
Happy CR
Neut FA
Angry FA
Happy FA
Neut Miss
Angry Miss
Happy Miss
ELS Score
MDS Dim 1
MDS Dim 2

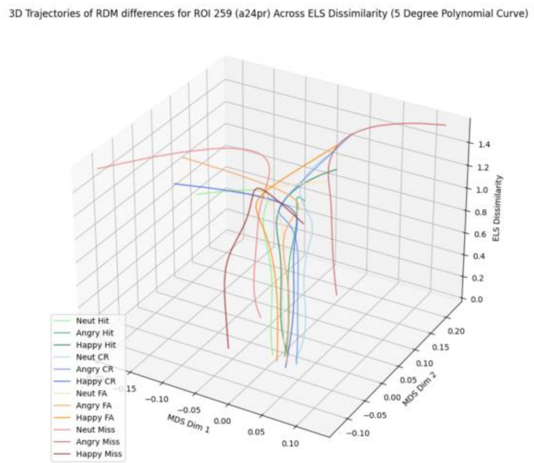
3D Trajectories of RDM differences for ROI 259 (a24pr) Across ELS Dissimilarity (5 Degree Polynomial Curve)
ELS Dissimilarity
MDS Dim 1
MDS Dim 2
Neut Hit
Angry Hit
Happy Hit
Neut CR
Angry CR
Happy CR
Neut FA
Angry FA
Happy FA
Neut Miss
Angry Miss
Happy Miss

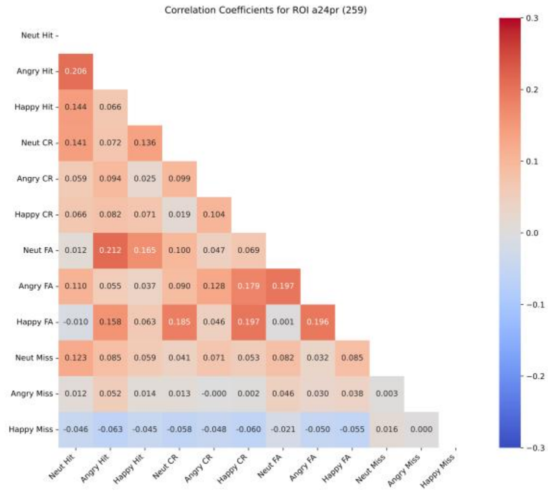
Correlation Coefficients for ROI a24pr (259)
Neut Hit
Angry Hit 0.206
Happy Hit 0.144 0.066
Neut CR 0.141 0.072 0.136
Angry CR 0.059 0.094 0.025 0.099
Happy CR 0.066 0.082 0.071 0.019 0.104
Neut FA 0.012 0.212 0.165 0.100 0.047 0.069
Angry FA 0.110 0.055 0.037 0.090 0.128 0.179 0.197
Happy FA -0.010 0.158 0.063 0.185 0.046 0.197 0.001 0.196
Neut Miss 0.123 0.085 0.059 0.041 0.071 0.053 0.082 0.032 0.085
Angry Miss 0.012 0.052 0.014 0.013 -0.000 0.002 0.046 0.030 0.038 0.003
Happy Miss -0.046 -0.063 -0.045 -0.058 -0.048 -0.060 -0.021 -0.050 -0.055 0.016 0.000
Neut Hit Angry Hit Happy Hit Neut CR Angry CR Happy CR Neut FA Angry FA Happy FA Neut Miss Angry Miss Happy Miss

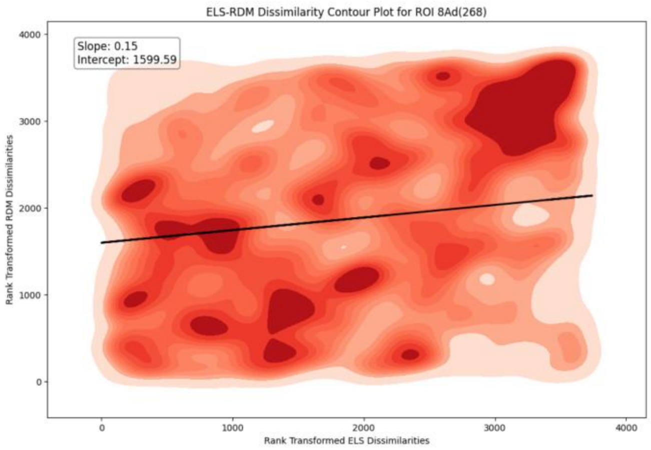
ELS-RDM Dissimilarity Contour Plot for ROI 8Ad(268)
Slope: 0.15
Intercept: 1599.59
Rank Transformed RDM Dissimilarities
Rank Transformed ELS Dissimilarities
0
1000
2000
3000
4000

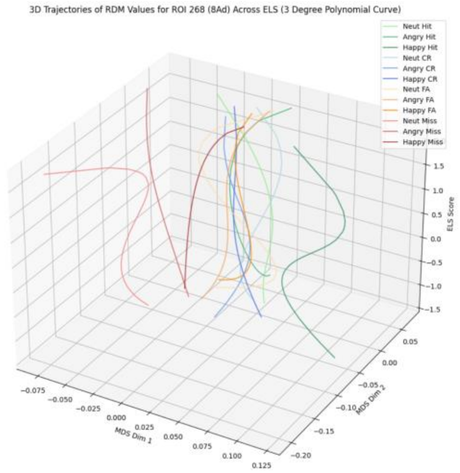
3D Trajectories of RDM Values for ROI 268 (8Ad) Across ELS (3 Degree Polynomial Curve)
Neut Hit
Angry Hit
Happy Hit
Neut CR
Angry CR
Happy CR
Neut FA
Angry FA
Happy FA
Neut Miss
Angry Miss
Happy Miss
ELS Score
MDS Dim 1
MDS Dim 2

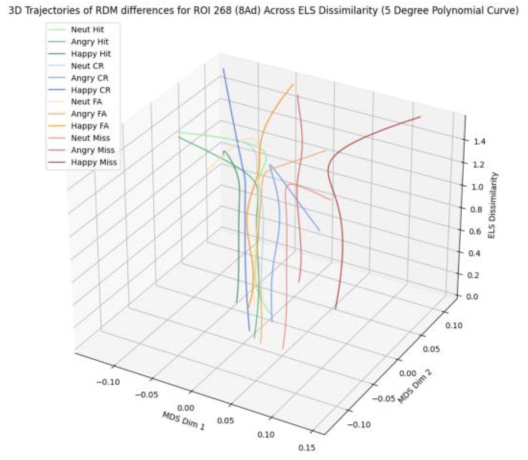
3D Trajectories of RDM differences for ROI 268 (8Ad) Across ELS Dissimilarity (5 Degree Polynomial Curve)
Neut Hit
Angry Hit
Happy Hit
Neut CR
Angry CR
Happy CR
Neut FA
Angry FA
Happy FA
Neut Miss
Angry Miss
Happy Miss
ELS Dissimilarity
MDS Dim 1
MDS Dim 2

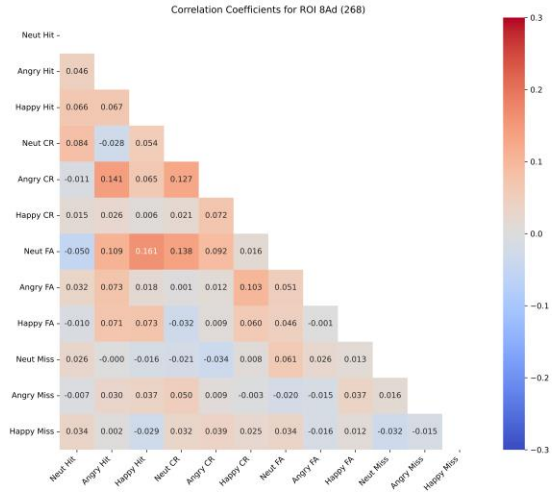
Correlation Coefficients for ROI 8Ad (268)
Neut Hit
Angry Hit 0.046
Happy Hit 0.066 0.067
Neut CR 0.084 -0.028 0.054
Angry CR -0.011 0.141 0.065 0.127
Happy CR 0.015 0.026 0.006 0.021 0.072
Neut FA -0.050 0.109 0.161 0.138 0.092 0.016
Angry FA 0.032 0.073 0.018 0.001 0.012 0.103 0.051
Happy FA -0.010 0.071 0.073 -0.032 0.009 0.060 0.046 -0.001
Neut Miss 0.026 -0.000 -0.016 -0.021 -0.034 0.008 0.061 0.026 0.013
Angry Miss -0.007 0.030 0.037 0.050 0.009 -0.003 -0.020 -0.015 0.037 0.016
Happy Miss 0.034 0.002 -0.029 0.032 0.039 0.025 0.034 -0.016 0.012 -0.032 -0.015
Neut Hit Angry Hit Happy Hit Neut CR Angry CR Happy CR Neut FA Angry FA Happy FA Neut Miss Angry Miss Happy Miss
0.3
0.2
0.1
0.0
−0.1
−0.2
−0.3

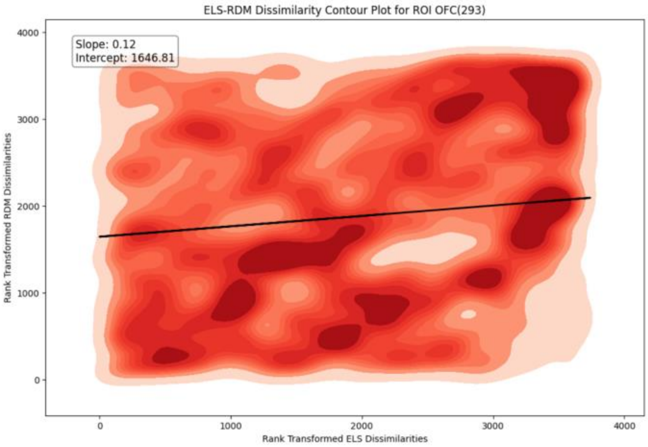
ELS-RDM Dissimilarity Contour Plot for ROI OFC(293)
Slope: 0.12
Intercept: 1646.81
Rank Transformed RDM Dissimilarities
Rank Transformed ELS Dissimilarities

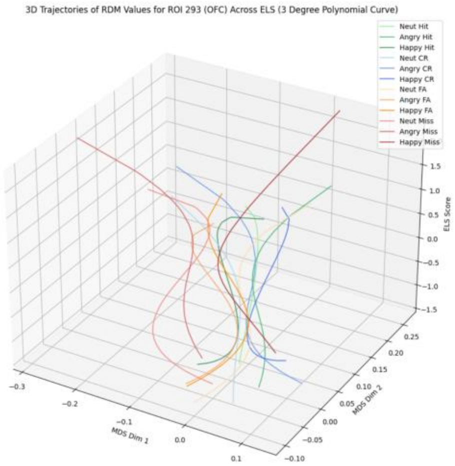
3D Trajectories of RDM Values for ROI 293 (OFC) Across ELS (3 Degree Polynomial Curve)
Neut Hit
Angry Hit
Happy Hit
Neut CR
Angry CR
Happy CR
Neut FA
Angry FA
Happy FA
Neut Miss
Angry Miss
Happy Miss
ELS Score
MDS Dim 1
MDS Dim 2

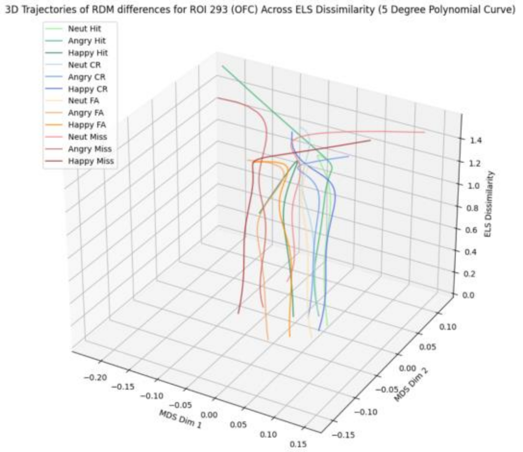
3D Trajectories of RDM differences for ROI 293 (OFC) Across ELS Dissimilarity (5 Degree Polynomial Curve)
Neut Hit
Angry Hit
Happy Hit
Neut CR
Angry CR
Happy CR
Neut FA
Angry FA
Happy FA
Neut Miss
Angry Miss
Happy Miss
ELS Dissimilarity
MDS Dim 1
MDS Dim 2

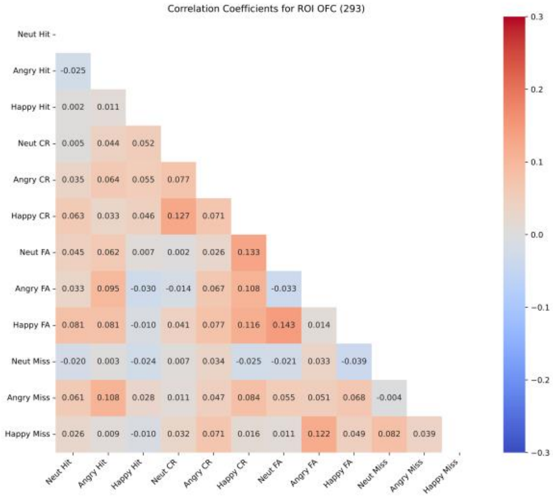
Correlation Coefficients for ROI OFC (293)
Neut Hit
Angry Hit -0.025
Happy Hit 0.002 0.011
Neut CR 0.005 0.044 0.052
Angry CR 0.035 0.064 0.055 0.077
Happy CR 0.063 0.033 0.046 0.127 0.071
Neut FA 0.045 0.062 0.007 0.002 0.026 0.133
Angry FA 0.033 0.095 -0.030 -0.014 0.067 0.108 -0.033
Happy FA 0.081 0.081 -0.010 0.041 0.077 0.116 0.143 0.014
Neut Miss -0.020 0.003 -0.024 0.007 0.034 -0.025 -0.021 0.033 -0.039
Angry Miss 0.061 0.108 0.028 0.011 0.047 0.084 0.055 0.051 0.068 -0.004
Happy Miss 0.026 0.009 -0.010 0.032 0.071 0.016 0.011 0.122 0.049 0.082 0.039
Neut Hit Angry Hit Happy Hit Neut CR Angry CR Happy CR Neut FA Angry FA Happy FA Neut Miss Angry Miss Happy Miss

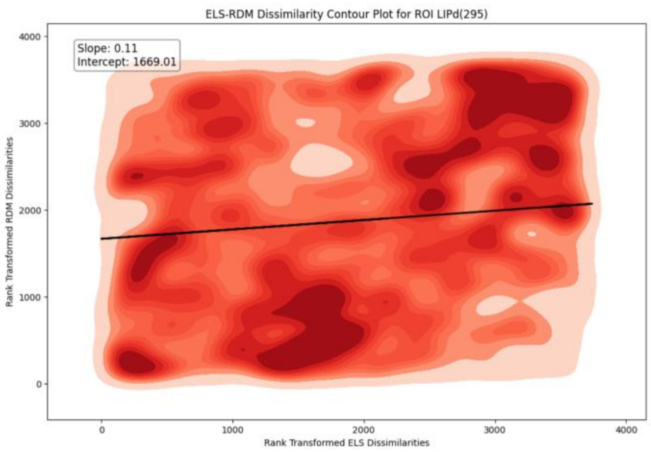
ELS-RDM Dissimilarity Contour Plot for ROI LIPd(295)
Slope: 0.11
Intercept: 1669.01
Rank Transformed RDM Dissimilarities
Rank Transformed ELS Dissimilarities

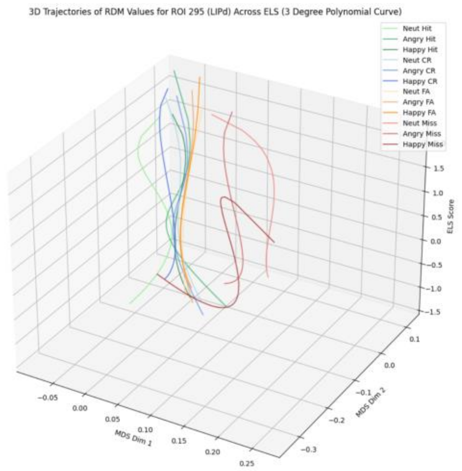
3D Trajectories of RDM Values for ROI 295 (LIPd) Across ELS (3 Degree Polynomial Curve)
Neut Hit
Angry Hit
Happy Hit
Neut CR
Angry CR
Happy CR
Neut FA
Angry FA
Happy FA
Neut Miss
Angry Miss
Happy Miss
ELS Score
MDS Dim 1
MDS Dim 2

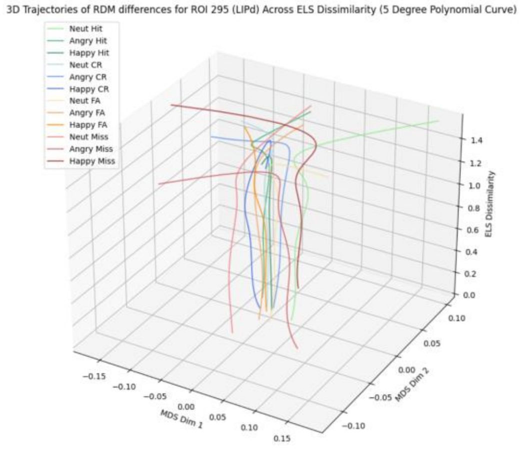
3D Trajectories of RDM differences for ROI 295 (LIPd) Across ELS Dissimilarity (5 Degree Polynomial Curve)
Neut Hit
Angry Hit
Happy Hit
Neut CR
Angry CR
Happy CR
Neut FA
Angry FA
Happy FA
Neut Miss
Angry Miss
Happy Miss
ELS Dissimilarity
MDS Dim 1
MDS Dim 2

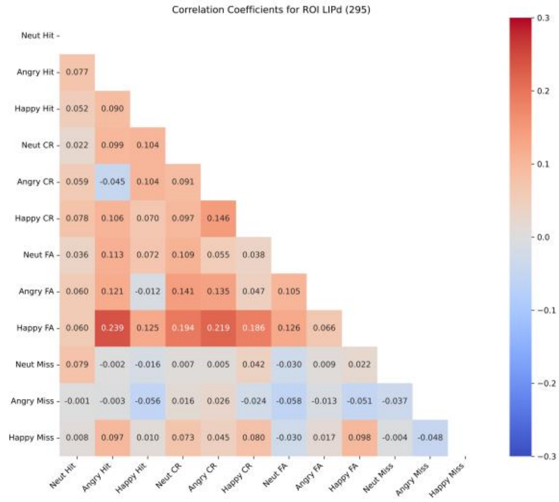
Correlation Coefficients for ROI LIPd (295)
Neut Hit
Angry Hit 0.077
Happy Hit 0.052 0.090
Neut CR 0.022 0.099 0.104
Angry CR 0.059 -0.045 0.104 0.091
Happy CR 0.078 0.106 0.070 0.097 0.146
Neut FA 0.036 0.113 0.072 0.109 0.055 0.038
Angry FA 0.060 0.121 -0.012 0.141 0.135 0.047 0.105
Happy FA 0.060 0.239 0.125 0.194 0.219 0.186 0.126 0.066
Neut Miss 0.079 -0.002 -0.016 0.007 0.005 0.042 -0.030 0.009 0.022
Angry Miss -0.001 -0.003 -0.056 0.016 0.026 -0.024 -0.058 -0.013 -0.051 -0.037
Happy Miss 0.008 0.097 0.010 0.073 0.045 0.080 -0.030 0.017 0.098 -0.004 -0.048
Neut Hit Angry Hit Happy Hit Neut CR Angry CR Happy CR Neut FA Angry FA Happy FA Neut Miss Angry Miss Happy Miss

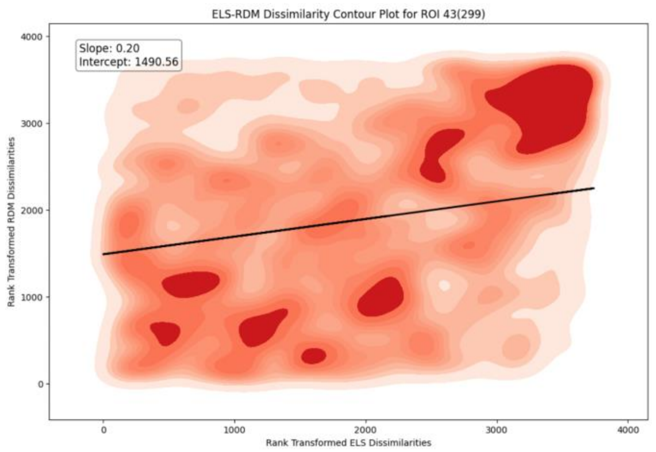
ELS-RDM Dissimilarity Contour Plot for ROI 43(299)
Slope: 0.20
Intercept: 1490.56
Rank Transformed RDM Dissimilarities
Rank Transformed ELS Dissimilarities

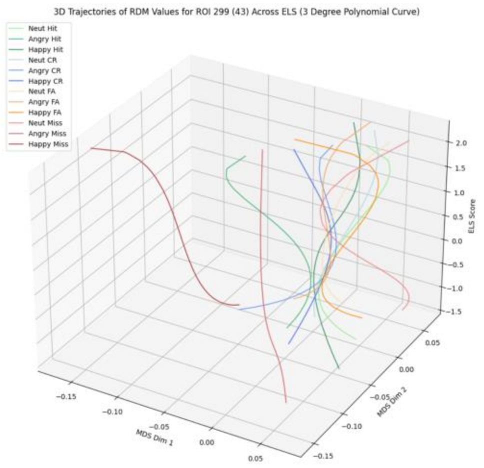
3D Trajectories of RDM Values for ROI 299 (43) Across ELS (3 Degree Polynomial Curve)
Neut Hit
Angry Hit
Happy Hit
Neut CR
Angry CR
Happy CR
Neut FA
Angry FA
Happy FA
Neut Miss
Angry Miss
Happy Miss
ELS Score
MDS Dim 1
MDS Dim 2

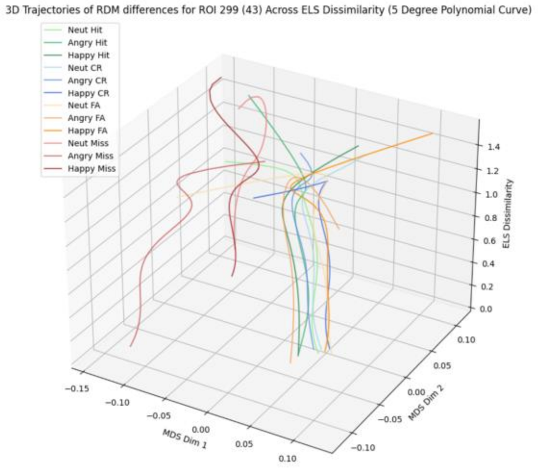
3D Trajectories of RDM differences for ROI 299 (43) Across ELS Dissimilarity (5 Degree Polynomial Curve)
Neut Hit
Angry Hit
Happy Hit
Neut CR
Angry CR
Happy CR
Neut FA
Angry FA
Happy FA
Neut Miss
Angry Miss
Happy Miss
ELS Dissimilarity
MDS Dim 1
MDS Dim 2

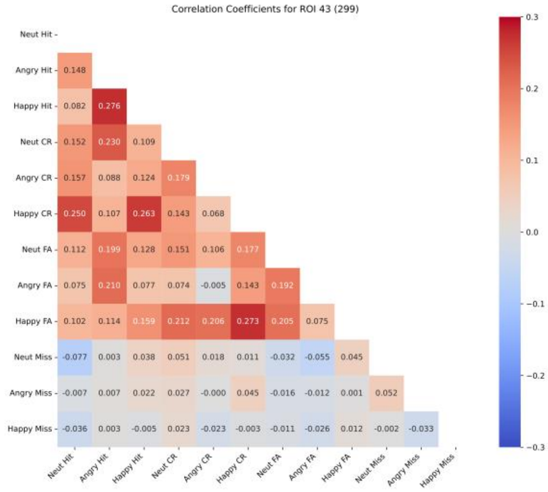
Correlation Coefficients for ROI 43 (299)
Neut Hit
Angry Hit 0.148
Happy Hit 0.082 0.276
Neut CR 0.152 0.230 0.109
Angry CR 0.157 0.088 0.124 0.179
Happy CR 0.250 0.107 0.263 0.143 0.068
Neut FA 0.112 0.199 0.128 0.151 0.106 0.177
Angry FA 0.075 0.210 0.077 0.074 -0.005 0.143 0.192
Happy FA 0.102 0.114 0.159 0.212 0.206 0.273 0.205 0.075
Neut Miss -0.077 0.003 0.038 0.051 0.018 0.011 -0.032 -0.055 0.045
Angry Miss -0.007 0.007 0.022 0.027 -0.000 0.045 -0.016 -0.012 0.001 0.052
Happy Miss -0.036 0.003 -0.005 0.023 -0.023 -0.003 -0.011 -0.026 0.012 -0.002 -0.033
Neut Hit Angry Hit Happy Hit Neut CR Angry CR Happy CR Neut FA Angry FA Happy FA Neut Miss Angry Miss Happy Miss

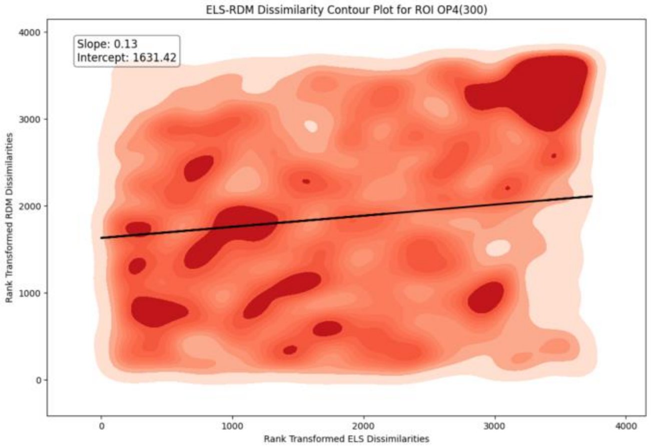
ELS-RDM Dissimilarity Contour Plot for ROI OP4(300)
Slope: 0.13
Intercept: 1631.42
Rank Transformed RDM Dissimilarities
Rank Transformed ELS Dissimilarities
0
1000
2000
3000
4000

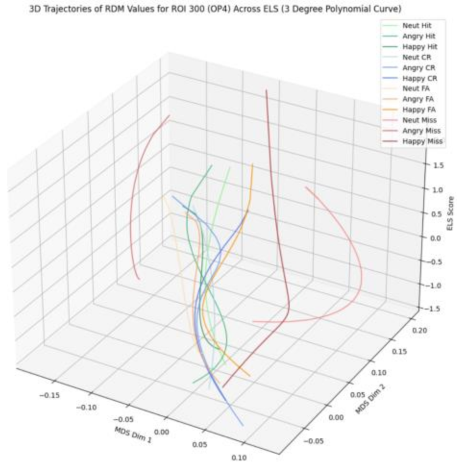
3D Trajectories of RDM Values for ROI 300 (OP4) Across ELS (3 Degree Polynomial Curve)
Neut Hit
Angry Hit
Happy Hit
Neut CR
Angry CR
Happy CR
Neut FA
Angry FA
Happy FA
Neut Miss
Angry Miss
Happy Miss
ELS Score
MDS Dim 1
MDS Dim 2

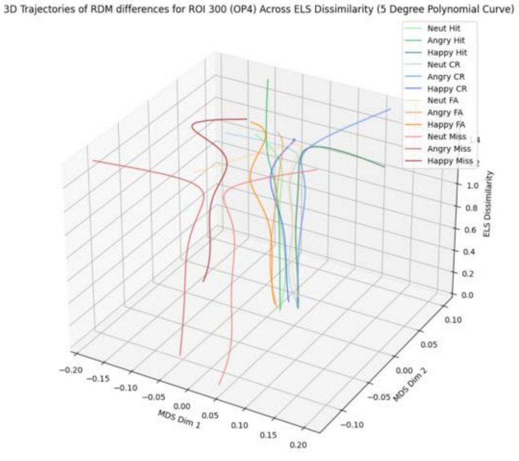
3D Trajectories of RDM differences for ROI 300 (OP4) Across ELS Dissimilarity (5 Degree Polynomial Curve)
Neut Hit
Angry Hit
Happy Hit
Neut CR
Angry CR
Happy CR
Neut FA
Angry FA
Happy FA
Neut Miss
Angry Miss
Happy Miss
ELS Dissimilarity
MDS Dim 1
MDS Dim 2

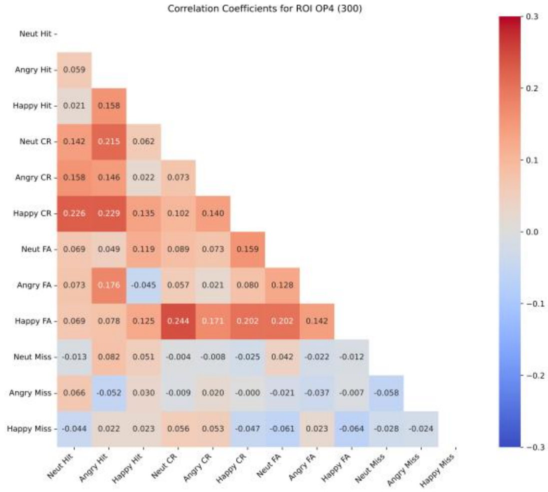
Correlation Coefficients for ROI OP4 (300)
Neut Hit
Angry Hit 0.059
Happy Hit 0.021 0.158
Neut CR 0.142 0.215 0.062
Angry CR 0.158 0.146 0.022 0.073
Happy CR 0.226 0.229 0.135 0.102 0.140
Neut FA 0.069 0.049 0.119 0.089 0.073 0.159
Angry FA 0.073 0.176 -0.045 0.057 0.021 0.080 0.128
Happy FA 0.069 0.078 0.125 0.244 0.171 0.202 0.202 0.142
Neut Miss -0.013 0.082 0.051 -0.004 -0.008 -0.025 0.042 -0.022 -0.012
Angry Miss 0.066 -0.052 0.030 -0.009 0.020 -0.000 -0.021 -0.037 -0.007 -0.058
Happy Miss -0.044 0.022 0.023 0.056 0.053 -0.047 -0.061 0.023 -0.064 -0.028 -0.024
Neut Hit Angry Hit Happy Hit Neut CR Angry CR Happy CR Neut FA Angry FA Happy FA Neut Miss Angry Miss Happy Miss

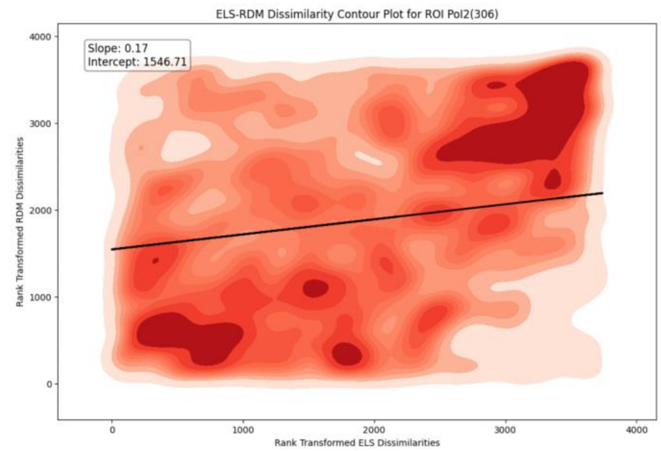
ELS-RDM Dissimilarity Contour Plot for ROI Pol2(306)
Slope: 0.17
Intercept: 1546.71
Rank Transformed RDM Dissimilarities
Rank Transformed ELS Dissimilarities

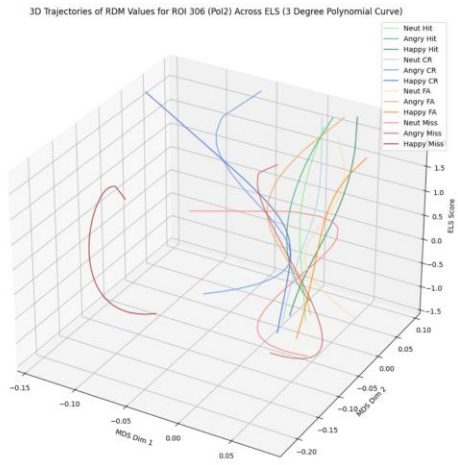
3D Trajectories of RDM Values for ROI 306 (Pol2) Across ELS (3 Degree Polynomial Curve)
Neut Hit
Angry Hit
Happy Hit
Neut CR
Angry CR
Happy CR
Neut FA
Angry FA
Happy FA
Neut Miss
Angry Miss
Happy Miss
ELS Score
MDS Dim 1
MDS Dim 2

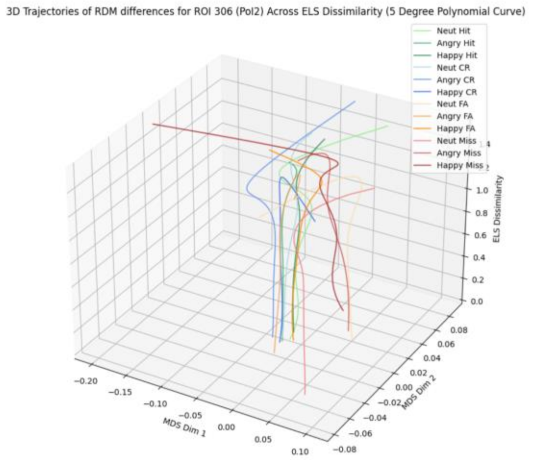
3D Trajectories of RDM differences for ROI 306 (Pol2) Across ELS Dissimilarity (5 Degree Polynomial Curve)
Neut Hit
Angry Hit
Happy Hit
Neut CR
Angry CR
Happy CR
Neut FA
Angry FA
Happy FA
Neut Miss
Angry Miss
Happy Miss
ELS Dissimilarity
MDS Dim 1
MDS Dim 2

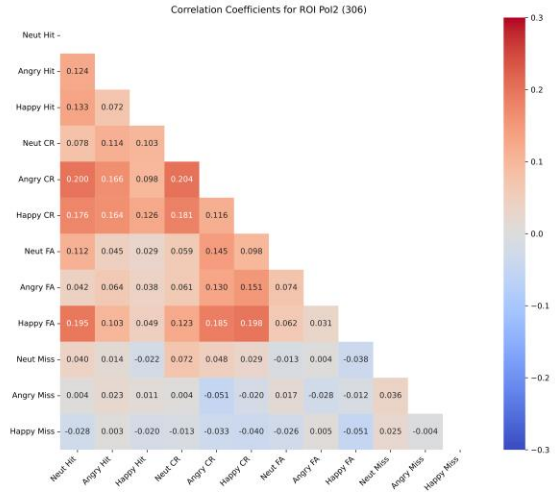
Correlation Coefficients for ROI Pol2 (306)
Neut Hit
Angry Hit 0.124
Happy Hit 0.133 0.072
Neut CR 0.078 0.114 0.103
Angry CR 0.200 0.166 0.098 0.204
Happy CR 0.176 0.164 0.126 0.181 0.116
Neut FA 0.112 0.045 0.029 0.059 0.145 0.098
Angry FA 0.042 0.064 0.038 0.061 0.130 0.151 0.074
Happy FA 0.195 0.103 0.049 0.123 0.185 0.198 0.062 0.031
Neut Miss 0.040 0.014 -0.022 0.072 0.048 0.029 -0.013 0.004 -0.038
Angry Miss 0.004 0.023 0.011 0.004 -0.051 -0.020 0.017 -0.028 -0.012 0.036
Happy Miss -0.028 0.003 -0.020 -0.013 -0.033 -0.040 -0.026 0.005 -0.051 0.025 -0.004
Neut Hit Angry Hit Happy Hit Neut CR Angry CR Happy CR Neut FA Angry FA Happy FA Neut Miss Angry Miss Happy Miss

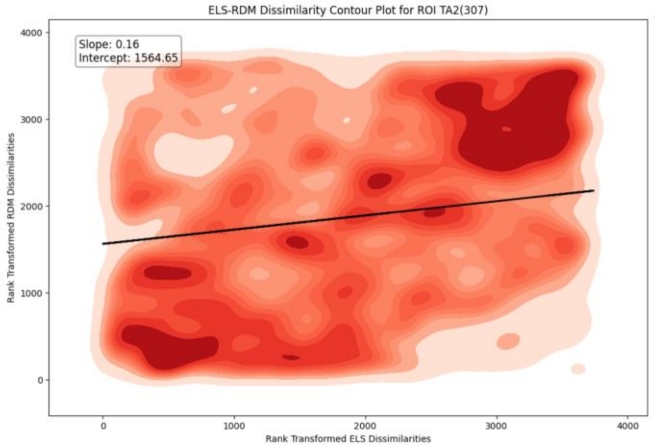
ELS-RDM Dissimilarity Contour Plot for ROI TA2(307)
Slope: 0.16
Intercept: 1564.65
Rank Transformed RDM Dissimilarities
Rank Transformed ELS Dissimilarities

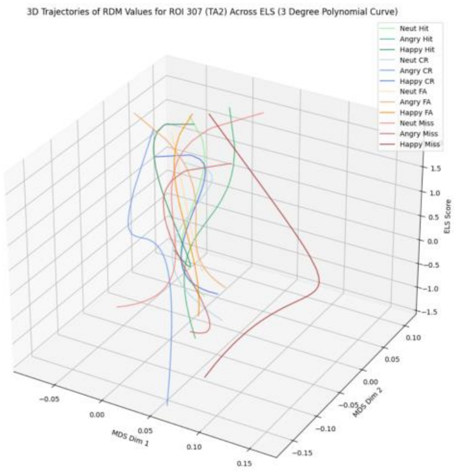
3D Trajectories of RDM Values for ROI 307 (TA2) Across ELS (3 Degree Polynomial Curve)
Neut Hit
Angry Hit
Happy Hit
Neut CR
Angry CR
Happy CR
Neut FA
Angry FA
Happy FA
Neut Miss
Angry Miss
Happy Miss
ELS Score
MDS Dim 1
MDS Dim 2

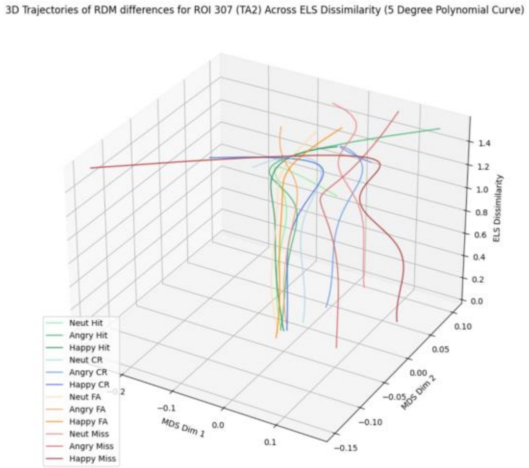
3D Trajectories of RDM differences for ROI 307 (TA2) Across ELS Dissimilarity (5 Degree Polynomial Curve)
ELS Dissimilarity
MDS Dim 1
MDS Dim 2
Neut Hit
Angry Hit
Happy Hit
Neut CR
Angry CR
Happy CR
Neut FA
Angry FA
Happy FA
Neut Miss
Angry Miss
Happy Miss

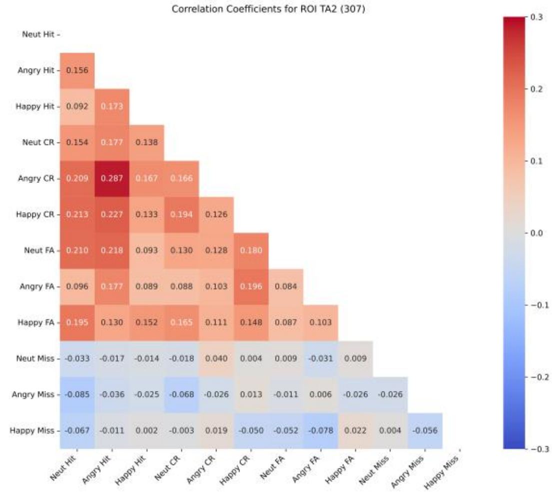
Correlation Coefficients for ROI TA2 (307)
Neut Hit
Angry Hit 0.156
Happy Hit 0.092 0.173
Neut CR 0.154 0.177 0.138
Angry CR 0.209 0.287 0.167 0.166
Happy CR 0.213 0.227 0.133 0.194 0.126
Neut FA 0.210 0.218 0.093 0.130 0.128 0.180
Angry FA 0.096 0.177 0.089 0.088 0.103 0.196 0.084
Happy FA 0.195 0.130 0.152 0.165 0.111 0.148 0.087 0.103
Neut Miss -0.033 -0.017 -0.014 -0.018 0.040 0.004 0.009 -0.031 0.009
Angry Miss -0.085 -0.036 -0.025 -0.068 -0.026 0.013 -0.011 0.006 -0.026 -0.026
Happy Miss -0.067 -0.011 0.002 -0.003 0.019 -0.050 -0.052 -0.078 0.022 0.004 -0.056
Neut Hit, Angry Hit, Happy Hit, Neut CR, Angry CR, Happy CR, Neut FA, Angry FA, Happy FA, Neut Miss, Angry Miss, Happy Miss

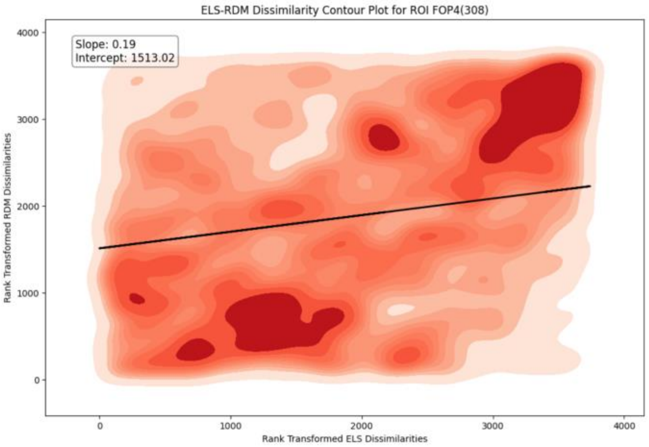
ELS-RDM Dissimilarity Contour Plot for ROI FOP4(308)
Slope: 0.19
Intercept: 1513.02
Rank Transformed RDM Dissimilarities
Rank Transformed ELS Dissimilarities
0
1000
2000
3000
4000

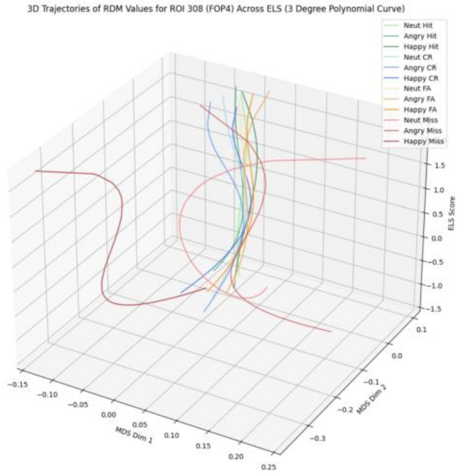
3D Trajectories of RDM Values for ROI 308 (FOP4) Across ELS (3 Degree Polynomial Curve)
Neut Hit
Angry Hit
Happy Hit
Neut CR
Angry CR
Happy CR
Neut FA
Angry FA
Happy FA
Neut Miss
Angry Miss
Happy Miss
ELS Score
MDS Dim 1
MDS Dim 2

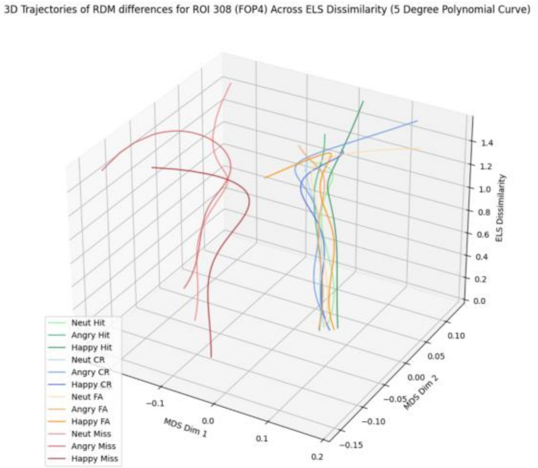
3D Trajectories of RDM differences for ROI 308 (FOP4) Across ELS Dissimilarity (5 Degree Polynomial Curve)
ELS Dissimilarity
MDS Dim 1
MDS Dim 2
Neut Hit
Angry Hit
Happy Hit
Neut CR
Angry CR
Happy CR
Neut FA
Angry FA
Happy FA
Neut Miss
Angry Miss
Happy Miss

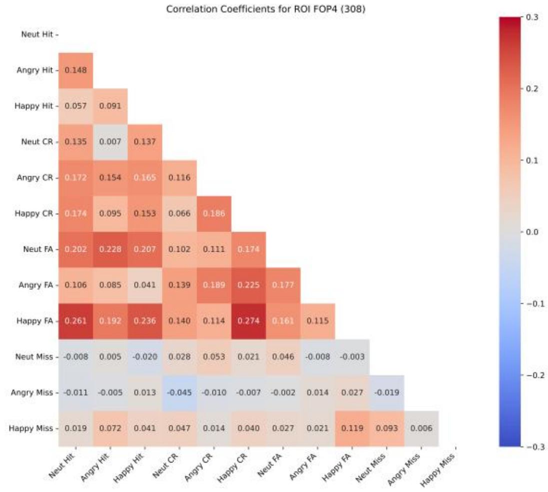
Correlation Coefficients for ROI FOP4 (308)
Neut Hit
Angry Hit 0.148
Happy Hit 0.057 0.091
Neut CR 0.135 0.007 0.137
Angry CR 0.172 0.154 0.165 0.116
Happy CR 0.174 0.095 0.153 0.066 0.186
Neut FA 0.202 0.228 0.207 0.102 0.111 0.174
Angry FA 0.106 0.085 0.041 0.139 0.189 0.225 0.177
Happy FA 0.261 0.192 0.236 0.140 0.114 0.274 0.161 0.115
Neut Miss -0.008 0.005 -0.020 0.028 0.053 0.021 0.046 -0.008 -0.003
Angry Miss -0.011 -0.005 0.013 -0.045 -0.010 -0.007 -0.002 0.014 0.027 -0.019
Happy Miss 0.019 0.072 0.041 0.047 0.014 0.040 0.027 0.021 0.119 0.093 0.006
Neut Hit Angry Hit Happy Hit Neut CR Angry CR Happy CR Neut FA Angry FA Happy FA Neut Miss Angry Miss Happy Miss

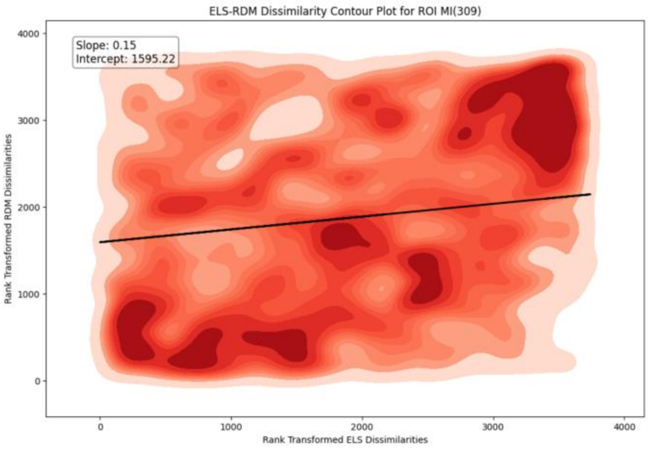
ELS-RDM Dissimilarity Contour Plot for ROI MI(309)
Slope: 0.15
Intercept: 1595.22
Rank Transformed RDM Dissimilarities
Rank Transformed ELS Dissimilarities

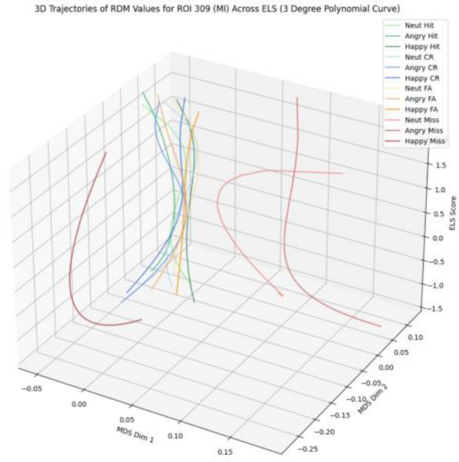
3D Trajectories of RDM Values for ROI 309 (MI) Across ELS (3 Degree Polynomial Curve)
Neut Hit
Angry Hit
Happy Hit
Neut CR
Angry CR
Happy CR
Neut FA
Angry FA
Happy FA
Neut Miss
Angry Miss
Happy Miss
ELS Score
MDS Dim 1
MDS Dim 2

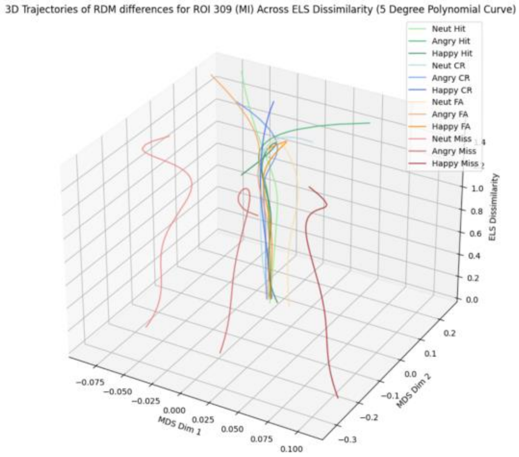
3D Trajectories of RDM differences for ROI 309 (MI) Across ELS Dissimilarity (5 Degree Polynomial Curve)
Neut Hit
Angry Hit
Happy Hit
Neut CR
Angry CR
Happy CR
Neut FA
Angry FA
Happy FA
Neut Miss
Angry Miss
Happy Miss
ELS Dissimilarity
MDS Dim 1
MDS Dim 2

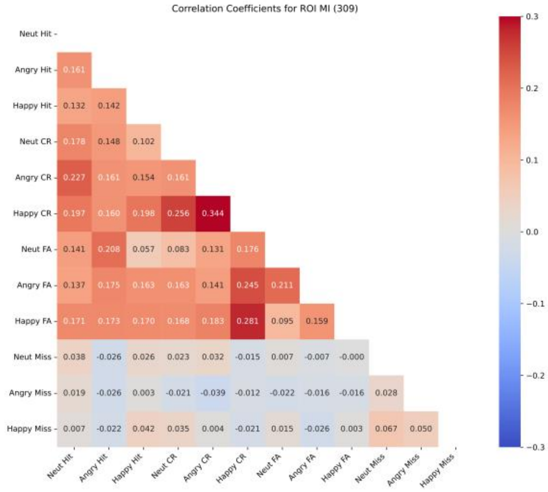
Correlation Coefficients for ROI MI (309)
Neut Hit
Angry Hit 0.161
Happy Hit 0.132 0.142
Neut CR 0.178 0.148 0.102
Angry CR 0.227 0.161 0.154 0.161
Happy CR 0.197 0.160 0.198 0.256 0.344
Neut FA 0.141 0.208 0.057 0.083 0.131 0.176
Angry FA 0.137 0.175 0.163 0.163 0.141 0.245 0.211
Happy FA 0.171 0.173 0.170 0.168 0.183 0.281 0.095 0.159
Neut Miss 0.038 -0.026 0.026 0.023 0.032 -0.015 0.007 -0.007 -0.000
Angry Miss 0.019 -0.026 0.003 -0.021 -0.039 -0.012 -0.022 -0.016 -0.016 0.028
Happy Miss 0.007 -0.022 0.042 0.035 0.004 -0.021 0.015 -0.026 0.003 0.067 0.050
Neut Hit Angry Hit Happy Hit Neut CR Angry CR Happy CR Neut FA Angry FA Happy FA Neut Miss Angry Miss Happy Miss

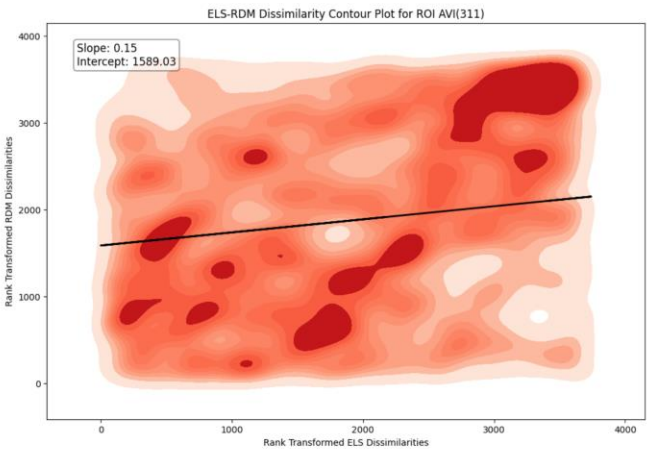
ELS-RDM Dissimilarity Contour Plot for ROI AVI(311)
Slope: 0.15
Intercept: 1589.03
Rank Transformed RDM Dissimilarities
Rank Transformed ELS Dissimilarities

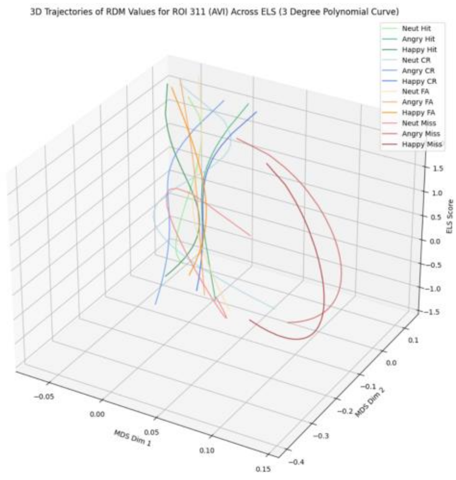
3D Trajectories of RDM Values for ROI 311 (AVI) Across ELS (3 Degree Polynomial Curve)
Neut Hit
Angry Hit
Happy Hit
Neut CR
Angry CR
Happy CR
Neut FA
Angry FA
Happy FA
Neut Miss
Angry Miss
Happy Miss
ELS Score
MDS Dim 1
MDS Dim 2

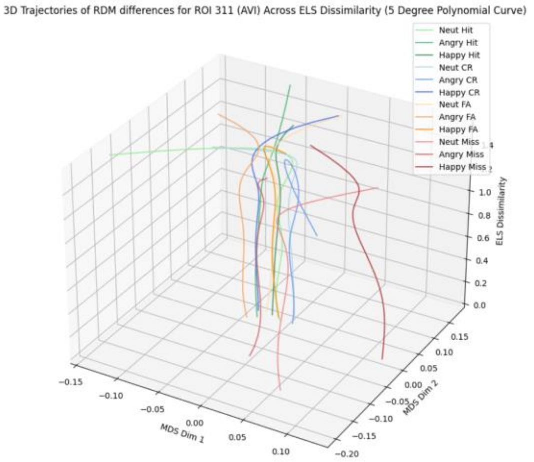
3D Trajectories of RDM differences for ROI 311 (AVI) Across ELS Dissimilarity (5 Degree Polynomial Curve)
Neut Hit
Angry Hit
Happy Hit
Neut CR
Angry CR
Happy CR
Neut FA
Angry FA
Happy FA
Neut Miss
Angry Miss
Happy Miss
ELS Dissimilarity
MDS Dim 1
MDS Dim 2

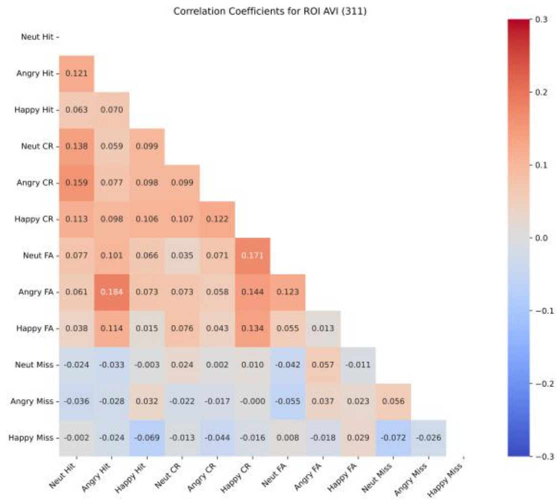
Correlation Coefficients for ROI AVI (311)
Neut Hit
Angry Hit 0.121
Happy Hit 0.063 0.070
Neut CR 0.138 0.059 0.099
Angry CR 0.159 0.077 0.098 0.099
Happy CR 0.113 0.098 0.106 0.107 0.122
Neut FA 0.077 0.101 0.066 0.035 0.071 0.171
Angry FA 0.061 0.184 0.073 0.073 0.058 0.144 0.123
Happy FA 0.038 0.114 0.015 0.076 0.043 0.134 0.055 0.013
Neut Miss -0.024 -0.033 -0.003 0.024 0.002 0.010 -0.042 0.057 -0.011
Angry Miss -0.036 -0.028 0.032 -0.022 -0.017 -0.000 -0.055 0.037 0.023 0.056
Happy Miss -0.002 -0.024 -0.069 -0.013 -0.044 -0.016 0.008 -0.018 0.029 -0.072 -0.026
Neut Hit Angry Hit Happy Hit Neut CR Angry CR Happy CR Neut FA Angry FA Happy FA Neut Miss Angry Miss Happy Miss

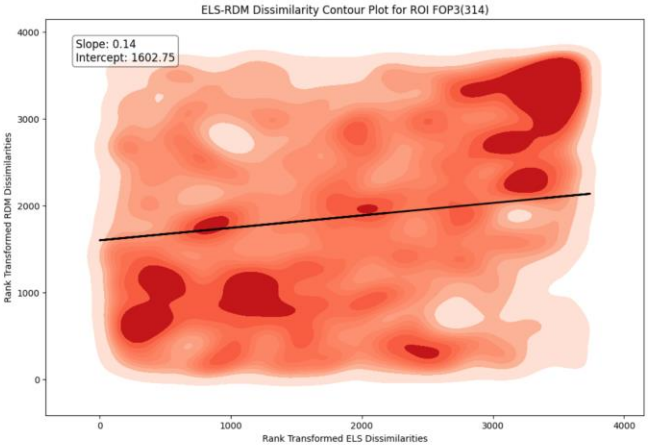
ELS-RDM Dissimilarity Contour Plot for ROI FOP3(314)
Slope: 0.14
Intercept: 1602.75
Rank Transformed RDM Dissimilarities
Rank Transformed ELS Dissimilarities

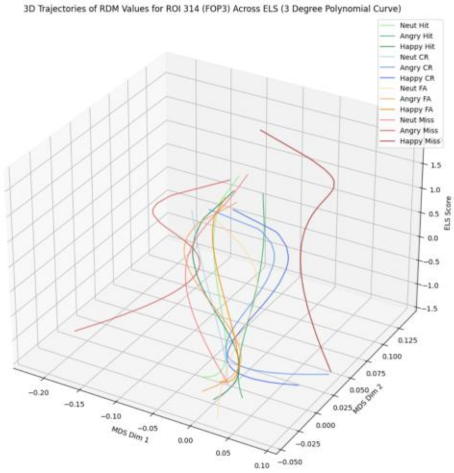
3D Trajectories of RDM Values for ROI 314 (FOP3) Across ELS (3 Degree Polynomial Curve)
Neut Hit
Angry Hit
Happy Hit
Neut CR
Angry CR
Happy CR
Neut FA
Angry FA
Happy FA
Neut Miss
Angry Miss
Happy Miss
MDS Dim 1
MDS Dim 2
ELS Score

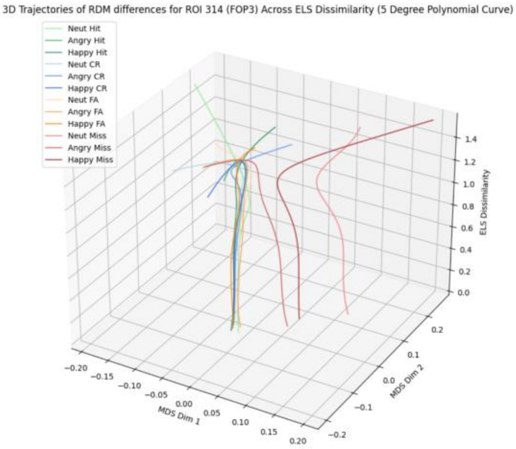
3D Trajectories of RDM differences for ROI 314 (FOP3) Across ELS Dissimilarity (5 Degree Polynomial Curve)
Neut Hit
Angry Hit
Happy Hit
Neut CR
Angry CR
Happy CR
Neut FA
Angry FA
Happy FA
Neut Miss
Angry Miss
Happy Miss
MDS Dim 1
MDS Dim 2
ELS Dissimilarity

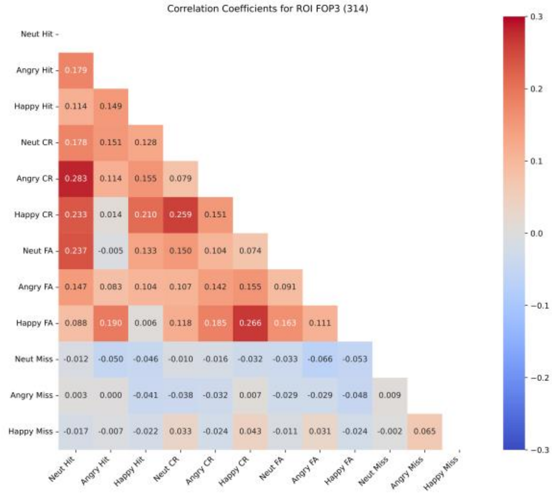
Correlation Coefficients for ROI FOP3 (314)
Neut Hit
Angry Hit 0.179
Happy Hit 0.114 0.149
Neut CR 0.178 0.151 0.128
Angry CR 0.283 0.114 0.155 0.079
Happy CR 0.233 0.014 0.210 0.259 0.151
Neut FA 0.237 -0.005 0.133 0.150 0.104 0.074
Angry FA 0.147 0.083 0.104 0.107 0.142 0.155 0.091
Happy FA 0.088 0.190 0.006 0.118 0.185 0.266 0.163 0.111
Neut Miss -0.012 -0.050 -0.046 -0.010 -0.016 -0.032 -0.033 -0.066 -0.053
Angry Miss 0.003 0.000 -0.041 -0.038 -0.032 0.007 -0.029 -0.029 -0.048 0.009
Happy Miss -0.017 -0.007 -0.022 0.033 -0.024 0.043 -0.011 0.031 -0.024 -0.002 0.065
Neut Hit Angry Hit Happy Hit Neut CR Angry CR Happy CR Neut FA Angry FA Happy FA Neut Miss Angry Miss Happy Miss

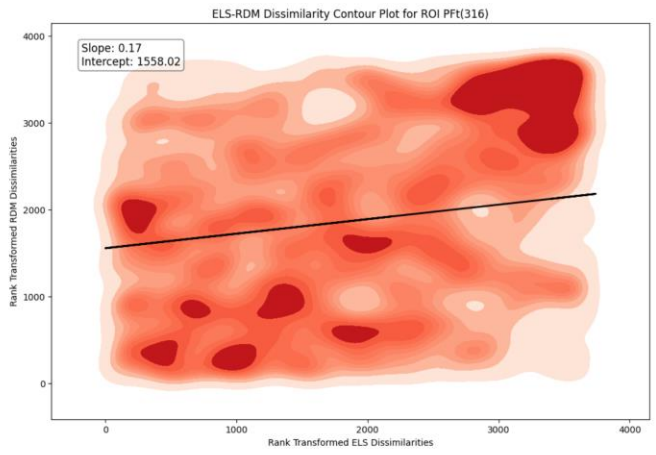
ELS-RDM Dissimilarity Contour Plot for ROI PFt(316)
Slope: 0.17
Intercept: 1558.02
Rank Transformed RDM Dissimilarities
Rank Transformed ELS Dissimilarities
0
1000
2000
3000
4000

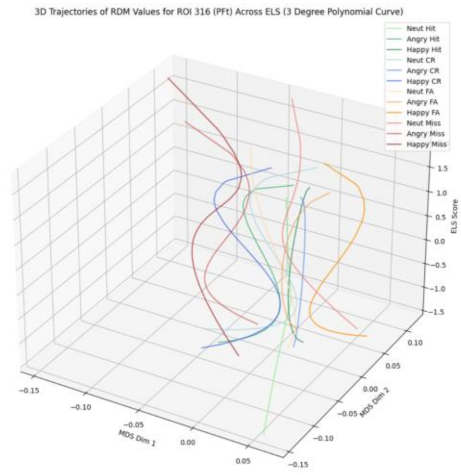
3D Trajectories of RDM Values for ROI 316 (PFt) Across ELS (3 Degree Polynomial Curve)
Neut Hit
Angry Hit
Happy Hit
Neut CR
Angry CR
Happy CR
Neut FA
Angry FA
Happy FA
Neut Miss
Angry Miss
Happy Miss
ELS Score
MDS Dim 1
MDS Dim 2

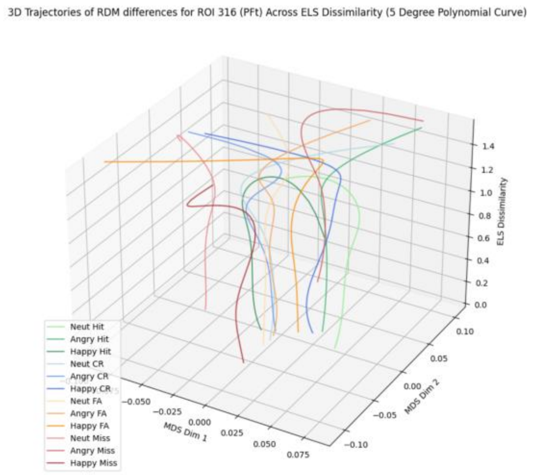
3D Trajectories of RDM differences for ROI 316 (PFt) Across ELS Dissimilarity (5 Degree Polynomial Curve)
ELS Dissimilarity
MDS Dim 1
MDS Dim 2
Neut Hit
Angry Hit
Happy Hit
Neut CR
Angry CR
Happy CR
Neut FA
Angry FA
Happy FA
Neut Miss
Angry Miss
Happy Miss

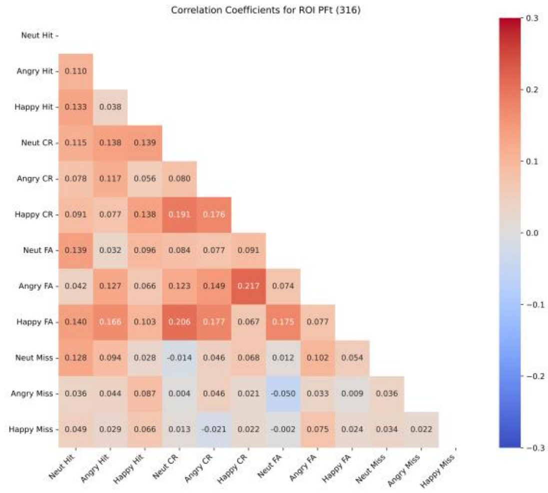
Correlation Coefficients for ROI PFt (316)
Neut Hit
Angry Hit 0.110
Happy Hit 0.133 0.038
Neut CR 0.115 0.138 0.139
Angry CR 0.078 0.117 0.056 0.080
Happy CR 0.091 0.077 0.138 0.191 0.176
Neut FA 0.139 0.032 0.096 0.084 0.077 0.091
Angry FA 0.042 0.127 0.066 0.123 0.149 0.217 0.074
Happy FA 0.140 0.166 0.103 0.206 0.177 0.067 0.175 0.077
Neut Miss 0.128 0.094 0.028 -0.014 0.046 0.068 0.012 0.102 0.054
Angry Miss 0.036 0.044 0.087 0.004 0.046 0.021 -0.050 0.033 0.009 0.036
Happy Miss 0.049 0.029 0.066 0.013 -0.021 0.022 -0.002 0.075 0.024 0.034 0.022
Neut Hit Angry Hit Happy Hit Neut CR Angry CR Happy CR Neut FA Angry FA Happy FA Neut Miss Angry Miss Happy Miss

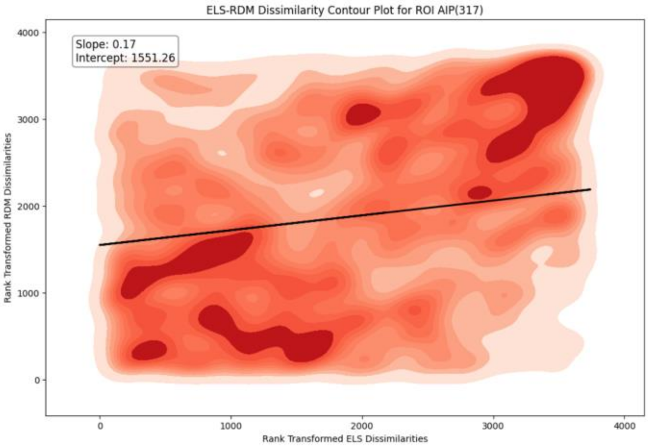
ELS-RDM Dissimilarity Contour Plot for ROI AIP(317)
Slope: 0.17
Intercept: 1551.26
Rank Transformed RDM Dissimilarities
Rank Transformed ELS Dissimilarities

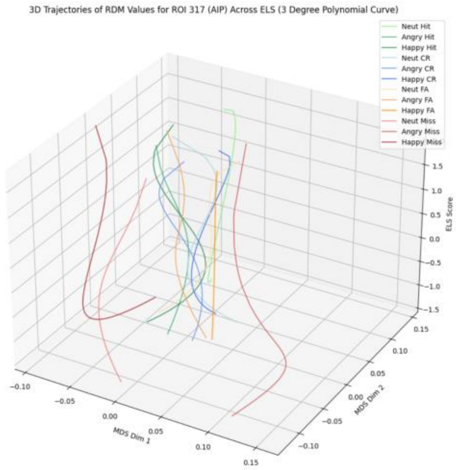
3D Trajectories of RDM Values for ROI 317 (AIP) Across ELS (3 Degree Polynomial Curve)
Neut Hit
Angry Hit
Happy Hit
Neut CR
Angry CR
Happy CR
Neut FA
Angry FA
Happy FA
Neut Miss
Angry Miss
Happy Miss
ELS Score
MDS Dim 1
MDS Dim 2

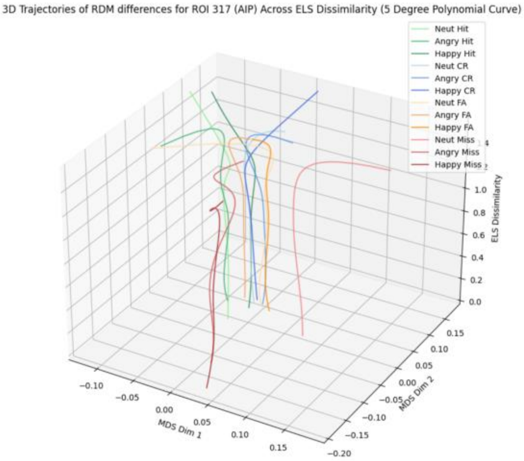
3D Trajectories of RDM differences for ROI 317 (AIP) Across ELS Dissimilarity (5 Degree Polynomial Curve)
Neut Hit
Angry Hit
Happy Hit
Neut CR
Angry CR
Happy CR
Neut FA
Angry FA
Happy FA
Neut Miss
Angry Miss
Happy Miss
ELS Dissimilarity
MDS Dim 1
MDS Dim 2

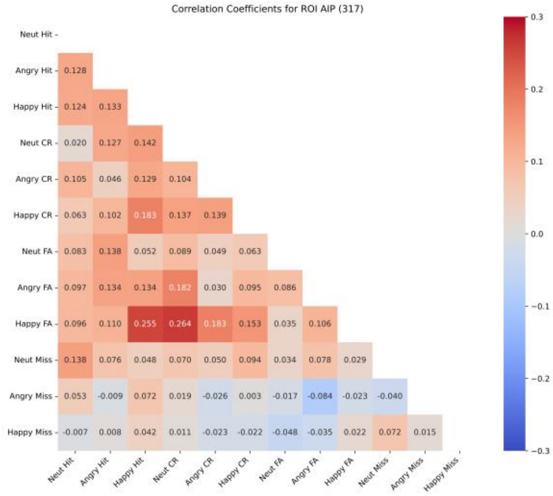
Correlation Coefficients for ROI AIP (317)
Neut Hit
Angry Hit 0.128
Happy Hit 0.124 0.133
Neut CR 0.020 0.127 0.142
Angry CR 0.105 0.046 0.129 0.104
Happy CR 0.063 0.102 0.183 0.137 0.139
Neut FA 0.083 0.138 0.052 0.089 0.049 0.063
Angry FA 0.097 0.134 0.134 0.182 0.030 0.095 0.086
Happy FA 0.096 0.110 0.255 0.264 0.183 0.153 0.035 0.106
Neut Miss 0.138 0.076 0.048 0.070 0.050 0.094 0.034 0.078 0.029
Angry Miss 0.053 -0.009 0.072 0.019 -0.026 0.003 -0.017 -0.084 -0.023 -0.040
Happy Miss -0.007 0.008 0.042 0.011 -0.023 -0.022 -0.048 -0.035 0.022 0.072 0.015
Neut Hit
Angry Hit
Happy Hit
Neut CR
Angry CR
Happy CR
Neut FA
Angry FA
Happy FA
Neut Miss
Angry Miss
Happy Miss

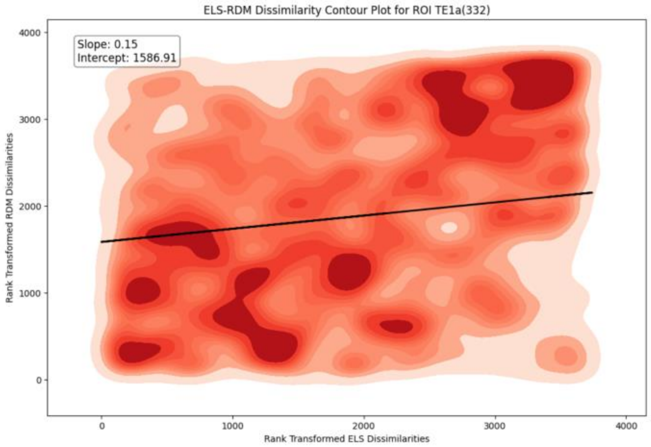
ELS-RDM Dissimilarity Contour Plot for ROI TE1a(332)
Slope: 0.15
Intercept: 1586.91
Rank Transformed RDM Dissimilarities
Rank Transformed ELS Dissimilarities

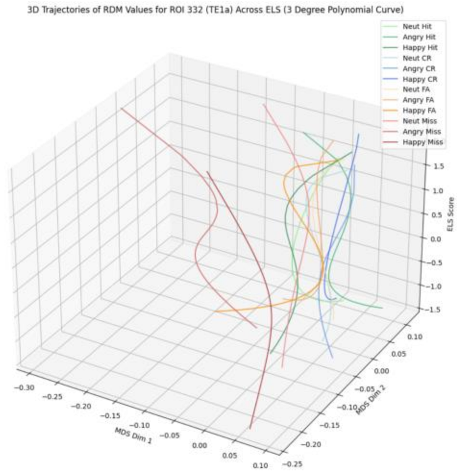
3D Trajectories of RDM Values for ROI 332 (TE1a) Across ELS (3 Degree Polynomial Curve)
Neut Hit
Angry Hit
Happy Hit
Neut CR
Angry CR
Happy CR
Neut FA
Angry FA
Happy FA
Neut Miss
Angry Miss
Happy Miss
MDS Dim 1
MDS Dim 2
ELS Score

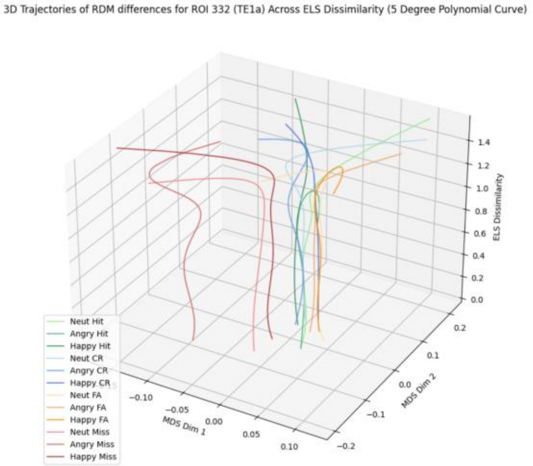
3D Trajectories of RDM differences for ROI 332 (TE1a) Across ELS Dissimilarity (5 Degree Polynomial Curve)
Neut Hit
Angry Hit
Happy Hit
Neut CR
Angry CR
Happy CR
Neut FA
Angry FA
Happy FA
Neut Miss
Angry Miss
Happy Miss
MDS Dim 1
MDS Dim 2
ELS Dissimilarity

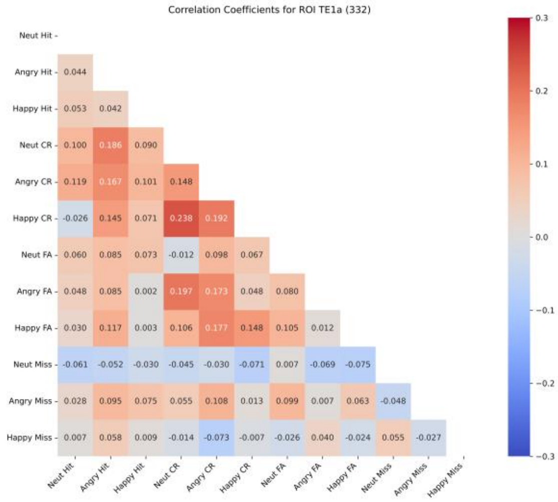
Correlation Coefficients for ROI TE1a (332)
Neut Hit
Angry Hit 0.044
Happy Hit 0.053 0.042
Neut CR 0.100 0.186 0.090
Angry CR 0.119 0.167 0.101 0.148
Happy CR -0.026 0.145 0.071 0.238 0.192
Neut FA 0.060 0.085 0.073 -0.012 0.098 0.067
Angry FA 0.048 0.085 0.002 0.197 0.173 0.048 0.080
Happy FA 0.030 0.117 0.003 0.106 0.177 0.148 0.105 0.012
Neut Miss -0.061 -0.052 -0.030 -0.045 -0.030 -0.071 0.007 -0.069 -0.075
Angry Miss 0.028 0.095 0.075 0.055 0.108 0.013 0.099 0.007 0.063 -0.048
Happy Miss 0.007 0.058 0.009 -0.014 -0.073 -0.007 -0.026 0.040 -0.024 0.055 -0.027
Neut Hit Angry Hit Happy Hit Neut CR Angry CR Happy CR Neut FA Angry FA Happy FA Neut Miss Angry Miss Happy Miss

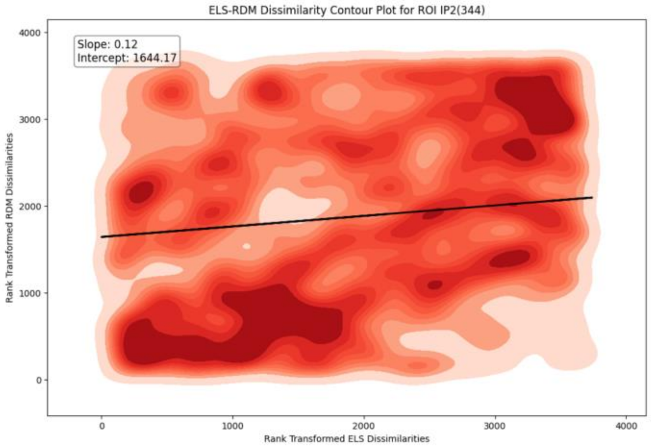
ELS-RDM Dissimilarity Contour Plot for ROI IP2(344)
Slope: 0.12
Intercept: 1644.17
Rank Transformed RDM Dissimilarities
Rank Transformed ELS Dissimilarities
0
1000
2000
3000
4000

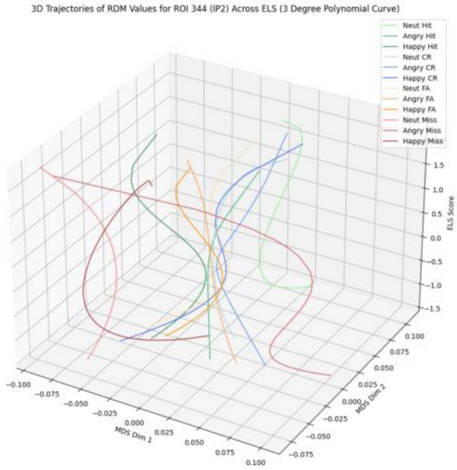
3D Trajectories of RDM Values for ROI 344 (IP2) Across ELS (3 Degree Polynomial Curve)
Neut Hit
Angry Hit
Happy Hit
Neut CR
Angry CR
Happy CR
Neut FA
Angry FA
Happy FA
Neut Miss
Angry Miss
Happy Miss
MDS Dim 1
MDS Dim 2
ELS Score

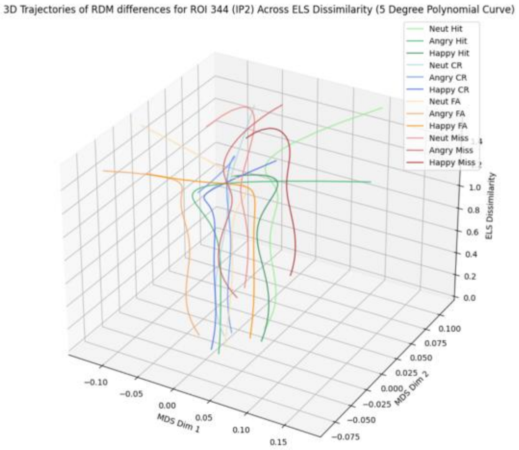
3D Trajectories of RDM differences for ROI 344 (IP2) Across ELS Dissimilarity (5 Degree Polynomial Curve)
Neut Hit
Angry Hit
Happy Hit
Neut CR
Angry CR
Happy CR
Neut FA
Angry FA
Happy FA
Neut Miss
Angry Miss
Happy Miss
MDS Dim 1
MDS Dim 2
ELS Dissimilarity

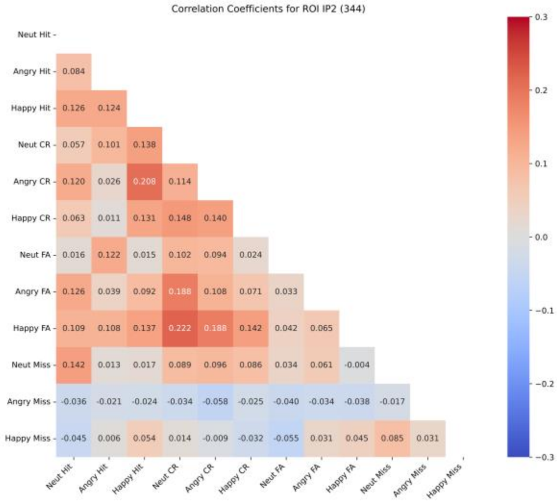
Correlation Coefficients for ROI IP2 (344)
Neut Hit
Angry Hit 0.084
Happy Hit 0.126 0.124
Neut CR 0.057 0.101 0.138
Angry CR 0.120 0.026 0.208 0.114
Happy CR 0.063 0.011 0.131 0.148 0.140
Neut FA 0.016 0.122 0.015 0.102 0.094 0.024
Angry FA 0.126 0.039 0.092 0.188 0.108 0.071 0.033
Happy FA 0.109 0.108 0.137 0.222 0.188 0.142 0.042 0.065
Neut Miss 0.142 0.013 0.017 0.089 0.096 0.086 0.034 0.061 -0.004
Angry Miss -0.036 -0.021 -0.024 -0.034 -0.058 -0.025 -0.040 -0.034 -0.038 -0.017
Happy Miss -0.045 0.006 0.054 0.014 -0.009 -0.032 -0.055 0.031 0.045 0.085 0.031
Neut Hit Angry Hit Happy Hit Neut CR Angry CR Happy CR Neut FA Angry FA Happy FA Neut Miss Angry Miss Happy Miss

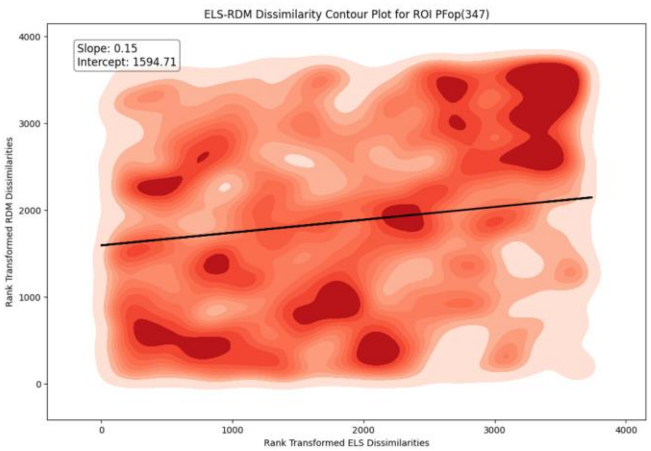
ELS-RDM Dissimilarity Contour Plot for ROI PFop(347)
Slope: 0.15
Intercept: 1594.71
Rank Transformed RDM Dissimilarities
Rank Transformed ELS Dissimilarities
0
1000
2000
3000
4000

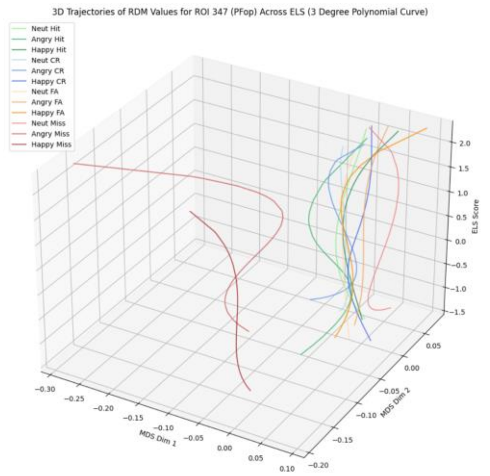
3D Trajectories of RDM Values for ROI 347 (PFop) Across ELS (3 Degree Polynomial Curve)
Neut Hit
Angry Hit
Happy Hit
Neut CR
Angry CR
Happy CR
Neut FA
Angry FA
Happy FA
Neut Miss
Angry Miss
Happy Miss
MDS Dim 1
MDS Dim 2
ELS Score

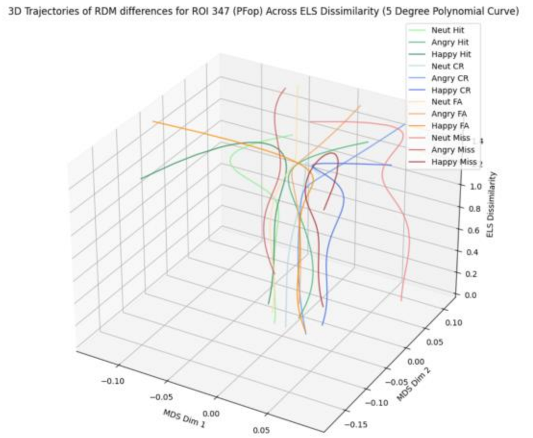
3D Trajectories of RDM differences for ROI 347 (PFop) Across ELS Dissimilarity (5 Degree Polynomial Curve)
Neut Hit
Angry Hit
Happy Hit
Neut CR
Angry CR
Happy CR
Neut FA
Angry FA
Happy FA
Neut Miss
Angry Miss
Happy Miss
MDS Dim 1
MDS Dim 2
ELS Dissimilarity

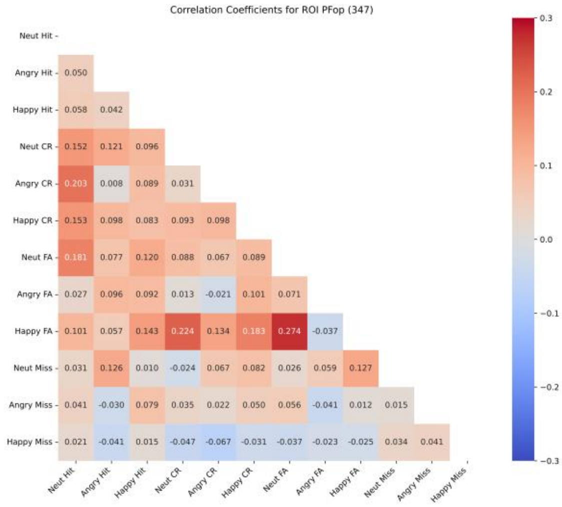
Correlation Coefficients for ROI PFop (347)
Neut Hit
Angry Hit 0.050
Happy Hit 0.058 0.042
Neut CR 0.152 0.121 0.096
Angry CR 0.203 0.008 0.089 0.031
Happy CR 0.153 0.098 0.083 0.093 0.098
Neut FA 0.181 0.077 0.120 0.088 0.067 0.089
Angry FA 0.027 0.096 0.092 0.013 -0.021 0.101 0.071
Happy FA 0.101 0.057 0.143 0.224 0.134 0.183 0.274 -0.037
Neut Miss 0.031 0.126 0.010 -0.024 0.067 0.082 0.026 0.059 0.127
Angry Miss 0.041 -0.030 0.079 0.035 0.022 0.050 0.056 -0.041 0.012 0.015
Happy Miss 0.021 -0.041 0.015 -0.047 -0.067 -0.031 -0.037 -0.023 -0.025 0.034 0.041
Neut Hit Angry Hit Happy Hit Neut CR Angry CR Happy CR Neut FA Angry FA Happy FA Neut Miss Angry Miss Happy Miss

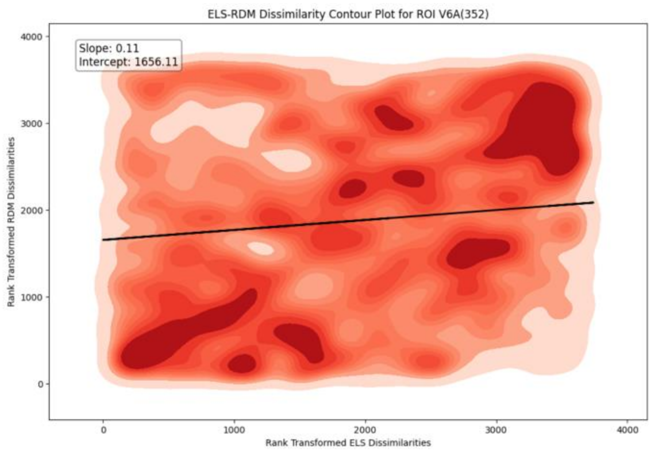
ELS-RDM Dissimilarity Contour Plot for ROI V6A(352)
Slope: 0.11
Intercept: 1656.11
Rank Transformed RDM Dissimilarities
Rank Transformed ELS Dissimilarities
0
1000
2000
3000
4000

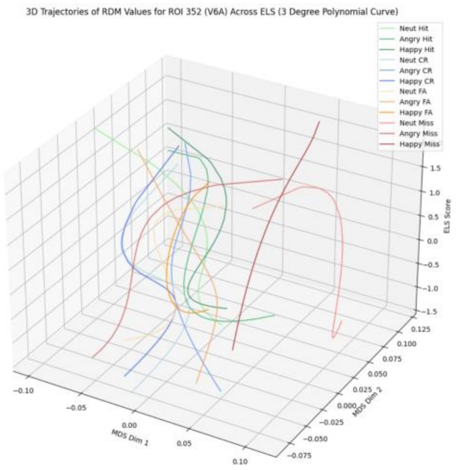
3D Trajectories of RDM Values for ROI 352 (V6A) Across ELS (3 Degree Polynomial Curve)
Neut Hit
Angry Hit
Happy Hit
Neut CR
Angry CR
Happy CR
Neut FA
Angry FA
Happy FA
Neut Miss
Angry Miss
Happy Miss
ELS Score
MDS Dim 1
MDS Dim 2

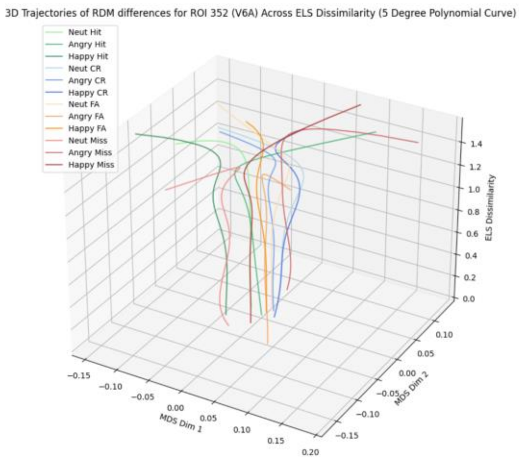
3D Trajectories of RDM differences for ROI 352 (V6A) Across ELS Dissimilarity (5 Degree Polynomial Curve)
Neut Hit
Angry Hit
Happy Hit
Neut CR
Angry CR
Happy CR
Neut FA
Angry FA
Happy FA
Neut Miss
Angry Miss
Happy Miss
ELS Dissimilarity
MDS Dim 1
MDS Dim 2

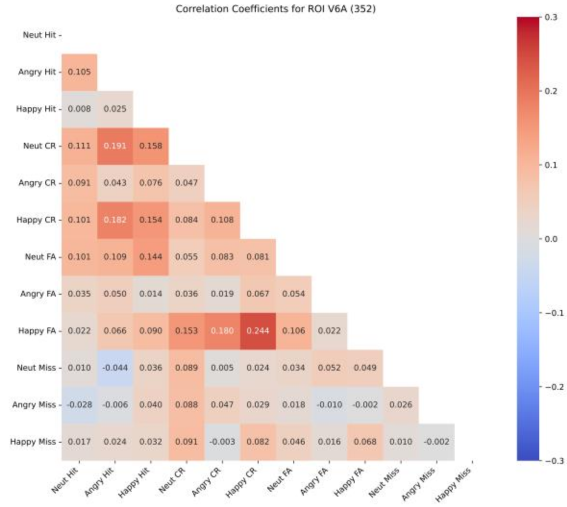
Correlation Coefficients for ROI V6A (352)
Neut Hit
Angry Hit 0.105
Happy Hit 0.008 0.025
Neut CR 0.111 0.191 0.158
Angry CR 0.091 0.043 0.076 0.047
Happy CR 0.101 0.182 0.154 0.084 0.108
Neut FA 0.101 0.109 0.144 0.055 0.083 0.081
Angry FA 0.035 0.050 0.014 0.036 0.019 0.067 0.054
Happy FA 0.022 0.066 0.090 0.153 0.180 0.244 0.106 0.022
Neut Miss 0.010 -0.044 0.036 0.089 0.005 0.024 0.034 0.052 0.049
Angry Miss -0.028 -0.006 0.040 0.088 0.047 0.029 0.018 -0.010 -0.002 0.026
Happy Miss 0.017 0.024 0.032 0.091 -0.003 0.082 0.046 0.016 0.068 0.010 -0.002
Neut Hit Angry Hit Happy Hit Neut CR Angry CR Happy CR Neut FA Angry FA Happy FA Neut Miss Angry Miss Happy Miss
0.3
0.2
0.1
0.0
−0.1
−0.2
−0.3

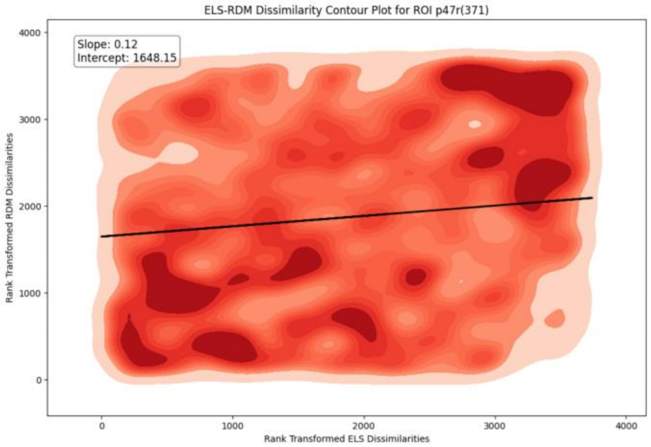
ELS-RDM Dissimilarity Contour Plot for ROI p47r(371)
Slope: 0.12
Intercept: 1648.15
Rank Transformed RDM Dissimilarities
Rank Transformed ELS Dissimilarities

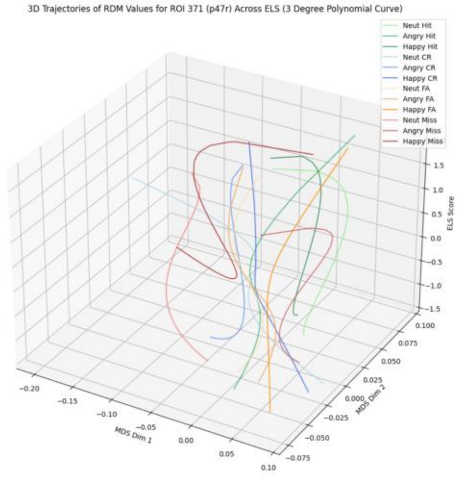
3D Trajectories of RDM Values for ROI 371 (p47r) Across ELS (3 Degree Polynomial Curve)
Neut Hit
Angry Hit
Happy Hit
Neut CR
Angry CR
Happy CR
Neut FA
Angry FA
Happy FA
Neut Miss
Angry Miss
Happy Miss
ELS Score
MDS Dim 1
MDS Dim 2

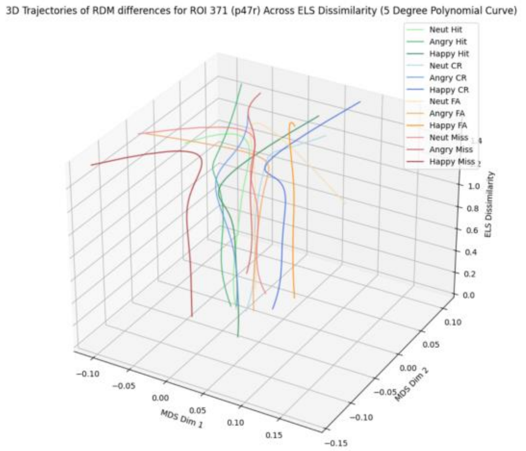
3D Trajectories of RDM differences for ROI 371 (p47r) Across ELS Dissimilarity (5 Degree Polynomial Curve)
Neut Hit
Angry Hit
Happy Hit
Neut CR
Angry CR
Happy CR
Neut FA
Angry FA
Happy FA
Neut Miss
Angry Miss
Happy Miss
ELS Dissimilarity
MDS Dim 1
MDS Dim 2

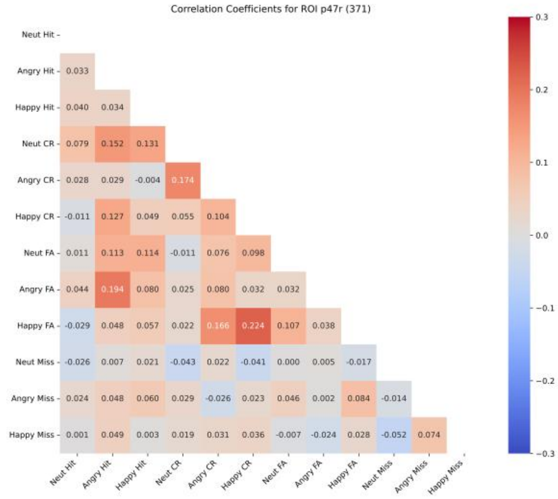
Correlation Coefficients for ROI p47r (371)
Neut Hit
Angry Hit 0.033
Happy Hit 0.040 0.034
Neut CR 0.079 0.152 0.131
Angry CR 0.028 0.029 -0.004 0.174
Happy CR -0.011 0.127 0.049 0.055 0.104
Neut FA 0.011 0.113 0.114 -0.011 0.076 0.098
Angry FA 0.044 0.194 0.080 0.025 0.080 0.032 0.032
Happy FA -0.029 0.048 0.057 0.022 0.166 0.224 0.107 0.038
Neut Miss -0.026 0.007 0.021 -0.043 0.022 -0.041 0.000 0.005 -0.017
Angry Miss 0.024 0.048 0.060 0.029 -0.026 0.023 0.046 0.002 0.084 -0.014
Happy Miss 0.001 0.049 0.003 0.019 0.031 0.036 -0.007 -0.024 0.028 -0.052 0.074
Neut Hit Angry Hit Happy Hit Neut CR Angry CR Happy CR Neut FA Angry FA Happy FA Neut Miss Angry Miss Happy Miss

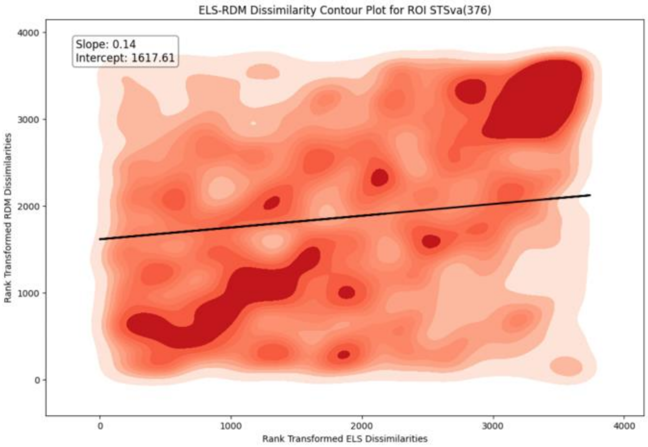
ELS-RDM Dissimilarity Contour Plot for ROI STSva(376)
Slope: 0.14
Intercept: 1617.61
Rank Transformed RDM Dissimilarities
Rank Transformed ELS Dissimilarities

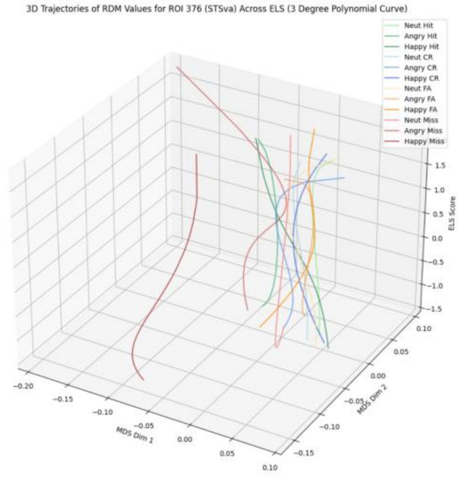
3D Trajectories of RDM Values for ROI 376 (STSva) Across ELS (3 Degree Polynomial Curve)
Neut Hit
Angry Hit
Happy Hit
Neut CR
Angry CR
Happy CR
Neut FA
Angry FA
Happy FA
Neut Miss
Angry Miss
Happy Miss
MDS Dim 1
MDS Dim 2
ELS Score

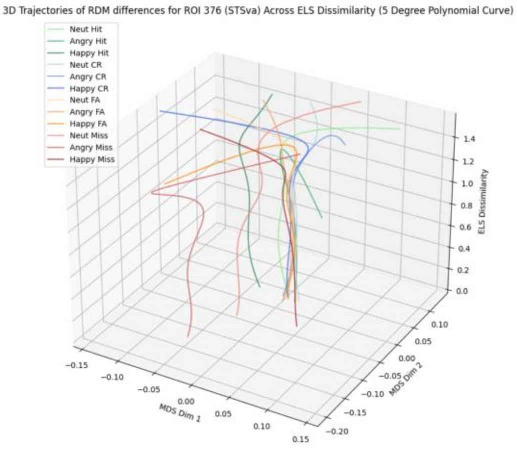
3D Trajectories of RDM differences for ROI 376 (STSva) Across ELS Dissimilarity (5 Degree Polynomial Curve)
Neut Hit
Angry Hit
Happy Hit
Neut CR
Angry CR
Happy CR
Neut FA
Angry FA
Happy FA
Neut Miss
Angry Miss
Happy Miss
MDS Dim 1
MDS Dim 2
ELS Dissimilarity

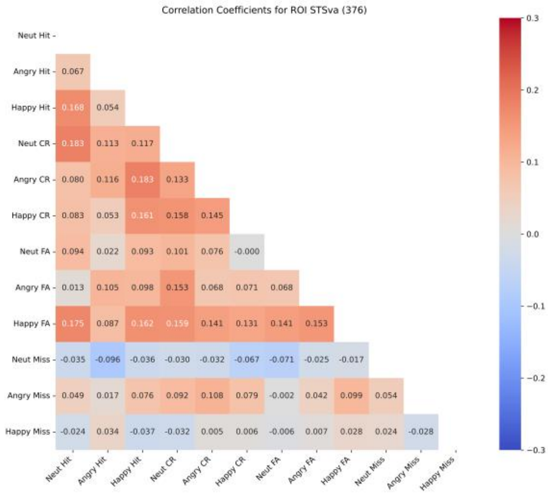
Correlation Coefficients for ROI STSva (376)
Neut Hit
Angry Hit 0.067
Happy Hit 0.168 0.054
Neut CR 0.183 0.113 0.117
Angry CR 0.080 0.116 0.183 0.133
Happy CR 0.083 0.053 0.161 0.158 0.145
Neut FA 0.094 0.022 0.093 0.101 0.076 -0.000
Angry FA 0.013 0.105 0.098 0.153 0.068 0.071 0.068
Happy FA 0.175 0.087 0.162 0.159 0.141 0.131 0.141 0.153
Neut Miss -0.035 -0.096 -0.036 -0.030 -0.032 -0.067 -0.071 -0.025 -0.017
Angry Miss 0.049 0.017 0.076 0.092 0.108 0.079 -0.002 0.042 0.099 0.054
Happy Miss -0.024 0.034 -0.037 -0.032 0.005 0.006 -0.006 0.007 0.028 0.024 -0.028
Neut Hit Angry Hit Happy Hit Neut CR Angry CR Happy CR Neut FA Angry FA Happy FA Neut Miss Angry Miss Happy Miss
