## Supplemental Information for "Similarity in Early Life Stress Exposure is Associated with Similarity in Neural Representations in Early Adulthood"

### *Supplementary figure legends*

**Supplementary Figure 1**: Sorted correlation coefficients for intersubject representational similarity analysis correlations between different ELS measurements and the 360 ROI specific RDMs. Correlations have been sorted within each ELS measurement to demonstrate the distribution and ratio of negative and positive correlations. Correlations were attained from partial Spearman's correlation between the vectorized intersubject dissimilarity matrices constructed from various ELS measurements (Prospective and Retrospective ELS, and the 7 different domains of the Prospective ELS), and the 360 intersubject dissimilarity matrices constructed from pairwise correlation distances between vectorized representational dissimilarity matrices for each cortical region of the Human Connectome Project Multimodal Parcellation 1.0.

**Supplementary Figure 2**: Cortical surface projections of the within-subjects main effects (1000 permutations, $p = .005$ cluster p threshold; $p = .001$ cluster forming p threshold) on inflated left and right hemisphere surfaces. Sulci are highlighted in dark grey, gyri in light grey. Displayed are main effects of correctness (correct vs. incorrect trials; Fmin = 40, Fmax = 100), motor response (response vs. nonresponse trials; Fmin = 15, Fmax = 50), valence (happy vs. angry vs. neutral facial expression stimuli trials; Fmin = 9, Fmax = 20), and interactions for correctness × motor response (Fmin = 9, Fmax = 70) and valence × motor response (Fmin = 9; Fmax = 20).

**Supplementary Figures 3-41**: Four visualizations computed for each region with significant Prospective ELS ISDM × ROI ISDM correlations. The contour plot (top left) displays the association between the rank transformed pairwise distances of the region's RDMs and the rank transformed Prospective ELS pairwise distances, highlighting their positive association. The 3-dimensional MDS plot (top right) on the raw values of the region's RDMs shows how the 2-dimensional representational structure, or space, extracted from the RDM changes as a function of increasing Prospective ELS. Here, a 3-degree polynomial curve was used for smoothing the lines. The 3-dimensional MDS plot (bottom left) on the difference RDM values of the region's RDMs against Prospective ELS dissimilarities displays how the differences between each participant pair's difference RDMs evolve as a function of Prospective ELS dissimilarity. Difference RDMs were calculated as absolute elementwise differences between the pair's RDMs. The plot highlights how pairwise differences in Prospective ELS are associated with relatively stable absolute differences in most of the RDM's elements, except for large dissimilarities in ELS which are associated with large deviations in how the RDM's elements change as a function of Prospective ELS dissimilarity. The heatmap (bottom right) displays Pearson's correlations between the pairwise differences in each RDM element and pairwise differences in Prospective ELS for the region, highlighting the direction and consistency of change in each element as a function of Prospective ELS.

### *Titles for supplementary figures 3-41*

**Supplementary Figure3**: lh_26_SFL

**Supplementary Figure4**: lh_51_1

**Supplementary Figure5**: lh_54_6d

**Supplementary Figure6**: lh_60_p32pr

**Supplementary Figure7**: lh_94_47

**Supplementary Figure8**: lh_107_TA2

**Supplementary Figure 9**: lh_111_AVI

**Supplementary Figure 10**: lh_112_AAIC

**Supplementary Figure 11**: lh_128_STSva

**Supplementary Figure 12**: lh_153_VMV1

**Supplementary Figure 13**: lh_160_VMV2

**Supplementary Figure 14**: lh_162_31a

**Supplementary Figure 15**: lh_179_a32pr

**Supplementary Figure 16**: rh_211_PEF

**Supplementary Figure 17**: rh_215_POS2

**Supplementary Figure 18**: rh_247_7PC

**Supplementary Figure 19**: rh_248_LIPv

**Supplementary Figure 20**: rh_252_2

**Supplementary Figure 21**: rh_256_6v

**Supplementary Figure 22**: rh_259_a24pr

**Supplementary Figure 23**: rh_268_8Ad

**Supplementary Figure 24**: rh_293_OFC

**Supplementary Figure 25**: rh_295_LIPd

**Supplementary Figure 26**: rh_299_43

**Supplementary Figure 27**: rh_300_OP4

**Supplementary Figure 28**: rh_306_PoI2

**Supplementary Figure 29**: rh_307_TA2

**Supplementary Figure 30**: rh_308_FOP4

**Supplementary Figure 31**: rh_309_MI

**Supplementary Figure 32**: rh_311_AVI

**Supplementary Figure 33**: rh_314_FOP3

**Supplementary Figure 34**: rh_316_PFt

**Supplementary Figure 35**: rh_317_AIP

**Supplementary Figure 36**: rh_332_TE1a

**Supplementary Figure 37**: rh_344_IP2

**Supplementary Figure 38**: rh_347_PFop

**Supplementary Figure 39**: rh_352_V6A

**Supplementary Figure 40**: rh_371_p47r

**Supplementary Figure 41**: rh_376_STSva
